# Multimodal integration of single cell ATAC-seq data enables highly accurate delineation of clinically relevant tumor cell subpopulations

**DOI:** 10.1101/2024.10.11.617736

**Authors:** Kewei Xiong, Ruofan Ding, Yangmei Qin, Xudong Zou, Jiguang Wang, Chen Yu, Lei Li

## Abstract

Accurately deciphering tumor cellular heterogeneity is crucial for developing effective treatments. While single-cell epigenomics assays offer powerful insights into studying tumor heterogeneity beyond the transcriptional level, their application remains a significant challenge. To address this, we introduce Multimodal-based Analysis of scATAC-Seq data (MAAS), a method designed to identify functional tumor cell subpopulations and reconstruct their lineages from single-cell sequencing assay for transposase-accessible chromatin (scATAC-seq). MAAS uniquely integrates multimodal information, encompassing chromatin accessibility, copy number variations (CNVs), and single-nucleotide variants, through self-expressive multimodal matrix factorization. MAAS outperforms existing methods in accuracy and robustness on both simulated and real datasets. When applied to multiple glioma scATAC-seq datasets, MAAS uncovered a previously hidden tumor cell subpopulation resistant to temozolomide treatment. Furthermore, MAAS effectively detected clinically relevant subpopulations with low CNVs, such as those found in pediatric ependymoma and B-cell lymphoma. Additionally, we developed a novel MAAS-derived clinical signature that provides superior prognostic prediction than traditional clinicopathologic features across multiple cancer types. In summary, MAAS significantly advances the identification of clinically relevant tumor cell subpopulations, thereby accelerating the discovery of potential therapeutic targets.

## Introduction

Cancer cells undergo various genetic and epigenetic changes that drive the formation of distinct subpopulations during tumor progression^1, 2^. Identifying these critical subpopulations accurately is essential for developing effective treatments^2^. Single-cell sequencing technologies have revolutionized our understanding of tumor cellular composition compared to traditional bulk analyses. For instance, single-cell RNA sequencing (scRNA-seq) is a widely used approach for profiling clinical phenotype-associated subpopulations^3-5^. Although scRNA-seq can distinguish malignant from non-malignant subpopulations and examine cell heterogeneity by analyzing expression-derived genetic mutations such as copy number variations^6, 7^, it often fails to identify the clinically relevant subpopulations that are not solely linked to gene expression^8^. This is mainly because scRNA-seq primarily captures transcriptional activity and often misses critical epigenetic and regulatory changes^9, 10^.

Single-cell epigenomics assays, such as single-cell assay for transposase-accessible chromatin using sequencing (scATAC-seq), enable robust profiling of cell subpopulations beyond gene expression and can reveal distinct regulatory information that controls gene expression. However, applying scATAC-seq to study tumor subpopulations has been computationally challenging. Traditional methods mainly rely on CNVs^11-13^, such as copy-scAT and epiAneufinder, to determine the genetic heterogeneity of tumor cells. Despite these advances, CNV analysis alone often fails to detect clinically relevant tumor subpopulations. For example, melanoma subpopulations with varying anti-PD-1 responses are characterized more by their distinct mutation profiles than by CNV events^7^. Furthermore, the interplay between genetic and epigenetic changes enables subpopulations to circumvent therapeutic barriers, leading to cancer progression^14, 15^, thus highlighting the need to integrate these features. Unfortunately, existing methods fail to fully utilize the spectrum of epigenetic and genetic information available from scATAC-seq data. These methods are also limited in delineating tumor cell subpopulations with low CNV heterogeneity, such as those found in hematopoietic and pediatric cancers^16^. Additionally, scATAC-seq data often suffers from inherent high sparsity and technical noise, posing significant challenges in determining informative features for dissecting tumor cell subpopulations^17^.

Here, we have developed a novel computational method called Multimodal-based Analysis of scATAC-Seq data (MAAS) that accurately identifies tumor cell subpopulations and infers their evolutionary lineages by leveraging and integrating informative multimodal features such as CNVs, single-nucleotide variants (SNVs), and chromatin accessibility data. Furthermore, MAAS outperformed other state-of-the-art methods on both simulated and real datasets. When applied to a glioma tumor, MAAS uncovered a previously overlooked subpopulation that was resistant to temozolomide. In pediatric ependymoma with low CNVs, MAAS enables the discovery of a progressive tumor cell subpopulation that was associated with multidrug resistance. We further developed an MAAS-derived multimodal clinical signature by integrating subpopulation-specific transcription factor (TF) activity and expression, which provided a more accurate prognostic prediction than traditional clinicopathologic characteristics across multiple cancer types. In conclusion, MAAS is a reliable and robust tool for identifying clinically relevant tumor cell subpopulations, helping to uncover new disease mechanisms and improve tumor diagnosis and therapeutic strategies.

## Results

### MAAS achieved superior accuracy in predicting tumor cell subpopulations

To delineate cellular heterogeneity in tumors, we developed an algorithm called MAAS to accurately identify tumor cell subpopulations by integrating genetic and epigenetic features derived from scATAC-seq data (Fig. 1a and Supplementary Figs. 1-2; Methods). To ensure our results were not biased by CNVs, which often affect the quantification of chromatin accessibility^12^, MAAS implemented a weighted correction strategy to adjust for this confounding effect. Additionally, since SNVs derived from scATAC-seq data can be highly sparse and noisy, MAAS utilized a modified robust principal component analysis and the inexact augmented Lagrange multiplier algorithm to accurately detect SNV information (Fig. 1a). We then estimated cell similarities using cosine distance for chromatin accessibility and hamming distance for CNV and SNV, and integrated them using multimodal non-negative matrix factorization (Fig. 1b). To further enhance clustering accuracy, MAAS decomposed the cell similarities into a latent variable matrix **W** and diagonal coefficient matrix **H**, and used the correlation matrices **A** as self-expressions for better clustering^18^. To obtain the optimal result, MAAS employed the multiplication update rule to minimize the value of the loss function. Finally, the tumor cells were classified into subpopulations using K-means clustering, and a phylogenetic tree depicting the cell lineages was also constructed (Fig. 1c).

**Fig. 1.**
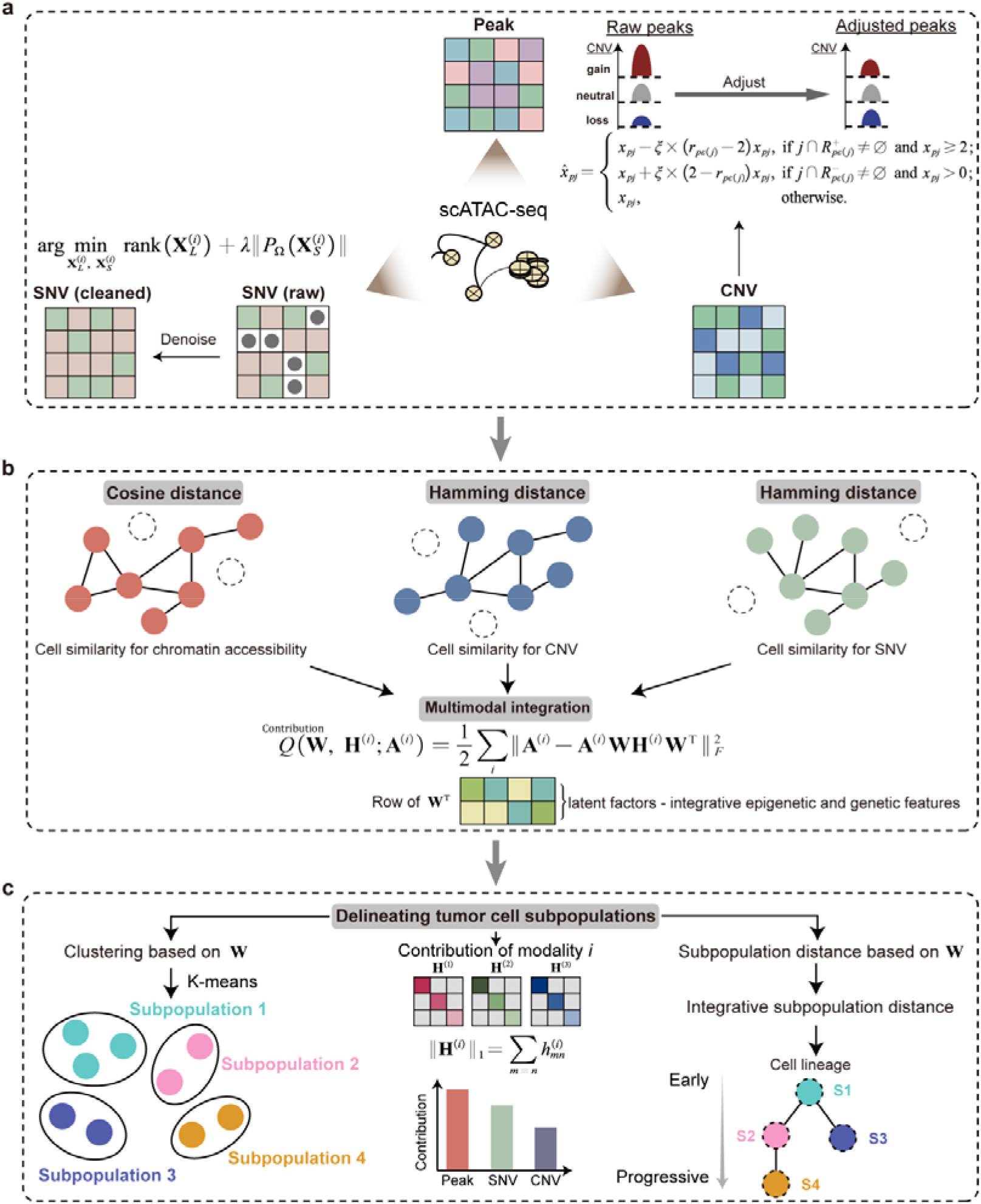
The MAAS workflow. **a**. MAAS takes as input a cell-by-peak matrix, a cell-by-CNV matrix, and a cell-by-SNV matrix. Raw peak data are adjusted based on the copy number values of the corresponding genomic regions. A robust principal component analysis is applied to the SNV data to reduce noise and generate a low-rank matrix. **b**. Cell similarities for each omics layer are calculated using cosine or Hamming distances. These similarities are integrated through a modified matrix factorization strategy, enabling the inference of a latent space that captures both genetic and epigenetic features through iterative updates. **c**. Tumor cell subpopulations are identified using the latent factors. The contribution of each modality to the subpopulation is determined by the first-order norm of the coefficient matrix **H**. Consensus cell distances are derived by calculating the Euclidean distance from the cell-by-latent factor matrix, which is then used to reconstruct cell lineage represented by a minimum evolution phylogeny.

To systematically evaluate the performance of MAAS, we conducted a simulation analysis to assess whether MAAS could accurately deconvolute the labeled tumor cell subpopulations. We first generated three simulated cell clusters as ground truth datasets, where clusters 1 and 2 had different genetic features from cluster 3, and clusters 1 and 2 had distinct chromatin accessibility profiles (Fig. 2a and Methods). MAAS successfully separated clusters 1 and 2 indistinguishable by the single-modality approach (Fig. 2a,b). Specifically, MAAS identified 83.3%, 100%, and 98.28% of the three clusters (Fig. 2c). Moreover, MAAS accurately recovered 98.7% of the differentially accessible chromatin regions (DACRs) between cluster 1 and cluster 2 (Supplementary Fig. 3). Compared with four simulated cell clusters, MAAS accurately identified 97.8% and 97% of cells in clusters 1 and 2, respectively, and correctly distinguished all cells within clusters 3 and 4 (Supplementary Fig. 4a-c).

**Fig. 2.**
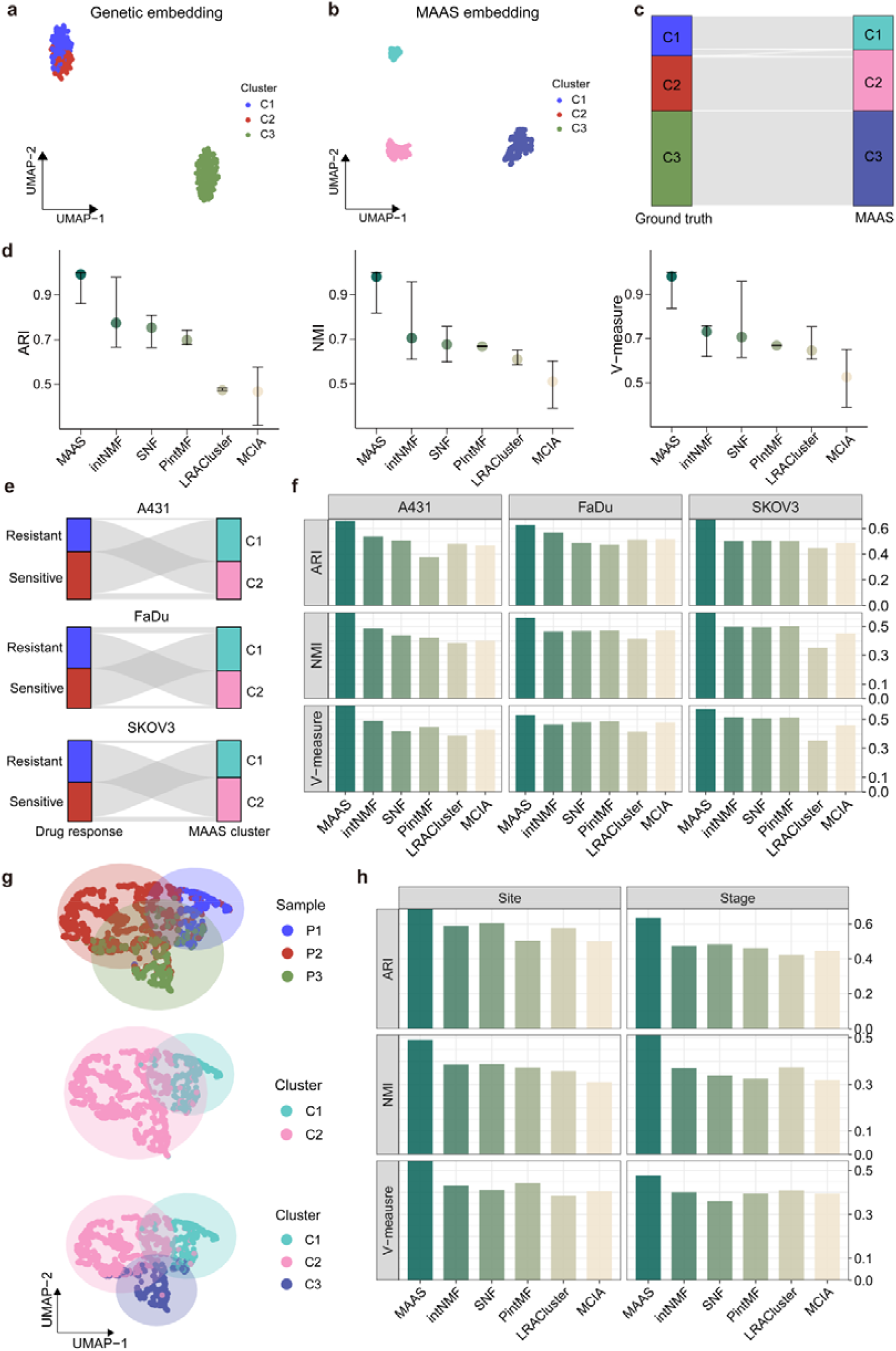
Benchmarking analysis of tumor subpopulation identification. **a**. UMAP embedding based on genetic features of three subpopulations. **b**. UMAP embedding based on MAAS latent factors for the three identified subpopulations. **c**. Consistency of cell distribution across the three subpopulations when comparing ground-truth to MAAS results. **d**. Consistency of cell distributions across four subpopulations between the ground-truth and MAAS results. **e**. Sankey plot illustrating the distribution of drug-sensitive and drug-resistant subpopulations compared with MAAS clusters across three different cell lines. **f**. Accuracy of classifying drug-sensitive and drug-resistant subpopulations by different methods, as measured by ARI (top), NMI (middle), and V-measure (bottom). **g**. UMAP embedding showing three ovarian cancer samples with different clinical features: P1 (metastasis and advanced stage), P2 (primary and early stage), and P3 (primary and advanced stage). **h**. Accuracy of classifying subpopulations based on pathological stages, as well as distinguishing between primary and metastatistic tumor cells, as measured by ARI (top), NMI (middle), and V-measure (bottom) across different methods.

To further evaluate the robustness of the MAAS method, we compared it with other multi-omics integration and clustering tools, including uniform manifold approximation and projection (UMAP)^19^, intNMF^20^, PintNMF^21^, SNF^22^, LRACluster^23^, and MCIA^24^ (Methods). We began by randomizing the tumor cell clusters and evaluated the performance based on three metrics: the adjusted Rand index (ARI), normalized mutual information (NMI), and V-measure (Methods). MAAS significantly outperformed UMAP multimodal clustering (Supplementary Fig. 4d) and other integration methods with the median ARI, NMI, and V-measure values of 0.992, 0.981, and 0.981, respectively (Fig. 2d). Additionally, we varied the number of cells in the tumor cell clusters for each simulation and found MAAS still consistently achieved the highest ARI, NMI, and V-measure scores (Supplementary Fig. 4e). Further examining the impact of the number of clusters on clustering performance demonstrated that MAAS exhibited the best performance across a diverse range of clusters (Supplementary Fig. 4f). We also evaluated the clustering performance across different levels of data sparsity, ranging from 10% to 90%, and found MAAS consistently showed superior performance, with over 50% classification even at the 90% of data sparsity (Supplementary Fig. 4g).

Moreover, we compared the performance of MAAS with CNV estimates derived from single-cell whole-genome sequencing (scWGS) data for gastric cancer (SNU601 cell line)^12^. MAAS accurately recovered the four tumor cell clusters characterized by the amplification of chromosomes 1 and 3 and the deletion of chromosomes 4 and 18, with an average Pearson’s correlation of 0.885 (Supplementary Fig. 5 and Supplementary Data 1). To further access the performance of MAAS in identifying clinically pertinent tumor cell subpopulations, we applied it to three cancer cell lines^25^: epidermoid carcinoma A431, hypopharyngeal cancer FaDu, and ovarian cancer (OC) SKOV3. These cell lines contain experimentally validated drug-sensitive and resistant subpopulations. MAAS accurately distinguished between drug-sensitive and resistant tumor cells with the mean AMI of 0.652, NMI of 0.551, and V-measure of 0.564, significantly outperforming other available methods (Fig. 2e,f and Supplementary Fig. 6). Additionally, in an OC scATAC-seq dataset collected from three tumors^26^, the MAAS method effectively identified metastatic tumor cells from primary ones and accurately distinguished early- and advanced-stage tumor cells (Fig. 2g,h). Overall, MAAS demonstrated superior performance in predicting tumor cell subpopulations compared to state-of-the-art methods.

### MAAS delineated a new glioma cell subpopulation with temozolomide resistance

Glioblastoma is the most common and aggressive primary brain malignancy in adults^27^. To explore the heterogeneity of glioblastoma, we applied MAAS to a scATAC-seq dataset of adult glioblastoma^11^ (GBM) containing 965 tumor cells (Supplementary Figs. 7-8). MAAS not only successfully identified the clusters recognized by the traditional single modality approaches, but also revealed a new cluster (cluster 1) (Fig. 3a,b and Supplementary Figs. 9-10). A function enrichment analysis of this new cluster showed significantly enriched in apoptosis, angiogenesis, and KRAS signaling pathways (Fig. 3c). To trace cell lineages, we constructed a MAAS-based phylogenetic tree to map the tumor development, revealing a stronger phylogenetic signal (average *K*-statistic = 1.62) than CNVs (average *K*-statistic = 1.11) and chromatin accessibility (average *K*-statistic = 1.07) (Fig. 3d,e and Supplementary Data 2; Methods). In cluster 1, we identified six SNVs (Fig. 3e and Supplementary Fig. 11) and 13,034 DACRs specific to this cluster (Fig. 3f and Supplementary Data 3). Furthermore, all six driven mutations were located within the cluster 1-activated DACRs (Fig. 3g and Supplementary Fig. 12), and these findings were cross-validated using an independent pediatric GBM (pGBM) dataset^11^ (Supplementary Fig. 13 and Supplementary Methods).

**Fig. 3.**
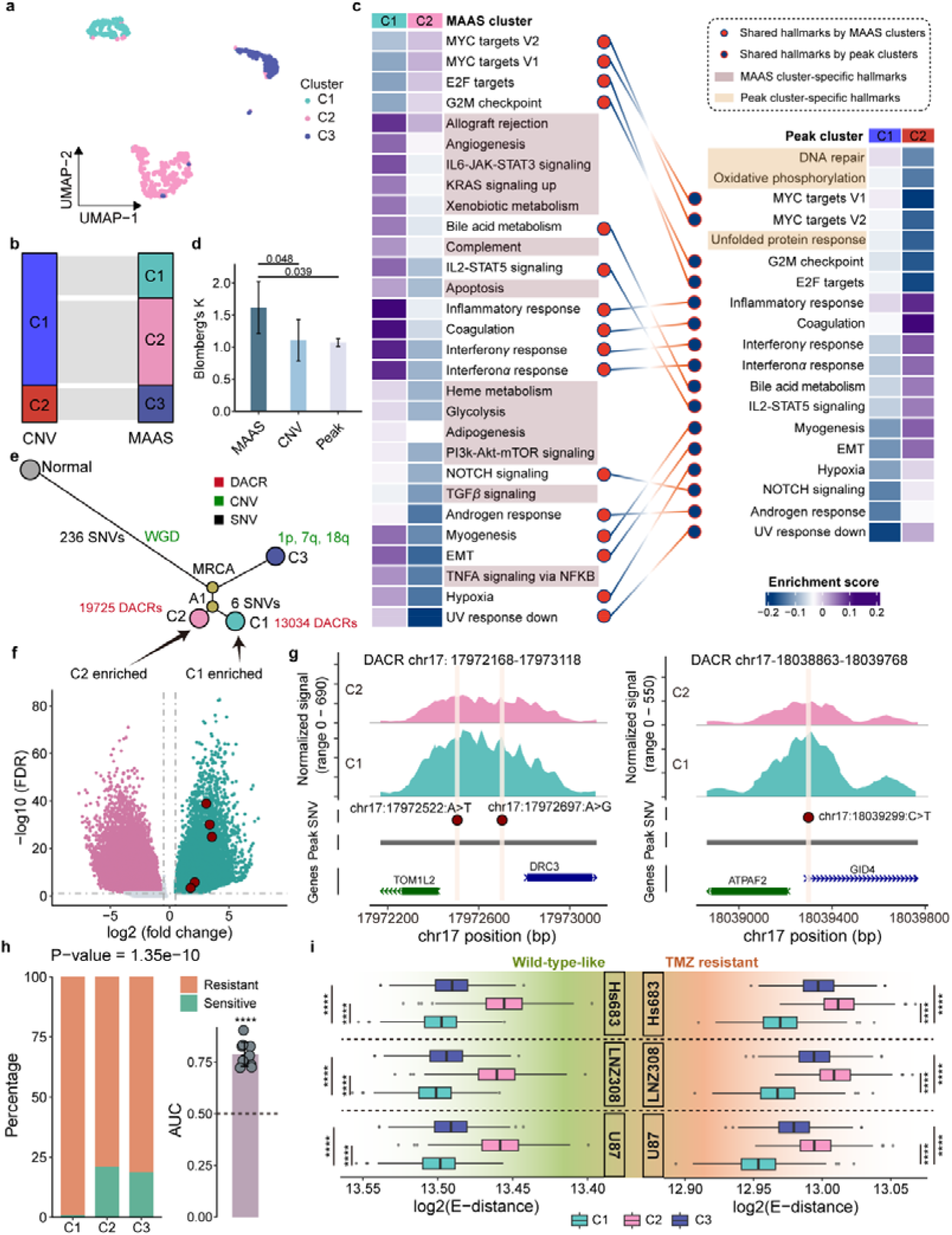
A new glioma subpopulation with drug resistance. **a**. UMAP embedding of the three tumor cell clusters determined by MAAS. **b**. Sankey plot showed the correspondence between clusters identified by CNVs and those identified by MAAS. **c**. Differentially enriched cancer hallmark signatures between clusters. The left panel displays pathways enriched in MAAS clusters, while the right panel shows pathways enriched in tumor cell clusters identified by chromatin accessibility. The heatmap colors represent enrichment scores, with color shades indicating unique hallmarks detected by either MAAS or chromatin accessibility. **d**. Blomberg’s K statistic comparisons of each feature of MAAS, chromatin accessibility and CNVs. The P-value was determined by a Welch t-test. **e**. Phylogenetic tree of MAAS-identified clusters, depicting the timing of CNVs, SNVs, and DACRs relative to the most recent common ancestor (MRCA). The tree is reconstructed using a minimum evolution model, with annotations for the common ancestor (A1) and whole-genome doubling (WGD) events. **f**. Volcano plot showing DACRs (FDR < 0.05 and |logFC| > 0.5) between MAAS-identified clusters 1 and 2. Regions containing the six driver mutations specific to cluster 1 are highlighted. **g**. scATAC-seq peak tracks for accessible regions in clusters 1 and 2, with noncoding SNVs marked by dark red dots. **h**. Distribution of predicted temozolomide (TMZ)-sensitive and resistant cells across MAAS-identified clusters. The P-value, calculated by a chi-square test, is shown, along with the accuracy of predictions measured by the area under the curve (AUC). The dotted line represents the baseline of 0.5. Asterisks indicate significance based on permutations. **i**. Energy distance (E-distance) between the three MAAS clusters and TMZ-sensitive and resistant cells across three cell lines. The center line of the boxplot indicates the median, the box limits show the first and third quartiles, and the whiskers extend to the maximum and minimum values within 1.5 times the interquartile range from the hinge. The P-value was determined by a two-tailed Wilcoxon rank-sum test; **** indicates P-value < 0.0001.

To further characterize the clinical relevance of MAAS-identified clusters, we first linked the gene activities of each cluster with the half-maximal inhibitory concentration (IC_50_) of first-line glioma chemotherapeutic, temozolomide (TMZ) (Methods). Notably, cluster 1 exhibited significantly greater resistance to TMZ (Fisher’s exact test, P-value = 1.35×10^−10^), with accurate predictions reflected in the area under the curve (AUC) values of 0.788, 0.779, and 0.701 (Fig. 3h and Supplementary Fig. 14a-c). We also calculated the energy distance (E-distance)^28^ between MAAS clusters and TMZ-associated subpopulations from six glioma cell lines^29, 30^ (Methods). Cluster 1 consistently showed the shortest E-distance to TMZ-resistant subpopulations (Fig. 3i and Supplementary Fig. 14d; P-value < 0.0001). The new cluster 1 subpopulation, characterized by temozolomide resistance, was further validated using another independent GBM dataset (Supplementary Figs. 15-16). In summary, MAAS identified a novel glioma cell subpopulation with strong TMZ resistance, underscoring the potential of the MAAS method as a powerful tool for accurately classifying new clinically relevant glioma cell subpopulations.

### MAAS detects clinically relevant tumor cell subpopulations with low CNVs

Many tumors, such as pediatric ependymoma, exhibit a low frequency of CNV events, often less than 10%^31^, posing a great challenge for traditional methods to resolve tumor heterogeneity. To demonstrate the utility of the MAAS method in detecting subpopulations with low CNVs, we applied it to a scATAC-seq dataset of pediatric posterior fossa ependymoma (PPFE)^32^. Our analysis revealed that MAAS accurately predicted the two major tumor cell subpopulations from 1,689 tumor cells (Supplementary Figs. 17-18), providing a clearer distinction between tumor cell clusters than the traditional methods (Fig. 4a-d) (Supplementary Figs. 19-20). Functional enrichment analysis of the MAAS-predicted clusters showed that the MAAS not only recovered the traditional hallmark pathway but also identified an additional subset of cancer-related pathways enriched specifically in cluster 2, including *MYC* target, *NOTCH* signaling, *P53* pathway, unfolded protein response, and apoptosis (Fig. 4b). This suggests cluster 2, driven primarily by chromatin accessibility and SNVs, represents a new functional subpopulation detected by MAAS (Fig. 4c,d). Next, we used Monocle^33^ to infer the developmental trajectories of cells and observed a stepwise transition from cluster 1 to cluster 2 (Fig. 4e and Supplementary Fig. 21; Supplementary Methods). To further validate the dynamic changes between MAAS-predicted clusters, we generated a phylogenetic tree that demonstrated significantly superior performance, with an average *K*-statistic of 1.37, compared to single-modality methods in depicting evolutionary lineages. Our analysis revealed that cluster 2 had a higher mutation burden and chromatin accessibility than cluster 1 (Fig. 4f,g; Supplementary Data 4). We also evaluated the proliferative and migration characteristics of each cluster, using activities of proliferation signature genes *MKI67, PCNA, IGF1, ITGB2, PDGFC, JAG1, PHGDH, BCL2*^34, 35^ and migration-related genes *ARID5B* and *FAT1*^36, 37^ (Supplementary Methods). Cluster 2 exhibited significantly higher proliferation and migration scores than cluster 1 (Fig. 4g; Wilcoxon rank-sum test, P-value = 0.015 and 0.002 respectively), indicating that cluster 2 has significant metastatic potential and is highly progressive.

**Fig. 4.**
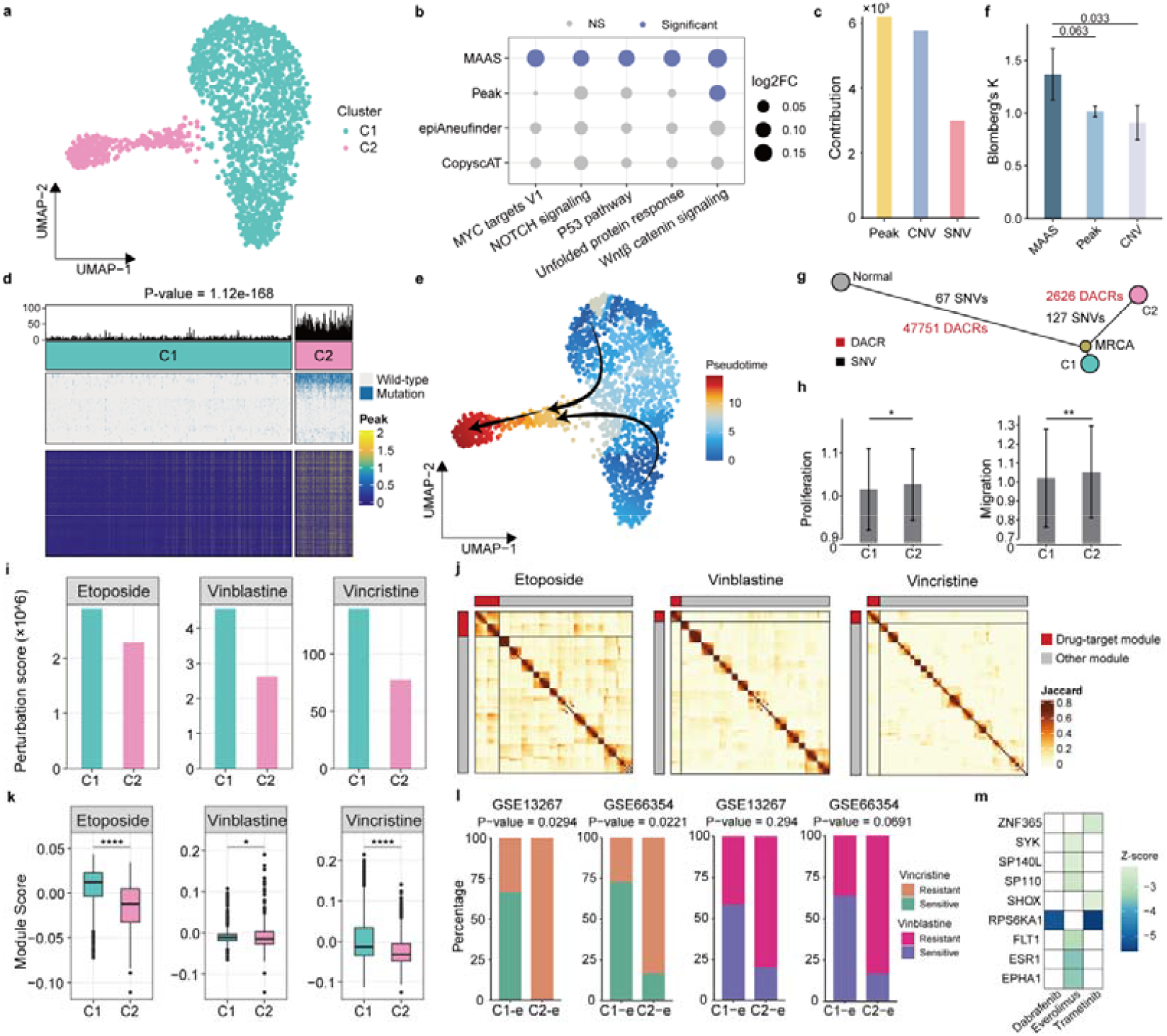
A new pediatric ependymoma cell subpopulation with low CNVs associated with multidrug resistance. **a**. UMAP embedding of the two tumor cell subpopulations identified by MAAS. **b**. Significantly enriched cancer hallmarks in MAAS-identified cluster 2, with thresholds of logFC > 0.1 and false discovery rate (FDR) < 0.05. **c**. Contribution of each modality to the identification of subpopulations. **d**. Distribution of SNVs and top 1,000 DACRs between clusters 1 and 2. The P-value for differences in mutational frequency was determined by the Kruskal-Wallis test. **e**. Pseudo-time ordering of tumor cell subpopulations showing their developmental trajectories. **f**. Blomberg’s K statistic comparisons of each feature of MAAS, chromatin accessibility and CNVs. The P-value was determined by a Welch t-test. **g**. Phylogenetic tree of MAAS-identified clusters, depicting the timing of SNVs and DACRs relative to the most recent common ancestry (MRCA). This tree is reconstructed using a minimum evolution model. **h**. Proliferation and migration scores for clusters 1 and 2 with error bars representing ± standard deviations. P-values were determined by the Wilcoxon rank-sum test. **i**. Perturbation scores for three first-line drugs where a lower score indicates a more resistant phenotype. **j**. Gene modules constructed based on co-expression networks of drug targets using non-negative matrix factorization. A low module score indicates a lower drug response rate. **k**. Box plots showing drug-target module scores between clusters 1 and 2. The center line represents the median, and the lower and upper hinges represent the first and third quartiles. The whiskers extend to the maximum and minimum values within 1.5 times the interquartile range from the hinge. P-values were determined by the Wilcoxon rank-sum test. **l**. Distribution of predicted drug-sensitive and resistant cells between MAAS-determined clusters across different bulk RNA-seq references. P-value were calculated using Fisher’s exact test. **m**. Heatmap showing the reduction in expression levels of upregulated transcription factors and kinases in cluster 1 following approved target therapy.

We also investigated the response of new cluster 2 to several first-line chemotherapeutics, including etoposide, vinblastine and vincristine. First, we estimated the drug-resistant subpopulation using scRank^38^ (Methods) and observed that new cluster 2 exhibited lower perturbation scores compared to cluster 1 for each of the three drugs (Fig. 4h), suggesting that cluster 2 is more drug-tolerant than cluster 1. Additionally, we noted that cluster 2 had significantly lower scores for the drug-target gene modules of the three drugs as determined by the activity of the drug-target gene module (Fig. 4i,j and Supplementary Data 5; Wilcoxon rank-sum test, P-value = 8.38 × 10^−49^, 0.015 and 7.68 × 10^−16^ respectively). We then performed deconvolution analysis for pediatric ependymoma patients (Supplementary Methods) and found that drug-resistant samples contained an average of 20.93% more cluster 2 cells (Fig. 4k). Further, we used data from LINCS consortium^39^ to screen for potential target therapy regimens against cluster 2-specific transcription factors and kinases (Supplementary Data 6). We found three FDA-approved antineoplastic drugs, including dabrafenib, everolimus, and trametinib, that significantly decreased the expression level of *EPHA1, ESR1, FLT1, RPS6KA1, SHOX, SP110, SP140L, SYK* and *ZNF365* (Fig. 4l). We then performed a perturbation analysis on *ESR1*, as it was the most down-regulated transcription factor following everolimus treatment (Supplementary Fig. 22a and Supplementary Data 7; Supplementary Methods). Interestingly, we found that inhibiting *ESR1* significantly reversed the original tumor progression trajectory from cluster 1 to cluster 2 (Supplementary Fig. 27b,c), indicating that everolimus could effectively suppress the proliferation and migration of cells from cluster 2.

Moreover, we applied MAAS to a 10x multiome dataset of B-cell lymphoma, which also exhibits low CNVs^40, 41^. This dataset includes scRNA-seq and scATAC-seq data measured simultaneously for each cell. Our analysis revealed that MAAS more effectively segregated the 2,077 tumor cells into three distinct clusters compared to traditional single-modality methods, such as inferCNV^42^ and CopyKAT^6^, which are widely used to identify tumor cell subpopulations using gene expression-derived copy numbers (Supplementary Figs. 23-24 and 25a,b). Notably, although these tumor cell clusters exhibited low CNVs, they displayed largely distinct SNV and chromatin accessibility profiles (Supplementary Fig. 25c,d and Supplementary Data 8; Wilcoxon rank-sum test, P-value = 4.29×10^−134^). We also found that while 3,077 genes had the same transcriptional level across clusters, the activity level of accessible chromatin regions of these genes was increased in cluster 3 (Supplementary Fig. 25e-g), suggesting that changes in the accessible chromatin regions may precede gene expression alterations. We further calculated the E-distance between MAAS clusters with wild-type versus SUMO-activating enzyme inhibitor-treated B-cell lymphoma cell lines, separately, based on their chromatin accessibility profiles (Supplementary Methods). Cluster 3 exhibited the significantly longest E-distance to the drug-treated subpopulation (Supplementary Fig. 25h; student’s t-test, P-value < 0.001), indicating cluster 3 had a highly insensitive drug response. Collectively, our MAAS analyses demonstrate the method’s effectiveness in detecting clinically relevant subpopulations with low CNVs, such as those found in pediatric ependymoma and B-cell lymphoma.

### A new MAAS-derived clinical signature across multiple cancer types

To evaluate the clinical utility of the MAAS method, we developed a new MAAS-derived clinical signature named MAASig, based on subpopulation-specific TF expression and activity inferred from RNA-seq data^43^, indicating the degree of tumor heterogeneity (Fig. 5a; Supplementary Methods). In addition to glioblastoma, ovarian cancer and B-cell lymphoma, we also applied MAASig to hepatocellular carcinoma (HCC) and clear cell renal cell carcinoma (ccRCC) (Supplementary Tables 1-2). Notably, MAASig demonstrated significant prognostic value and superior prediction accuracy across all five cancer types (Fig. 5b,c; log-rank test, P-value = 6.77 × 10^−57^ and 8.44 × 10^−77^ and 7.96 × 10^−240^, 3.11 × 10^−46^ and 1.84 × 10^−22^, respectively), outperforming single modality-based signature and traditional clinicopathologic variables (Supplementary Figs. 26-32). For example, in glioblastoma, MAASig, with an average concordance index (C-index) of 0.775 and a time-dependent AUC of 0.850, significantly outperformed clinical characteristics, including age, IDH mutation status, 1p19q copy numbers, and *MGMT* promoter methylation (Fig. 5b and Supplementary Figs. 31). In ccRCC, MAASig consistently achieved the highest prediction accuracy with an average C-index of 0.770 and a time-dependent AUC of 0.780 (Fig. 5b and Supplementary Figs. 31). Notably, MAASig remained significantly independent of other clinical features in both training and validation sets across the cancer types (Supplementary Figs. 33-37), demonstrating its superiority and robustness in prognosticating patient survival.

**Fig. 5.**
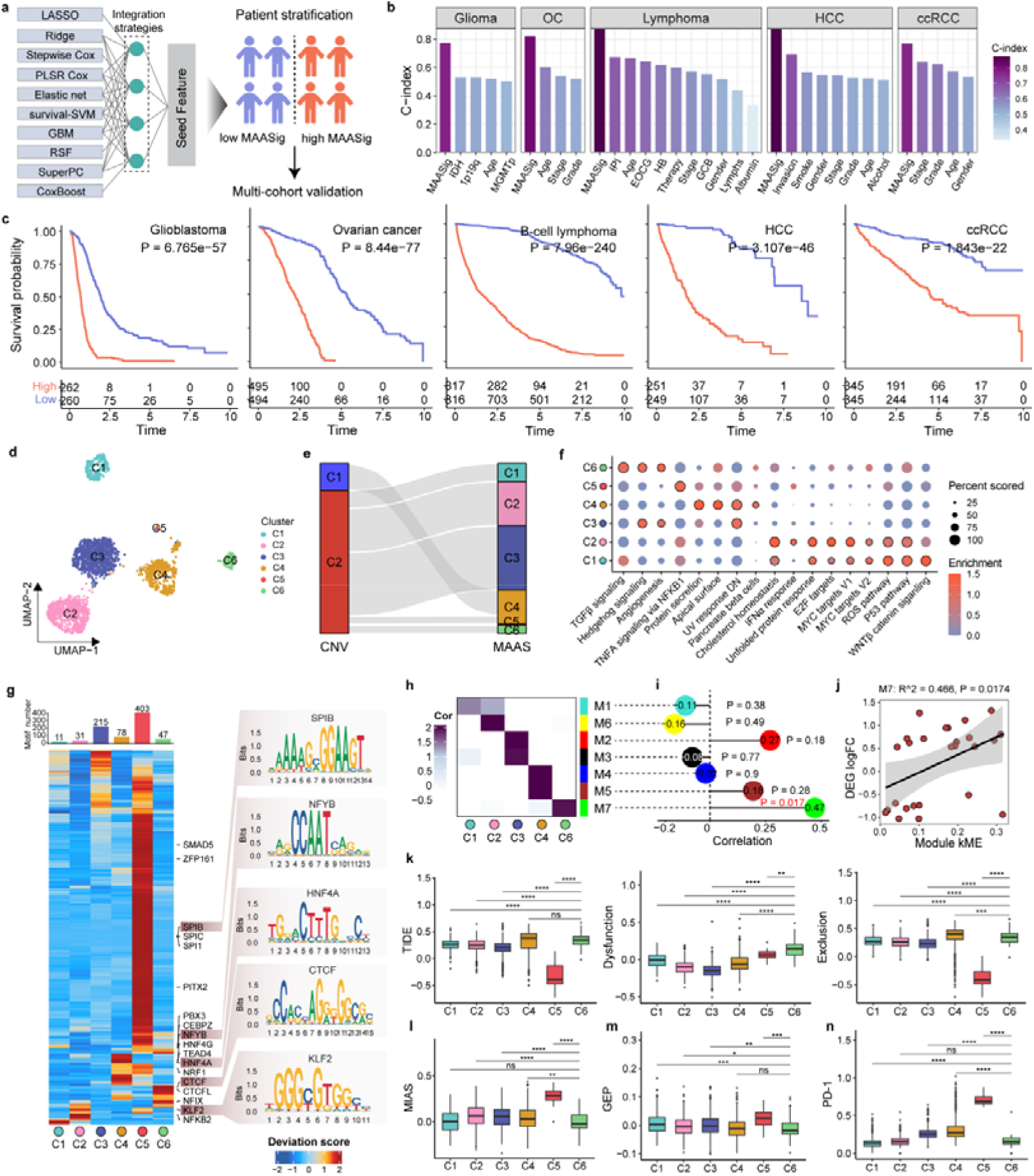
A MAAS-derived clinical signature accurately predict prognosis across multiple cancer types. **a**. Schematic illustration of the workflow for generating the MAAS-derived multimodal signature (MAASig) (see details in the Methods). **b**. Average concordance index (C-index) of traditional clinical features and MAASig for survival prediction across multiple cancer types, including glioblastoma, ovarian cancer (OC), B-cell lymphoma, hepatocellular carcinoma (HCC), and clear cell renal cell carcinoma (ccRCC). c. Kaplan–Meier survival curves demonstrating the clinical relevance of MAASig in a pan-cancer meta-cohort. Datasets for each cancer type were combined into a single cohort, with MAASig stratification determined at the median value. Statistical P-values were calculated using a two-tailed log-rank test. **d**. UMAP embedding of the six tumor cell subpopulations identified by MAAS. **e**. Sankey plot illustrating the clusters identified by CNVs and those identified by MAAS. **f**. Cancer hallmark pathways enriched in each cluster. **g**. Heatmap of chromVAR bias-corrected deviation scores for the differential TF motifs across clusters. The top bar indicates cluster-specific TF motifs with examples of sequence logos for the top TF motifs displayed on the right side of the plot. **h-j** Spearman correlations: (h) between eigengene-based connectivity (kME) of all modules, (i) between module 7 eigengene and kME, and (j) between log fold change (logFC) of differentially expressed genes and anti-PD-1 response versus non-response in patients. Shaded areas represent 95% confidence intervals. **k-n**. Degree of immunotherapeutic response measured by various metrics: (k) tumor immune dysfunction and exclusion (TIDE) scores, (l) MHC I association immunoscore (MIAS), (m) 18-gene expression profiles (GEP), and (n) PDCD1 (PD-1) gene activity. The center line in each box plot represents the median, the lower and upper hinges represent the first and third quartiles, and the whiskers extend to the maximum and minimum values within 1.5 times the interquartile range from the hinge. P-values were determined using the Wilcoxon rank-sum test.

To further examine the clinical significance, we focused on clear cell renal cell carcinoma^44^, which exhibits substantial intra-tumor heterogeneity that contributes to the drug-tolerant state^45^. MAAS accurately identified six cell subpopulations that were previously overlooked by the traditional methods (Fig. 5d-f and Supplementary Fig. 38). To explore potential molecular mechanisms, we examined potential transcription factors binding to these differentially accessible chromatin regions and inferred TF binding motif activity by estimating the gain or loss of chromatin accessibility (Supplementary Methods). Our analysis revealed significant variability in TF activity among the clusters (Fig. 5g). For example, *HNF4A* activity was elevated in cluster 4, while *CTCF* activity increased in cluster 6. Additionally, we assessed the correlation between the cluster-specific gene modules identified by weighted correlation network analysis (WGCNA)^46^ and immunotherapeutic sensitivity (Supplementary Methods). We found that cluster 6, represented by gene module 7, was the most resistant to anti-PD-1 blockade treated by nivolumab (Fig. 5h-j and Supplementary Figs. 39-40). This finding was further validated by the tumor immune dysfunction and exclusion (TIDE) score^47^, MHC I association immunoscore (MIAS)^48^, 18-gene gene expression profile (GEP)^49^, and PD-1 gene activity^50^ (Fig. 5k-n). In summary, MAAS-derived signature has strong prognostic value and robustness in predicting patient survival across many cancer types.

## Discussion

To our knowledge, MAAS is the first computational method for multimodal integration of scATAC-seq data that can identify critical tumor cell subpopulations distinct from those determined by traditional single-modality approaches, such as Copy-scAT^11^ and epiAneufinder^13^. We found that MAAS provides higher accuracy in identifying tumor cell subpopulations and lineage inference compared to other available methods. By integrating multimodal data, MAAS has uncovered new tumor cell subpopulations with significant biological and clinical relevance. The MAAS method is fundamentally different from previous subpopulation prediction methods.

First, MAAS maximizes the extraction of informative features from scATAC-seq data, rather than relying solely on single-modality features, which often overlook other crucial layers of epigenomic information. Additionally, the self-expressive multimodal matrix factorization strategy enhances the multimodal signals, enabling a more robust classification of tumor subpopulations. Furthermore, MAAS is an explainable multi-modality integration method that quantifies the contribution of each modality to cell cluster assignment. For example, the two pediatric ependymoma cell subpopulations predicted by MAAS were primarily driven by chromatin accessibility and SNVs, which had increased weights of 51.64% and 48.03% compared with CNVs, respectively (Fig. 4c). The feasibility of the MAAS method is also noteworthy, as it allows for the simultaneous examination of genetic mutations and epigenetic variations without the need for additional single-cell assays.

In conclusion, the MAAS method underscores the power of multimodal integration in dissecting tumor heterogeneity using single-cell epigenomics data. MAAS will enable the broad application of widely available tumor and other disease single-cell sequencing data, helping to reveal critical tumor cell subpopulations for cell-targeted treatments.

## Methods

### scATAC-seq data analysis

Data included in this study are available from the NCBI Sequence Read Archive. We used the SRA Toolkit (v2.10.9)^51^ to obtain FASTQ files of raw sequencing data, which were then aligned to the GRCh38 reference genome using 10× Genomics Cell Ranger ATAC (v2.1.0, https://support.10xgenomics.com/single-cell-atac/software) software with default parameters. We then used the Signac^52^ package to obtain a cell-by-peak matrix. High-quality cells were retained based on transcription start site enrichment (>3), the number of unique fragments (>1000), percentage of reads in peaks (>15%), blacklist ratio (<5%) and nucleosome signal (<4). To correct batch effects across samples, we performed term frequency-inverse document frequency normalization, which was then included in the ComBat^53^ analysis by the R package sva^54^. In addition, a cell-by-gene score matrix used for functional enrichment analysis was obtained using the GeneActivity function implemented in Signac. DACRs were identified using the FindAllMarkers function by regressing the library size. Details of tumor cell identification were provided in the Supplementary Methods and Supplementary Table 3.

### Mutation calling

We used SComatic for somatic SNV calling (Supplementary Methods). We benchmarked two CNV callers (epiAneufinder^13^ and Copy-scAT^11^) for scATAC-seq data, and we finally used epiAneufinder^13^ for MAAS analysis (Supplementary Note 1; Supplementary Figs. 1-2). Details are provided in the Supplementary Methods.

### Low-rank approximation of the mutation matrix

To dissociate noise and missing values from raw data and accommodate sparsity, we used robust principal component analysis^55^ to obtain low-rank matrices of mutation data. Specifically, we aimed to decompose the raw matrix **X** ^(i)^ into a low-rank matrix 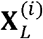 and a noise matrix 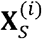 based on an optimization problem:

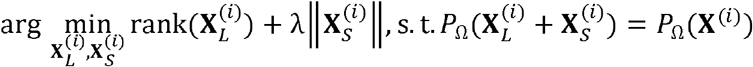

where λ and *P* _Ω_ (·) are linear operators^56, 57^:

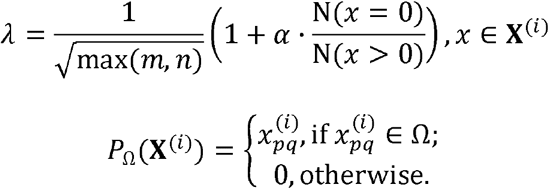

In the above equation, (*m,n*) represents the matrix size and N (*x* = 0) denotes the number of missing values. Under the convex condition, the constrained problem can be written as:

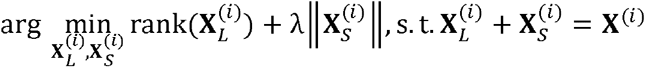

which was resolved by the inexact augmented Lagrange multiplier algorithm^58^..

### Cell affinity estimation in each feature layer

We first corrected the chromatin accessibility profile according to the prior knowledge that copy number gain leads to an aberrant high peak density and vice versa.

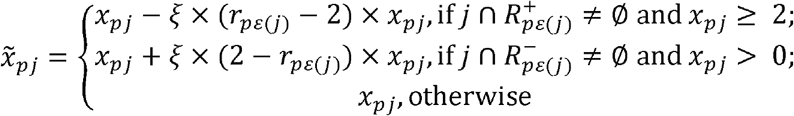

where *x*_*p j*_ and 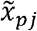 indicate the raw and adjusted peaks of cell, *p* respectively, and *r* _*p ε* (*j*)_ indicates the observed copy numbers containing the region 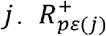 and 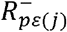 represent the copy number gain and loss region of cell *p*, respectively. Then, we calculated the affinity between cell *p* and *q* (default: cosine). The Hamming distance was used to estimate the cell similarity based on CNVs or SNVs:

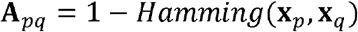

### MAAS structure

We used modified non-negative matrix factorization to jointly integrate multiple modalities for dimension reduction of affinity matrices **A**^(*i*)^The model aimed to seek a consensus low-dimensional space **W** to represent different layers simultaneously and diagonal matrices **H** (*i*) to represent the coefficients of latent factors to be projected in the space:

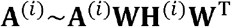

We noted that **A** ^(*i*)^ serve as self-expressive terms, indicating that our multimodal can learn and maintain local structure for subspace clustering. Given the input terms, our model minimizes the loss function as follows:

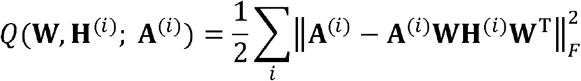

Multiplicative update rules were utilized through the stochastic gradient descent as follows:

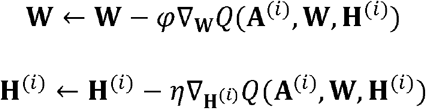

where

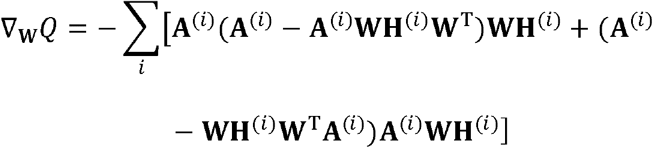

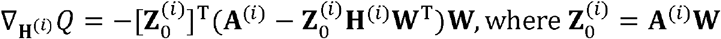

Therefore, we could obtain that

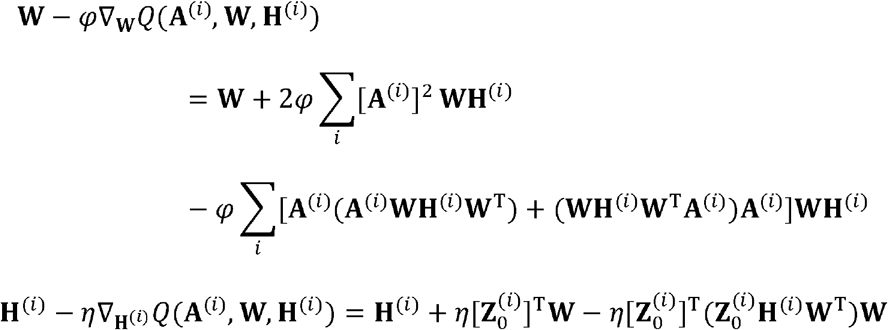

Based on the derivatives, the learning rate for the rule was denoted as

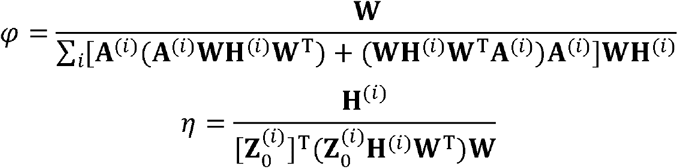

A block coordinate descent scheme was implemented, in which we optimized based on only one rule and kept others fixed. Finally, we implemented the decompositions using hand-solved equations

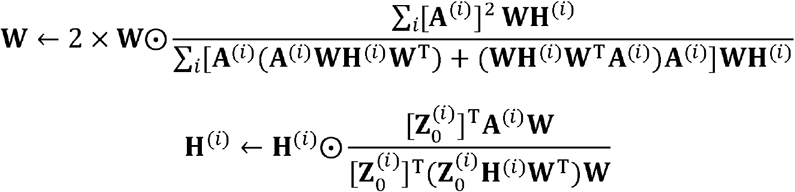

The gradient descent terminated when the condition 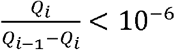 could be met.

The optimized process of our model is summarized in Supplementary Note 2.

### Contribution of modalities

We quantified the contribution of each modality by calculating the trace (the sum of diagonal values).

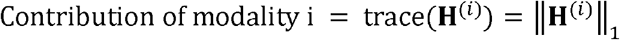

A more considerable trace value indicates a higher weight in cluster assignment.

### Cell clustering

In our study, we applied hierarchical clustering, K-means, and fuzzy C-means clustering to classify tumor subpopulations using a self-defined resolution.

Meanwhile, we designed a comprehensive score *S* based on existing metrics^59-62^ to guide the optimal clustering number

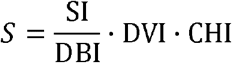

This metric evaluates the distance between inner objects within a subpopulation and estranged objects in distinct clusters. For simplicity of presentation, we omitted the superscript. The details of each term are introduced as follows.

#### 1. Silhouette index (SI)

For sample *i*, the SI is formulated as

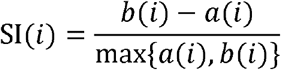

*a* (*i*)denotes the average distance between sample *i* and any other samples within a cluster, and *b*(*i*) denotes the minimum average distance between sample *i* and other clusters.

#### 2. Davies–Bouldin index (DBI)

For clusters *i* and *j*, the DBI is formulated as

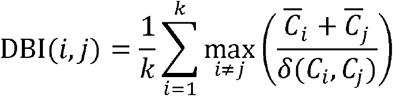

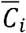 represents the average distance between samples in *i* and the clustering center of *i. δ* (*C*_*i*_, *C*_*j*_) represents the distance between the clustering centers of *i*. and *j*. A smaller DBI indicates a more satisfactory partition.

#### 3. Dunn validity index (DVI)

For clusters *i* and *j*, the DVI is defined as

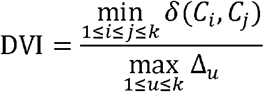

max _1 ≤*u* ≤*k*_ Δ_*u*_ represents the maximum distance between any two samples within the cluster _*u*_

#### 4. Calinski–Harabaz index (CHI)

This index uses covariance to evaluate clustering performance as follows:

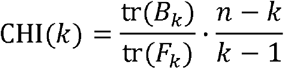

*B*_*k*_ is the inter-cluster covariance matrix, and *F*_*k*_ is the within-cluster covariance matrix. Additionally, *n* and *k* denote the sample size and clustering number. The partition with the largest *S* will be selected for comparison with our model. Uniform Manifold Approximation and Projection (UMAP) embedding was generated based on optimal principal component number determined using the R package findPC^63^.

### Reconstruction of phylogeny

The consensus affinity matrix was first calculated based on **W** by default. It could alternatively be computed based on single modality features. The phylogenetic tree was reconstructed using the minimum evolution algorithm^64^ implemented in the R package ape^65^. We used ggtree^66^ to visualize the phylogeny with the “ape” layout. To estimate performance of cell lineage reconstruction, we calculated the phylogenetic signal with the Blomberg’s *K-*statistic^67^ using the R package phytools^68^. The *K*-statistic ranges from 0 to infinity, with a value of 0 equal to no phylogenetic signal and values greater than 1 equal to strong phylogenetic signal for closely related subpopulations that share more similar features^69^.

### Simulation setup

We randomly generated 5000 accessible chromatin regions, 800 copy number regions, and 300 SNVs. We first created a correlation matrix of cells ranging from 0.3 to 1. The accessible chromatin and copy number regions followed a multivariate Gaussian distribution with a mean value of 4 using the R package MASS. SNVs of each cell were generated following binomial distribution with a probability of 0.3. The correlation matrix consisted of the input covariance matrix for the mvnorm function. To generate clusters with distinct accessible chromatin profiles and without CNV, we increased the counts of 10 regions of a cluster of cells with log2 fold change (FC) values ranging from 0.65 to 1.5 compared with other cells. Thus, this cluster can be considered an additional subpopulation that cannot be distinguished by SNVs or CNVs. The precision is defined as the true positive of the MAAS-identified ground-truth cells within a cluster among all cells of the same cluster.

### Benchmark analysis

We estimated the performance of the UMAP method using hierarchical density-based spatial clustering of applications with noise (HDBSCAN). Specifically, we combined the three single-modality matrices into one meta-matrix for UMAP analysis. We did not use other distance-based clustering methods based on UMAP embedding coordinates since the distance across cells is not directly interpretable. We also compared the accuracy of MAAS with five multi-model-based integrative clustering tools using the default parameters, including intNMF^20^, PintNMF^21^, SNF^22^, LRACluster^23^, and MCIA^24^. The numbers of cells and subpopulations were initially set to 400 and 3, respectively. Each method was run ten times with different random seeds. We also compared the robustness of each method by changing the number of cells (800, 1500, 2000, 2500, and 3000) and subpopulations (from four to eight). We used three metrics, including the normalized mutual information (NMI)^70^, ARI^71^, and V-measure^72^, to evaluate the unsupervised clustering performance. The metrics assess the difference between predicted clusters and reference labels via a quantified score, ranging from 0 to 1 for the AMI and V-measure and from −1 to 1 for the ARI. A higher score indicates a closer match between the obtained and known clustering assignment considering an inferred partition *C* = {*C*_1_,*C*_2_, …*C*_3_}and reference classes *U* = { *U*_1_, *U*_2_, … *U*_3_}. The details of each metric are as follows.

#### 1. AMI

This metric corrects the effect of the agreement solely due to chance between clusters in mutual information (MI).

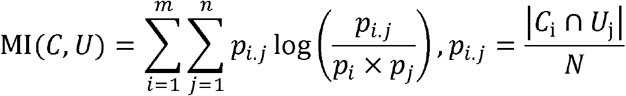

where 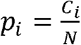 and 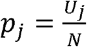 denote the probability of an object from cluster *C*_*i*_ or *U*_*j*_. We then can obtain the representation of the AMI:

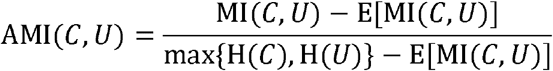

E[·]and H (·)denote the expectation and Shannon entropy, respectively.

#### 2. ARI

Similarly, this metric normalizes the Rand index (RI) to ensure a closer value to 0 for a random partition.

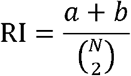

where a represents the number of sample pairs in the same cluster in both *C* and *U*; *b* represents the number of dis-concordant sample pairs. The ARI is denoted as

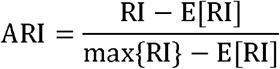

#### 3. V-measure

This metric calculates the harmonic mean of homogeneity and completeness.

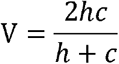

*h* represents homogeneity, which is defined as

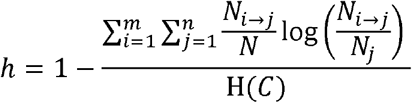

where *N*_*i* →*j*_ denotes the number of samples in cluster *C*_*i*_ divided into *U*_*j*_. Similarly, *c* is defined as

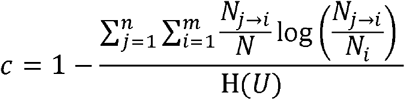

### TMZ response of GBM tumor cell subpopulations

We first used oncoPredict^73^ to predict IC_50_ of GBM patients in the CGGA693, CGGA325 and TCGA cohorts, respectively, by performing linear regression. Specifically, we retrieved drug response information from GDSC2^74^ as the training set, and the three bulk RNA-seq datasets were included in the test set. Genes with a median absolute deviation less than 0.15 were excluded for regression. We then used the calcPhenotype function to predict IC_50_ of each patients. The parameters were set as follows: powerTransformPhenotype was set to FALSE, removeLowVaryGenes was set to 0.2, removeLowVaringGenesFrom was specified as ‘rawData’, and minNumSamples was set to 10. Subsequently, patients were stratified into sensitive and resistant groups at the median cutoff of IC_50_ score. Next, we applied Scissor^3^ to predict the therapeutic phenotype of each cell using binomial regression model. The prediction performance was estimated using the reliability.test function with 1000 permutation times and 10-fold cross-validation. Additionally, we calculated the E-distance between tumor cell clusters and experimentally determined TMZ-sensitive and resistant subpopulations using the edist function from the R package Rfast (https://cran.r-project.org/web/packages/Rfast/) based on the gene activity inferred from scATAC-seq data. To reduce the impact of sample size on distance computation, we randomly selected 100 cells of each cluster and repeated 500 times of the procedure.

### Multidrug sensitivity of PPFE cell subpopulations

We used scRank^75^ to calculate perturbation scores of each drug, including etoposide, vinblastine and vincristine. Edges with weight lower than 0.9 were removed. To identify gene modules across samples, we used the NMF implemented in the R package GeneNMF^76^ by setting the number target NMF components for each sample from 4 to 9, and 10 target meta-modules were determined by hierarchical clustering with minimum confidence of 0.1. We then estimated drug-target module activity using the AddModuleScore function in the Seurat package. Additionally, we used oncoPredict^73^ to calculate the IC_50_ of ependymoma patients from the GSE13267 and GSE66354 cohorts.

### Drug screening

We employed the LINCS^39^ to screen for candidate drugs that can target a TF or kinase. The ExperimentHub (v1.16.1) R package was used to download drug-perturbed results, including a Z-score matrix from differential expression analysis of 12,328 genes across 8140 compound treatments. Only genes with a Z-score < –2 were considered differentially downregulated. The candidate drugs were defined as those that can significantly downregulate TF and are approved for disease treatment by the Food and Drug Administration.

### Functional enrichment analysis

We applied GSVA^77^ to calculate enrichment scores of individual cells for the pathway signatures^78^ based on the curated gene sets (http://www.gsea-msigdb.org/gsea/msigdb/). Differentially enriched pathways were defined with the thresholds of an adjusted P-value < 0.05 and |log_2_FC| > 0.1.

### Survival analysis

We performed several analyses to investigate the prognostic relevance of tumor subpopulation-specific genes. Samples were stratified into two groups based on the median cutoff. Survival curves of the two patient groups were evaluated using the Kaplan – Meier approach. The statistical significance was calculated using a two-tailed log-rank test. We used the survival R package for Cox analysis and the two-tailed Wald test. Features with a P-value < 0.1 in the univariate Cox model were included in the multivariate Cox analysis. Time-dependent AUC and C-index were calculated using the R packages survivalROC and survival, respectively.

## Data availability

The raw glioma, PPFE, and ccRCC scATAC-seq data are available in the NCBI database (https://www.ncbi.nlm.nih.gov/geo/) under accession numbers GSE139136 (GBM), GSE63655 (pGBM), GSE206579 and PRJNA768891, respectively. The raw ovarian cancer scATAC-seq data is available via the database of Genotypes and Phenotypes (dbGaP) under the accession number phs002340.v1.p1. The raw B-cell lymphoma scATAC-seq data are available from 10x Genomics (https://www.10xgenomics.com/resources/datasets/). The scATAC-seq dataset for the SNU601 cell line is available from the NCBI database under the accession number PRJNA674903, and the single-cell whole-genome sequencing data for the SNU601 cell line is available under the accession number PRJNA498809. The tumor samples of patients SU006 and SU008 are available in the NCBI database under the accession number PRJNA533341. Gene expression and clinical features of patients from The Cancer Genome Atlas (TCGA) cohort are available from the GDC portal (https://portal.gdc.cancer.gov/) and as reported by Liu et al^79^. Gene expression profiles of patients from the two cohorts CGGA693 and CGGA325 are publicly available from the Chinese Glioma Genome Atlas (http://www.cgga.org.cn/). Gene expression of experimentally determined wild-type and TMZ-resistant glioma cells were obtained from the NCBI database under the accession numbers GSE53014 and GSE68029. Two bulk RNA-seq datasets of PPFE were obtained from the NCBI database under the accession numbers GSE13267 and GSE66354. Bulk ATAC-seq data of B-cell lymphoma cell lines were obtained from the NCBI database under the accession number GSE254913. Datasets used for clinical signature analysis from the NCBI database are available under the following accession numbers: (1) ovarian cancer, GSE140082, and GSE32062; (2) B-cell lymphoma, GSE10846, and GSE136971; (3) hepatocellular carcinoma, GSE116174, and GSE76427. Gene expression and clinical information of ccRCC patients from the E-MTAB-1980 and CPTAC cohorts were retrieved from the European Bioinformatics Institute (https://www.ebi.ac.uk/) and LinkedOmics (https://kb.linkedomics.org/download), respectively.

## Code availability

The open-source MAAS is available from the following Github repository: https://github.com/Larrycpan/MAAS/.

## Acknowledgments

We thank Dr. Zheng Hu at the Chinese Academy of Sciences, Dr. Chenfei Wang at the Tongji University and members of the Li Laboratory for their helpful discussions. This work was supported by the National Natural Science Foundation of China (No. 32370721, 32100533) to L.L. We also thank Dr. Zhiqiang Ye at the Shenzhen Bay Laboratory Supercomputing Center for high-level computing support.

## Author contributions

L.L. conceived and supervised the project. K.X., R.D., and Y.Q. performed the bioinformatics analysis. J.W. and C.Y. contributed to the cancer analysis. K.X., R.D., X.Z., and L.L. wrote the manuscript with assistance from the other authors.

## Supplementary Data

**Supplementary Data 1**. List of differential chromatin accessible regions between SNU601 gastric cancer 2b identified by MAAS.

**Supplementary Data 2**. List of glioma tumor-specific SNVs.

**Supplementary Data 3**. List of differential chromatin accessible regions of glioma cluster 1 identified by MAAS.

**Supplementary Data 4**. List of differential chromatin accessible regions of pediatric posterior fossa ependymoma cluster 2 identified by MAAS.

**Supplementary Data 5**. List of genes in the drug-target gene module.

**Supplementary Data 6**. List of PPFE cluster 2-specific TFs and kinases included in the LINCS consortium.

**Supplementary Data 7**. List of target genes of ESR1 in the PPFE cluster 2.

**Supplementary Data 8**. List of differential chromatin accessible regions of B-cell lymphoma clusters identified by MAAS.

## References

1. Mazor, T. et al. DNA Methylation and Somatic Mutations Converge on the Cell Cycle and Define Similar Evolutionary Histories in Brain Tumors. Cancer Cell 28, 307–317 (2015).

2. Dagogo-Jack, I. & Shaw, A.T. Tumour heterogeneity and resistance to cancer therapies. Nat Rev Clin Oncol 15, 81–94 (2018).

3. Sun, D. et al. Identifying phenotype-associated subpopulations by integrating bulk and single-cell sequencing data. Nat Biotechnol 40, 527–538 (2022).

4. Zhao, J. et al. Detection of differentially abundant cell subpopulations in scRNA-seq data. Proc Natl Acad Sci U S A 118 (2021).

5. Levine, J.H. et al. Data-Driven Phenotypic Dissection of AML Reveals Progenitor-like Cells that Correlate with Prognosis. Cell 162, 184–197 (2015).

6. Gao, R. et al. Delineating copy number and clonal substructure in human tumors from single-cell transcriptomes. Nat Biotechnol 39, 599–608 (2021).

7. Zhou, Z., Xu, B., Minn, A. & Zhang, N.R. DENDRO: genetic heterogeneity profiling and subclone detection by single-cell RNA sequencing. Genome Biol 21, 10 (2020).

8. Corces, M.R. et al. Lineage-specific and single-cell chromatin accessibility charts human hematopoiesis and leukemia evolution. Nat Genet 48, 1193–1203 (2016).

9. Corces, M.R. et al. The chromatin accessibility landscape of primary human cancers. Science 362 (2018).

10. Buenrostro, J.D. et al. Integrated Single-Cell Analysis Maps the Continuous Regulatory Landscape of Human Hematopoietic Differentiation. Cell 173, 1535–1548 e1516 (2018).

11. Nikolic, A. et al. Copy-scAT: Deconvoluting single-cell chromatin accessibility of genetic subclones in cancer. Sci Adv 7, eabg6045 (2021).

12. Wu, C.Y. et al. Integrative single-cell analysis of allele-specific copy number alterations and chromatin accessibility in cancer. Nat Biotechnol 39, 1259–1269 (2021).

13. Ramakrishnan, A. et al. epiAneufinder identifies copy number alterations from single-cell ATAC-seq data. Nat Commun 14, 5846 (2023).

14. Choi, J.D. & Lee, J.S. Interplay between Epigenetics and Genetics in Cancer. Genomics Inform 11, 164–173 (2013).

15. Nam, A.S., Chaligne, R. & Landau, D.A. Integrating genetic and non-genetic determinants of cancer evolution by single-cell multi-omics. Nat Rev Genet 22, 3–18 (2021).

16. De Falco, A., Caruso, F., Su, X.D., Iavarone, A. & Ceccarelli, M. A variational algorithm to detect the clonal copy number substructure of tumors from scRNA-seq data. Nat Commun 14, 1074 (2023).

17. Chen, H. et al. Assessment of computational methods for the analysis of single-cell ATAC-seq data. Genome Biol 20, 241 (2019).

18. Zhang, J. et al. in 2019 IEEE/CVF Conference on Computer Vision and Pattern Recognition (CVPR) 5468–5477 (2019).

19. McInnes, L., Healy, J. & Melville, J. Umap: Uniform manifold approximation and projection for dimension reduction. arXiv preprint 1802.03426 (2018).

20. Chalise, P. & Fridley, B.L. Integrative clustering of multi-level ‘omic data based on non-negative matrix factorization algorithm. PLoS One 12, e0176278 (2017).

21. Pierre-Jean, M., Mauger, F., Deleuze, J.F. & Le Floch, E. PIntMF: Penalized Integrative Matrix Factorization method for multi-omics data. Bioinformatics 38, 900–907 (2022).

22. Wang, B. et al. Similarity network fusion for aggregating data types on a genomic scale. Nat Methods 11, 333–337 (2014).

23. Wu, D., Wang, D., Zhang, M.Q. & Gu, J. Fast dimension reduction and integrative clustering of multi-omics data using low-rank approximation: application to cancer molecular classification. BMC Genomics 16, 1022 (2015).

24. Meng, C., Kuster, B., Culhane, A.C. & Gholami, A.M. A multivariate approach to the integration of multi-omics datasets. BMC Bioinformatics 15, 162 (2014).

25. Zahavi, D.J. et al. Antibody dependent cell-mediated cytotoxicity selection pressure induces diverse mechanisms of resistance. Cancer Biol Ther 24, 2269637 (2023).

26. Regner, M.J. et al. A multi-omic single-cell landscape of human gynecologic malignancies. Mol Cell 81, 4924–4941 e4910 (2021).

27. Hara, T. et al. Interactions between cancer cells and immune cells drive transitions to mesenchymal-like states in glioblastoma. Cancer Cell 39, 779–792 e711 (2021).

28. Székely, G.J. & Rizzo, M.L. Energy statistics: A class of statistics based on distances. Journal of Statistical Planning and Inference 143, 1249–1272 (2013).

29. Hiddingh, L. et al. EFEMP1 induces gamma-secretase/Notch-mediated temozolomide resistance in glioblastoma. Oncotarget 5, 363–374 (2014).

30. Tso, J.L. et al. Bone morphogenetic protein 7 sensitizes O6-methylguanine methyltransferase expressing-glioblastoma stem cells to clinically relevant dose of temozolomide. Mol Cancer 14, 189 (2015).

31. Yang, Y. & Yang, L. Somatic structural variation signatures in pediatric brain tumors. Cell Rep 42, 113276 (2023).

32. Aubin, R.G. et al. Pro-inflammatory cytokines mediate the epithelial-to-mesenchymal-like transition of pediatric posterior fossa ependymoma. Nat Commun 13, 3936 (2022).

33. Cao, J. et al. The single-cell transcriptional landscape of mammalian organogenesis. Nature 566, 496–502 (2019).

34. Rojo de la Vega, M., Chapman, E. & Zhang, D.D. NRF2 and the Hallmarks of Cancer. Cancer Cell 34, 21–43 (2018).

35. Jurikova, M., Danihel, L., Polak, S. & Varga, I. Ki67, PCNA, and MCM proteins: Markers of proliferation in the diagnosis of breast cancer. Acta Histochem 118, 544–552 (2016).

36. Aiello, N.M. & Kang, Y. Context-dependent EMT programs in cancer metastasis. J Exp Med 216, 1016–1026 (2019).

37. Katoh, M. Function and cancer genomics of FAT family genes (review). Int J Oncol 41, 1913–1918 (2012).

38. Li, C. et al. scRank infers drug-responsive cell types from untreated scRNA-seq data using a target-perturbed gene regulatory network. Cell Rep Med, 101568 (2024).

39. Subramanian, A. et al. A Next Generation Connectivity Map: L1000 Platform and the First 1,000,000 Profiles. Cell 171, 1437–1452 e1417 (2017).

40. Steele, C.D. et al. Signatures of copy number alterations in human cancer. Nature 606, 984–991 (2022).

41. Flash-Frozen Lymph Node with B Cell Lymphoma. 10X Genomics https://support.10xgenomics.com/single-cell-multiome-atac-gex/datasets (2021).

42. Liau, B.B. et al. Adaptive Chromatin Remodeling Drives Glioblastoma Stem Cell Plasticity and Drug Tolerance. Cell Stem Cell 20, 233–246 e237 (2017).

43. Garcia-Alonso, L., Holland, C.H., Ibrahim, M.M., Turei, D. & Saez-Rodriguez, J. Benchmark and integration of resources for the estimation of human transcription factor activities. Genome Res 29, 1363–1375 (2019).

44. Long, Z. et al. Single-cell multiomics analysis reveals regulatory programs in clear cell renal cell carcinoma. Cell Discov 8, 68 (2022).

45. Posadas, E.M., Limvorasak, S. & Figlin, R.A. Targeted therapies for renal cell carcinoma. Nat Rev Nephrol 13, 496–511 (2017).

46. Morabito, S., Reese, F., Rahimzadeh, N., Miyoshi, E. & Swarup, V. hdWGCNA identifies co-expression networks in high-dimensional transcriptomics data. Cell Rep Methods 3, 100498 (2023).

47. Jiang, P. et al. Signatures of T cell dysfunction and exclusion predict cancer immunotherapy response. Nat Med 24, 1550–1558 (2018).

48. Wu, C.C., Wang, Y.A., Livingston, J.A., Zhang, J. & Futreal, P.A. Prediction of biomarkers and therapeutic combinations for anti-PD-1 immunotherapy using the global gene network association. Nat Commun 13, 42 (2022).

49. Cristescu, R. et al. Pan-tumor genomic biomarkers for PD-1 checkpoint blockade-based immunotherapy. Science 362 (2018).

50. Jiang, Y., Chen, M., Nie, H. & Yuan, Y. PD-1 and PD-L1 in cancer immunotherapy: clinical implications and future considerations. Hum Vaccin Immunother 15, 1111–1122 (2019).

51. Leinonen, R., Sugawara, H., Shumway, M. & International Nucleotide Sequence Database, C. The sequence read archive. Nucleic Acids Res 39, D19–21 (2011).

52. Stuart, T., Srivastava, A., Madad, S., Lareau, C.A. & Satija, R. Single-cell chromatin state analysis with Signac. Nat Methods 18, 1333–1341 (2021).

53. Johnson, W.E., Li, C. & Rabinovic, A. Adjusting batch effects in microarray expression data using empirical Bayes methods. Biostatistics 8, 118–127 (2007).

54. Leek, J.T., Johnson, W.E., Parker, H.S., Jaffe, A.E. & Storey, J.D. The sva package for removing batch effects and other unwanted variation in high-throughput experiments. Bioinformatics 28, 882–883 (2012).

55. Shang, F., Liu, Y., Cheng, J. & Cheng, H. in Proceedings of the 23rd ACM International Conference on Conference on Information and Knowledge Management 1149–1158 (2014).

56. Candes, E.J., Xiaodong, L.I., Yl, M.A. & Wright, J. Robust Principal Component Analysis? Journal of the ACM (JACM) 58 (2011).

57. Chen, Z., Gong, F., Wan, L. & Ma, L. RobustClone: a robust PCA method for tumor clone and evolution inference from single-cell sequencing data. Bioinformatics 36, 3299–3306 (2020).

58. Lin, Z., Chen, M. & Ma, Y. The Augmented Lagrange Multiplier Method for Exact Recovery of Corrupted Low-Rank Matrices. ArXiv abs/1009.5055 (2010).

59. Rousseeuw, P.J. Silhouettes: A graphical aid to the interpretation and validation of cluster analysis. Journal of Computational and Applied Mathematics 20, 53–65 (1987).

60. Davies, D.L. & Bouldin, D.W. A Cluster Separation Measure. IEEE Transactions on Pattern Analysis and Machine Intelligence PAMI-1, 224–227 (1979).

61. Pakhira, M.K., Bandyopadhyay, S. & Maulik, U. Validity index for crisp and fuzzy clusters. Pattern Recognition 37, 487–501 (2004).

62. Caliński, T. & Harabasz, J. A dendrite method for cluster analysis. Communications in Statistics 3, 1–27 (1974).

63. Zhuang, H., Wang, H. & Ji, Z. findPC: An R package to automatically select the number of principal components in single-cell analysis. Bioinformatics 38, 2949–2951 (2022).

64. Rzhetsky, A. & Nei, M. Theoretical foundation of the minimum-evolution method of phylogenetic inference. Mol Biol Evol 10, 1073–1095 (1993).

65. Paradis, E., Claude, J. & Strimmer, K. APE: Analyses of Phylogenetics and Evolution in R language. Bioinformatics 20, 289–290 (2004).

66. Yu, G. Using ggtree to Visualize Data on Tree-Like Structures. Curr Protoc Bioinformatics 69, e96 (2020).

67. Blomberg, S.P., Garland, T., Jr. & Ives, A.R. Testing for phylogenetic signal in comparative data: behavioral traits are more labile. Evolution 57, 717–745 (2003).

68. Revell, L.J. phytools 2.0: an updated R ecosystem for phylogenetic comparative methods (and other things). PeerJ 12, e16505 (2024).

69. Olival, K.J. et al. Host and viral traits predict zoonotic spillover from mammals. Nature 546, 646–650 (2017).

70. Meilă, M. Comparing clusterings—an information based distance. Journal of Multivariate Analysis 98, 873–895 (2007).

71. Hubert, L. & Arabie, P. Comparing partitions. Journal of Classification 2, 193–218 (1985).

72. Rosenberg, A. & Hirschberg, J. in Conference on Empirical Methods in Natural Language Processing (2007).

73. Maeser, D., Gruener, R.F. & Huang, R.S. oncoPredict: an R package for predicting in vivo or cancer patient drug response and biomarkers from cell line screening data. Brief Bioinform 22 (2021).

74. Yang, W. et al. Genomics of Drug Sensitivity in Cancer (GDSC): a resource for therapeutic biomarker discovery in cancer cells. Nucleic Acids Res 41, D955–961 (2013).

75. Li, C. et al. scRank infers drug-responsive cell types from untreated scRNA-seq data using a target-perturbed gene regulatory network. Cell Rep Med 5, 101568 (2024).

76. Yerly, L. et al. Wounding triggers invasive progression in human basal cell carcinoma. bioRxiv, 2024.2005.2031.596823 (2024).

77. Hanzelmann, S., Castelo, R. & Guinney, J. GSVA: gene set variation analysis for microarray and RNA-seq data. BMC Bioinformatics 14, 7 (2013).

78. Liberzon, A. et al. The Molecular Signatures Database (MSigDB) hallmark gene set collection. Cell Syst 1, 417–425 (2015).

79. Liu, J. et al. An Integrated TCGA Pan-Cancer Clinical Data Resource to Drive High-Quality Survival Outcome Analytics. Cell 173, 400–416 e411 (2018).

