## Supplementary Information for "Multimodal integration of single cell ATAC-seq data enables highly accurate delineation of clinically relevant tumor cell subpopulations"

**Contents**

Supplementary Note 1. Benchmark of mutation calling from scATAC-seq data.

Supplementary Note 2. Pseudo-code of the update process of MAAS.

Supplementary Methods.

Supplementary Figures 1-40.

Supplementary Tables 1-3.

**Supplementary Note 1. Benchmark of mutation calling from scATAC-seq data**

We benchmarked two existing tools for CNV calling, epiAneufinder^1^ and Copy-scAT^2^, using a gastric cancer cell line with matched single-cell whole genome sequencing (scWGS) data^3^. The results showed that epiAneufinder demonstrated similar accuracy to Copy-scAT (Supplementary Fig. 1). To further quantify the performance of these tools, we calculated the Spearman correlation coefficient of the consensus copy number value for each chromosome measured by scATAC-seq and scWGS data. The results indicated that the output of epiAneufinder was highly correlated with the pseudo-bulk copy numbers derived from DNA-seq data (Spearman correlation: R = 0.938, *P*-value = 1.7×10^-19^), slightly outperformed Copy-scAT (Spearman correlation: R = 0.877, *P*-value = 5.14×10^-14^; Supplementary Fig. 2a,b). A further comparison in sample SU006 demonstrated that epiAneufinder (Spearman correlation: R = 0.683, *P*-value = 1.2×10^-5^) significantly outperformed Copy-scAT (Spearman correlation: R = 0.241, *P*-value = 0.177; Supplementary Fig. 2c,d). In another sample, SU008, epiAneufinder again showed a higher correlation with ground truth compared to Copy-scAT (Supplementary Fig. 2e,f). Overall, we have verified the accuracy of epiAneufinder for CNV calling from scATAC-seq data and selected it as the preferred CNV calling method in our study.

**Supplementary Note 2. Pseudo-code of the update process of MAAS.**

**Algorithm of MAAS**

**Input:** Affinity matrix
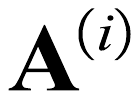
 of peak, CNV, and SNV, maximum dimension *k*

**Output:** Fundamental matrix
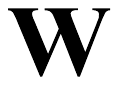
, and coefficient matrices
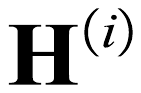
;

1. Initialize
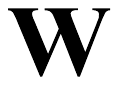
,
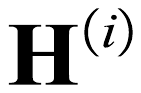
 randomly and *k*;

2. **for** *l* = 2 to *k*

3. **while** not converged **do**

4.
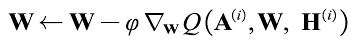

5.
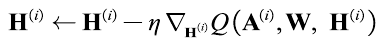

6. Determine the loss function
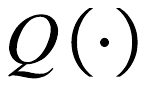

7. **until**
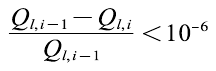

8. **end while**

9. **end for**

**return**
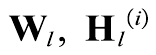

**Supplementary Methods**.

**CNV calling from scATAC-seq data.** We used the R package epiAneufinder^1^ to estimate CNVs from scATAC-seq data. Cells with the number of fragments less than 20,000 were filtered out. Next, the genome was binned into equally sized windows (100,000 bp) and removed from the ENCODE blacklist^4^. Aneuploid cells with known markers were considered tumor cells. We also used Copy-scAT^2^ using the function identifyCNVClusters and the default parameters.

**CNV calling from DNA-seq data.** For scWGS data, we used the R package CopyKit^5^ to estimate CNVs with the parameters variable bins = 1Mb, and removed cells having less than two mean bin counts. For WES data, we used VarScan2^6^ for CNV calling using default parameters.

**Somatic SNV calling from scATAC-seq data.** We used SComatic^7^ to detect somatic mutations for MAAS analysis from scATAC-seq data. We first split the alignment file into cell-type-specific BAM files with at least 30 minimum mapping quality. Subsequently, the base count information for each cell type and each position in the genome was recorded in a base count matrix indexed by cell types and genomic coordinates provided by the ATAC bed file. We required at least five cells to consider a genomic site for further analysis. The minimum base quality permited for the base counts was 20, and the minimum mapping quality required is 30 for this step. Next, we merged the base count matrices generated in the previous step for all cell types analysed into a single base count matrix. Afterwards, we called somatic SNVs with default parameters. Finally, we filtered variants based on the panel of germline polymorphisms and recurrent artefacts of scATAC-seq data, which was retrieved from the public GitHub repository (https://github.com/cortes-ciriano-lab/SComatic/blob/main/PoNs). To further alleviate noises, we included only SNVs occurring in at least ten tumor cells.

**Identification of tumor cells.** We used marker genes and aberrant copy numbers to identify tumor cells. The markers for each cancer are provided in Supplementary Table 3. Samples with more than 300 tumor cells and 30 SNVs were retained for MAAS analysis.

**Validation of C1 identified in the aGBM dataset.** We validated driven factors and drug resistance for the MAAS-identified cluster C1 in adult aGBM in a pGBM dataset^2^. To identify C1- and C2-like clusters in pGBM, we first extracted accessible chromatin regions that overlapped DACRs of aGBM clusters 1 and 2 with at least 200bp, which were included in *K*-means clustering for pGBM tumor cells.

**Module score calculation.** We used the AddModuleScore function implemented in the Seurat package^8^ to calculate enrichment score of a specific gene set (e.g. proliferation, migration and drug-target gene module) with 50 control cells and 24 bins of aggregate expression levels for all analyzed features.

**Sample deconvolution for bulk RNA-seq.** We used the R package MuSiC^9^ to calculate proportions of MAAS-determined subpopulations with maximum of 1000 iterations, taking the single-cell gene activity as the reference.

**Perturbation analysis.** We employed CellOracle^10^ to predict cell fate changes upon *ESR1* knockout *in silico* based on existing TF binding data^11^ for gene regulatory network construction. To simulate the knockout, we set the gene expression to 0. We then calculated the probability of cell cluster transitions based on the simulated data using the estimate_transition_prob function with 200 nearest neighbors, and the calculate_embedding_shift function with the parameter “sigma_corr” set to 0.05.

**Processing of bulk ATAC-seq of B-cell lymphoma cell lines.** The raw FASTQ files were generated using SRA Toolkit (v2.10.9)^12^. low-quality reads and the adapters were removed by Trim Galore (v0.6.6, https://www.bioinformatics.babraham.ac.uk/projects/trim_galore/) with the following parameters: --quality 20 and --paired. High-quality remaining reads were aligned to the reference of human genome GRCh38 using BWA-MEM (v.0.7.17)^13^. Subsequently, duplicated reads were removed using Picard (https://broadinstitute.github.io/picard/). We then removed the Tn5 adapters with alignmentSieve (v.3.3.2)^14^ with the parameter ‘ATACshift’ and indexed the final BAM files with SAMtools (v1.10)^15^. Finally, we used MACS2 for peak calling with the following parameters: --shift -100, --extsize 200, -f BAM, --SPMR -B, -g hs and --nomodel, which was then normalized. Additionally, we only retained overlapped accessible chromatin regions to calculate the E-distance between scATAC-seq and bulk ATAC-seq chromatin profiles.

**Single-cell TF binding motif activity analysis.** Single-cell TF motif activity was calculated for a set of 870 TFs from the Catalog of Inferred Sequence Binding Preferences (CIS-BP) database (from chromVAR motifs ‘human_pwms_v2’) using the RunChromVAR function implemented in Signac^16^. Differential TF activity across clusters were determined by the FindMarkers function (log_2_FC > 1 and FDR < 0.05).

**Single-cell WGCNA.** We used the R package hdWGCNA^17^ to perform single-cell WGCNA, taking the cluster assignment as the "phenotype". Specifically, we first constructed metacells from the scATAC-seq data based on inferred gene expression counts by the k-nearest neighborhood algorithm (k = 25). The maximum number of shared cells between two metacells was set to 10. We then applied the function TestSoftPowers to perform a parameter sweep to automatically determine the optimal soft power threshold, which was further used to constructing the co-expression network for identifying gene modules. The grey module was ignored for all downstream analysis and interpretation. To identify the hub genes of each module, we calculated the eigengene-based connectivity (kME) of each gene using the ModuleConnectivity function that essentially computes pairwise correlations between genes and module eigengenes.

**Development of MAASig based on integrative strategies.** We first estimated the TF activity from bulk RNA-seq data using the R package decoupleR^18^ based on the CollecTRI meta-resource^19^. Next, we combined gene expression and activity of all subpopulation-specific TFs of bulk RNA-seq data. To screen for potential prognostic feature, we first used univariate Cox proportional hazard models with a significance threshold of P-value < 0.01 in the training cohort. Subsequently, we integrated 10 machine learning methods, including least absolute shrinkage and selection operator (LASSO) regression^20^, ridge regression^21^, stepwise Cox regression^22^, partial least squares regression for Cox (PLSR Cox)^23^, elastic net^24^, survival support vector machine (survival-SVM)^25^, generalized boosted regression model (GBM)^26^, random survival forest (RSF)^27^, supervised principal components (SuperPC)^28^ and CoxBoost^29^, followed by the previous study^30^. Four algorithms, including LASSO, stepwise Cox, RSF, and CoxBoost, were employed to perform feature selection, which was included in the further 101 prediction strategies to obtain an optimal model. Models with less than five genes were excluded for performance estimation. All the algorithms were implemented via R packages. The LASSO, ridge and elastic net were performed via the glmnet package. The regularization parameter *λ* was determined via 10-fold leave-one-out cross-validation, while the L1-L2 regularization trade-off parameter *α*, was set from 0 to 1 in increments of 0.1. The stepwise Cox algorithm was performed by the survival package. The stepwise Cox algorithm was performed by the survival package using the Akaike information criterion, and the direction set as both, backward, and forward, respectively. The plsRcox model was analyzed by the plsRcox package with 10 components. The survival-SVM model was performed by survivalsvm package with regularization set to 1. The quadratic optimization problem was solved by the “ipop” algorithm. The GBM model was performed by the gbm package. First, the gbm function was used to select optimal number of trees with minimum cross-validation error with the following parameters: interaction.depth=3, n.minobsinnode=10 and shrinkage=0.001. Then, the gbm function was reused to fit the final GBM model. The SuperPC model was performed using the superpc package. Specifically, the superpc.train function was first to train an initial model with 50% of standard deviation values added to the denominator. The superpc.cv function was then used to estimate the optimal feature threshold by 10-fold cross-validation and default parameters. The CoxBoost model was implemented by the CoxBoost package. Specifically, we used the optimCoxBoostPenalty function to first determine the optimal penalty with start penalty of 500. Next, the number of boosting steps to perform was selected using the function cv.CoxBoost with the following parameters: maxstepno=500, K=10 and type="verweij". The dimension of the selected multivariate Cox model was finally determined by the CoxBoost function. Patients were stratified into two subtypes at the median cutoff. Sample with survival time less than 30 days or larger than 10 years were excluded for analysis.

**Reference**

1.	Ramakrishnan, A. et al. epiAneufinder identifies copy number alterations from single-cell ATAC-seq data. *Nat Commun* **14**, 5846 (2023).

2.	Nikolic, A. et al. Copy-scAT: Deconvoluting single-cell chromatin accessibility of genetic subclones in cancer. *Sci Adv* **7**, eabg6045 (2021).

3.	Wu, C.Y. et al. Integrative single-cell analysis of allele-specific copy number alterations and chromatin accessibility in cancer. *Nat Biotechnol* **39**, 1259-1269 (2021).

4.	Amemiya, H.M., Kundaje, A. & Boyle, A.P. The ENCODE Blacklist: Identification of Problematic Regions of the Genome. *Sci Rep* **9**, 9354 (2019).

5.	Minussi, D.C. et al. Resolving clonal substructure from single cell genomic data using CopyKit. *bioRxiv*, 2022.2003.2009.483497 (2022).

6.	Koboldt, D.C. et al. VarScan 2: somatic mutation and copy number alteration discovery in cancer by exome sequencing. *Genome Res* **22**, 568-576 (2012).

7.	Muyas, F. et al. De novo detection of somatic mutations in high-throughput single-cell profiling data sets. *Nat Biotechnol* **42**, 758-767 (2024).

8.	Butler, A., Hoffman, P., Smibert, P., Papalexi, E. & Satija, R. Integrating single-cell transcriptomic data across different conditions, technologies, and species. *Nat Biotechnol* **36**, 411-420 (2018).

9.	Wang, X., Park, J., Susztak, K., Zhang, N.R. & Li, M. Bulk tissue cell type deconvolution with multi-subject single-cell expression reference. *Nat Commun* **10**, 380 (2019).

10.	Kamimoto, K. et al. Dissecting cell identity via network inference and in silico gene perturbation. *Nature* **614**, 742-751 (2023).

11.	Paul, F. et al. Transcriptional Heterogeneity and Lineage Commitment in Myeloid Progenitors. *Cell* **163**, 1663-1677 (2015).

12.	Leinonen, R., Sugawara, H., Shumway, M. & International Nucleotide Sequence Database, C. The sequence read archive. *Nucleic Acids Res* **39**, D19-21 (2011).

13.	Li, H. & Durbin, R. Fast and accurate short read alignment with Burrows-Wheeler transform. *Bioinformatics* **25**, 1754-1760 (2009).

14.	Ramirez, F. et al. deepTools2: a next generation web server for deep-sequencing data analysis. *Nucleic Acids Res* **44**, W160-165 (2016).

15.	Li, H. et al. The Sequence Alignment/Map format and SAMtools. *Bioinformatics* **25**, 2078-2079 (2009).

16.	Stuart, T., Srivastava, A., Madad, S., Lareau, C.A. & Satija, R. Single-cell chromatin state analysis with Signac. *Nat Methods* **18**, 1333-1341 (2021).

17.	Morabito, S., Reese, F., Rahimzadeh, N., Miyoshi, E. & Swarup, V. hdWGCNA identifies co-expression networks in high-dimensional transcriptomics data. *Cell Rep Methods* **3**, 100498 (2023).

18.	Badia, I.M.P. et al. decoupleR: ensemble of computational methods to infer biological activities from omics data. *Bioinform Adv* **2**, vbac016 (2022).

19.	Muller-Dott, S. et al. Expanding the coverage of regulons from high-confidence prior knowledge for accurate estimation of transcription factor activities. *Nucleic Acids Res* **51**, 10934-10949 (2023).

20.	Tibshirani, R. Regression Shrinkage and Selection via the Lasso. *Journal of the Royal Statistical Society. Series B (Methodological)* **58**, 267-288 (1996).

21.	Hilt, D.E., Seegrist, D.W., United States. Forest, S. & Northeastern Forest Experiment, S. Ridge, a computer program for calculating ridge regression estimates, Vol. no.236. (Dept. of Agriculture, Forest Service, Northeastern Forest Experiment Station, Upper Darby, Pa; 1977).

22.	Asano, J., Hirakawa, A. & Hamada, C. Assessing the prediction accuracy of cure in the Cox proportional hazards cure model: an application to breast cancer data. *Pharmaceutical Statistics* **13**, 357 - 363 (2014).

23.	Nguyen, D.V. & Rocke, D.M. Partial least squares proportional hazard regression for application to DNA microarray survival data. *Bioinformatics* **18**, 1625-1632 (2002).

24.	Zou, H. & Hastie, T. Regularization and Variable Selection Via the Elastic Net. *Journal of the Royal Statistical Society Series B: Statistical Methodology* **67**, 301-320 (2005).

25.	Van Belle, V., Pelckmans, K., Suykens, J. & Van Huffel, S. in The Third International Conference on Computational Intelligence in Medicine and Healthcare (CIMED2007) (2007).

26.	Jerome, H.F. Greedy function approximation: A gradient boosting machine. *The Annals of Statistics* **29**, 1189-1232 (2001).

27.	Hemant, I., Udaya, B.K., Eugene, H.B. & Michael, S.L. Random survival forests. *The Annals of Applied Statistics* **2**, 841-860 (2008).

28.	Bair, E., Hastie, T., Paul, D. & Tibshirani, R. Prediction by Supervised Principal Components. *Journal of the American Statistical Association* **101**, 119-137 (2006).

29.	Binder, H., Allignol, A., Schumacher, M. & Beyersmann, J. Boosting for high-dimensional time-to-event data with competing risks. *Bioinformatics* **25**, 890-896 (2009).

30.	Liu, Z. et al. Machine learning-based integration develops an immune-derived lncRNA signature for improving outcomes in colorectal cancer. *Nat Commun* **13**, 816 (2022).

**Supplementary Figures**

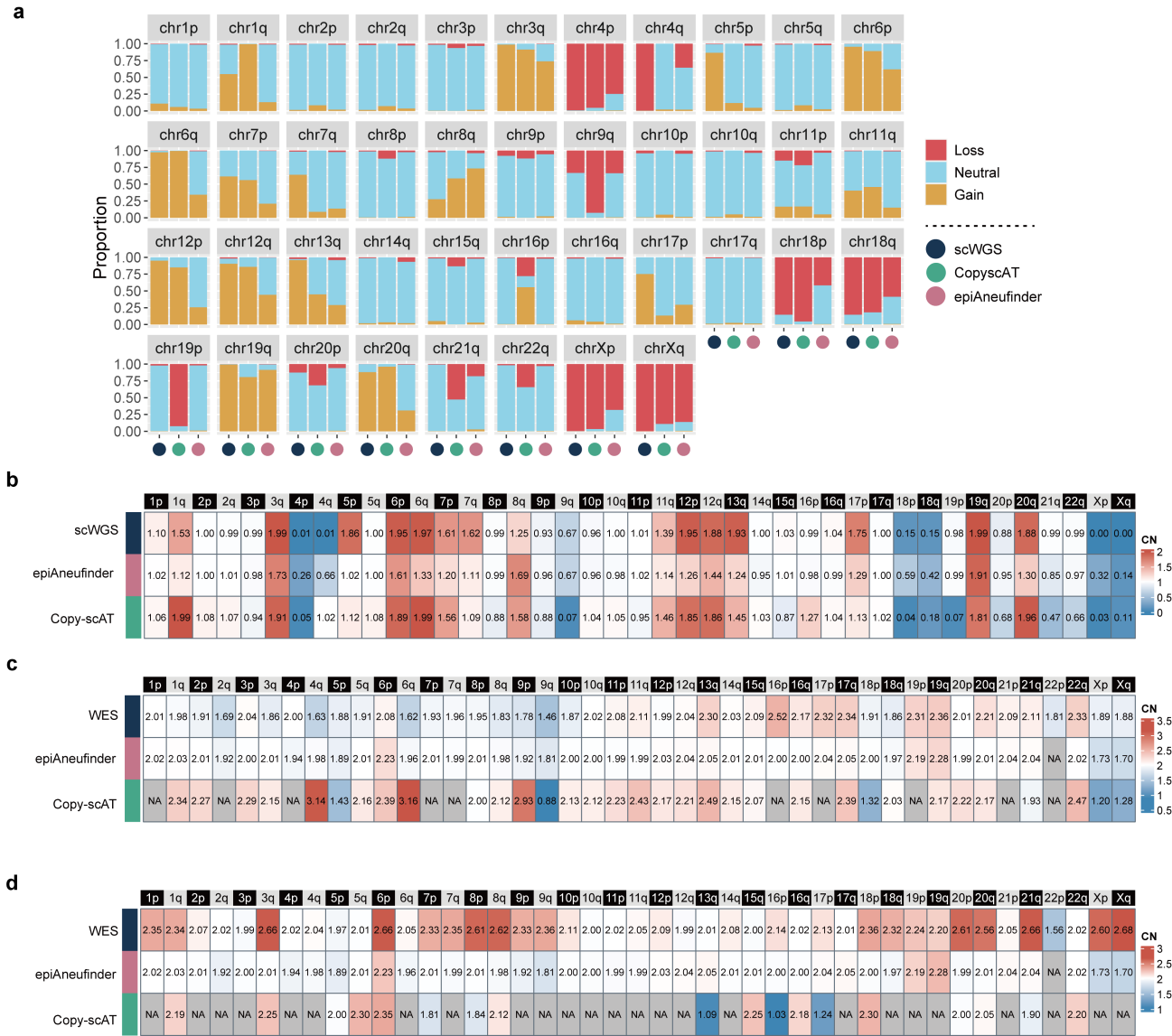

**Supplementary Fig. 1. Benchmarking CNV calling from scATAC-seq with DNA-seq data.** **a.** Cell proportions of copy number value in each chromosome measured by scATAC-seq and scWGS data in an SNU601 gastric cancer cell line. **b-d.** Consensus copy number profiles of pseudo-bulk DNA-seq and ATAC-seq in (b) SNU601, (c) SU006 and (d) SU008.

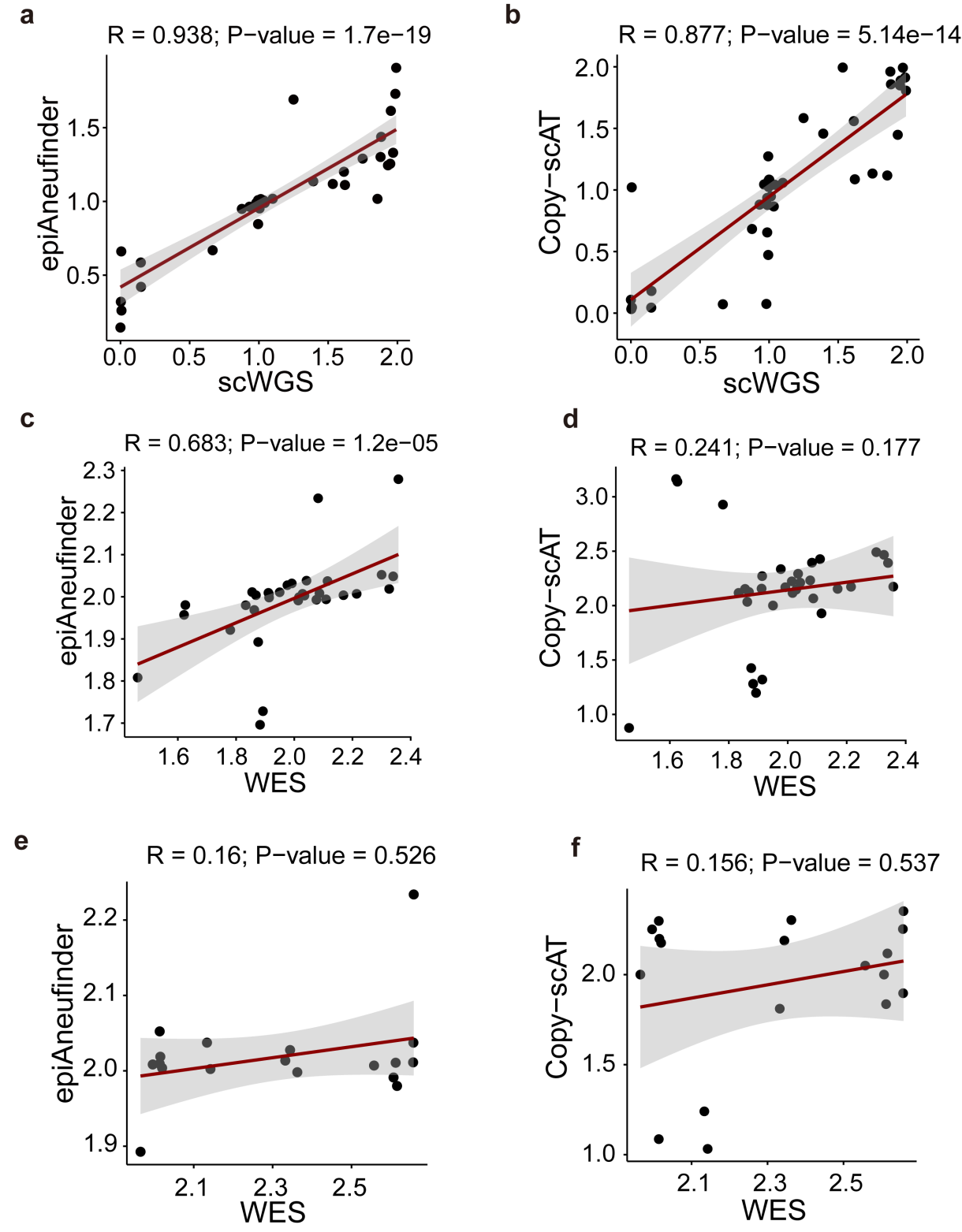

**Supplementary Fig. 2. Scatter plots showing Spearman correlation between the copy number values measured from DNA sequencing and pseudo-bulk copy numbers estimated from scATAC-seq. a-b.** Correlation between copy numbers estimated from scATAC-seq and single-cell whole genome sequencing data in sample SNU601. **c-d.** Correlation between copy numbers estimated from scATAC-seq and whole exome sequencing in sample SU006. **e-f.** Correlation between copy numbers estimated from scATAC-seq and whole exome sequencing in sample SU008. The red line indicates the fitted line, and the gray ribbons signify 95% confidence intervals.

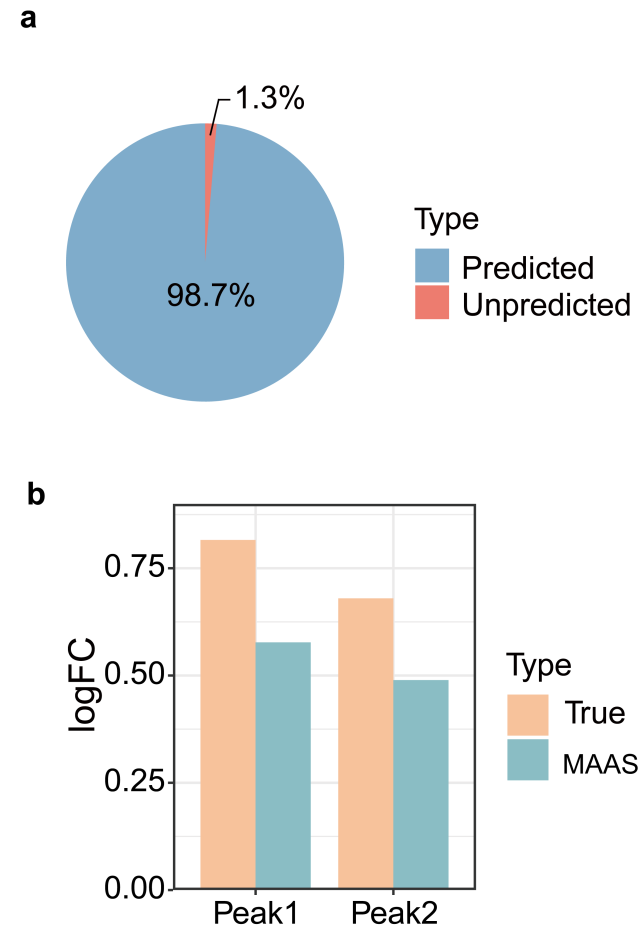

**Supplementary Fig. 3. Differential accessible chromatin regions between cluster 1 and cluster 2. a.** Proportions of predicted and unpredicted differential peaks between the subpopulations identified by MAAS. **b.** The logFC of the two unrecovered peaks.

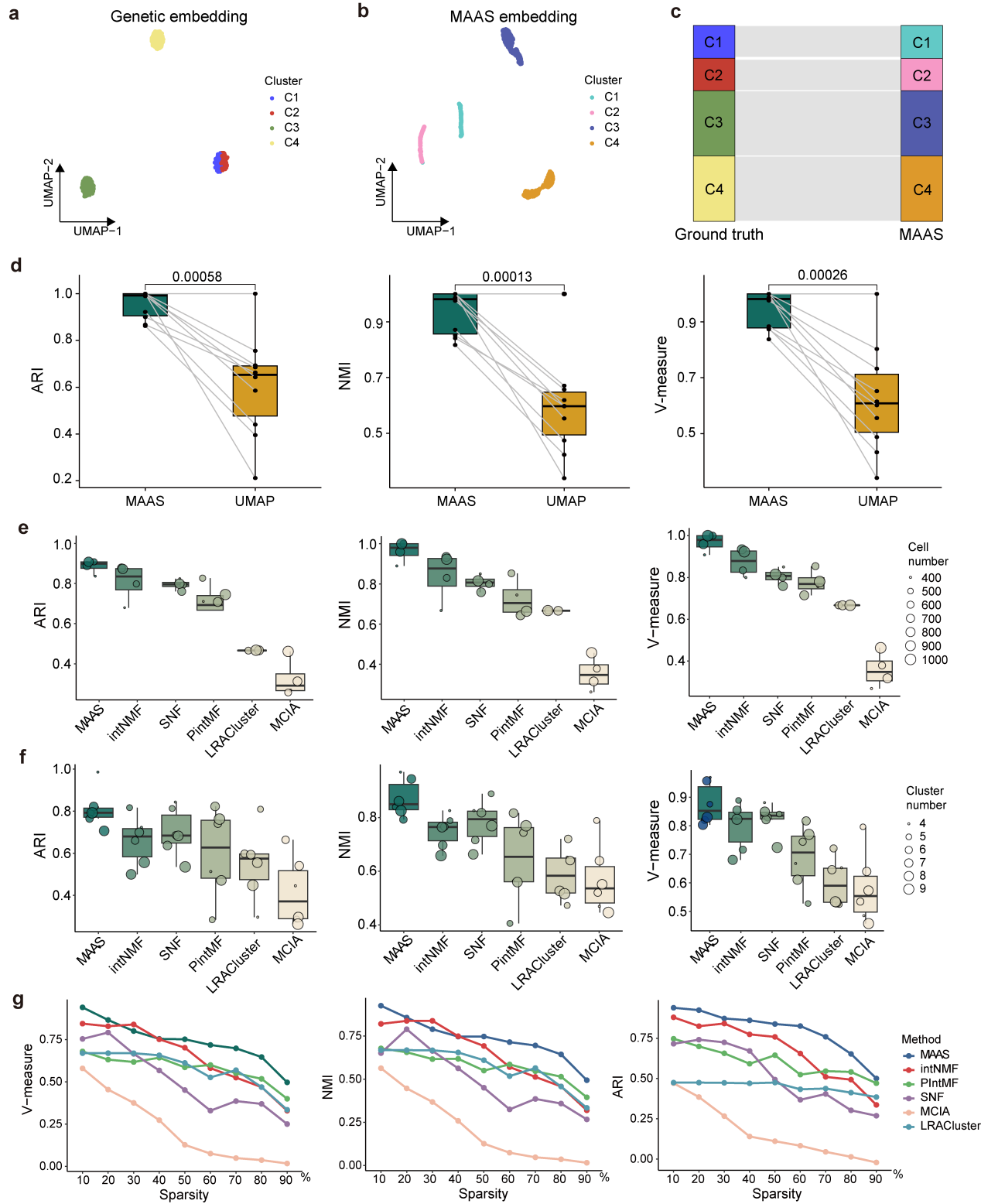

**Supplementary Fig. 4. Comparing performances of tumor subpopulation identification by MAAS and other state-of-the-art methods.** **a.** UMAP embedding based on genetic features of four subpopulations. **b.** UMAP embedding on MAAS latent factors of four subpopulations. **c.** Consistency of cell distribution across the three subpopulations when comparing ground-truth to MAAS results. **d.** Accuracy of classifying subpopulations by UMAP and MAAS, as measured by ARI (left), NMI (middle), and V-measure (right). P-value were determined by a two-tailed t-test. **e**,**f.** Accuracy of classifying varying (e) number of cells and (f) subpopulations by different methods, as measured by ARI (left), NMI (middle), and V-measure (right). The center line of the boxplot indicates the median, the box limits show the first and third quartiles, and the whiskers extend to the maximum and minimum values within 1.5 times the interquartile range from the hinge. **g.** Clustering performance across different levels of data sparsity.

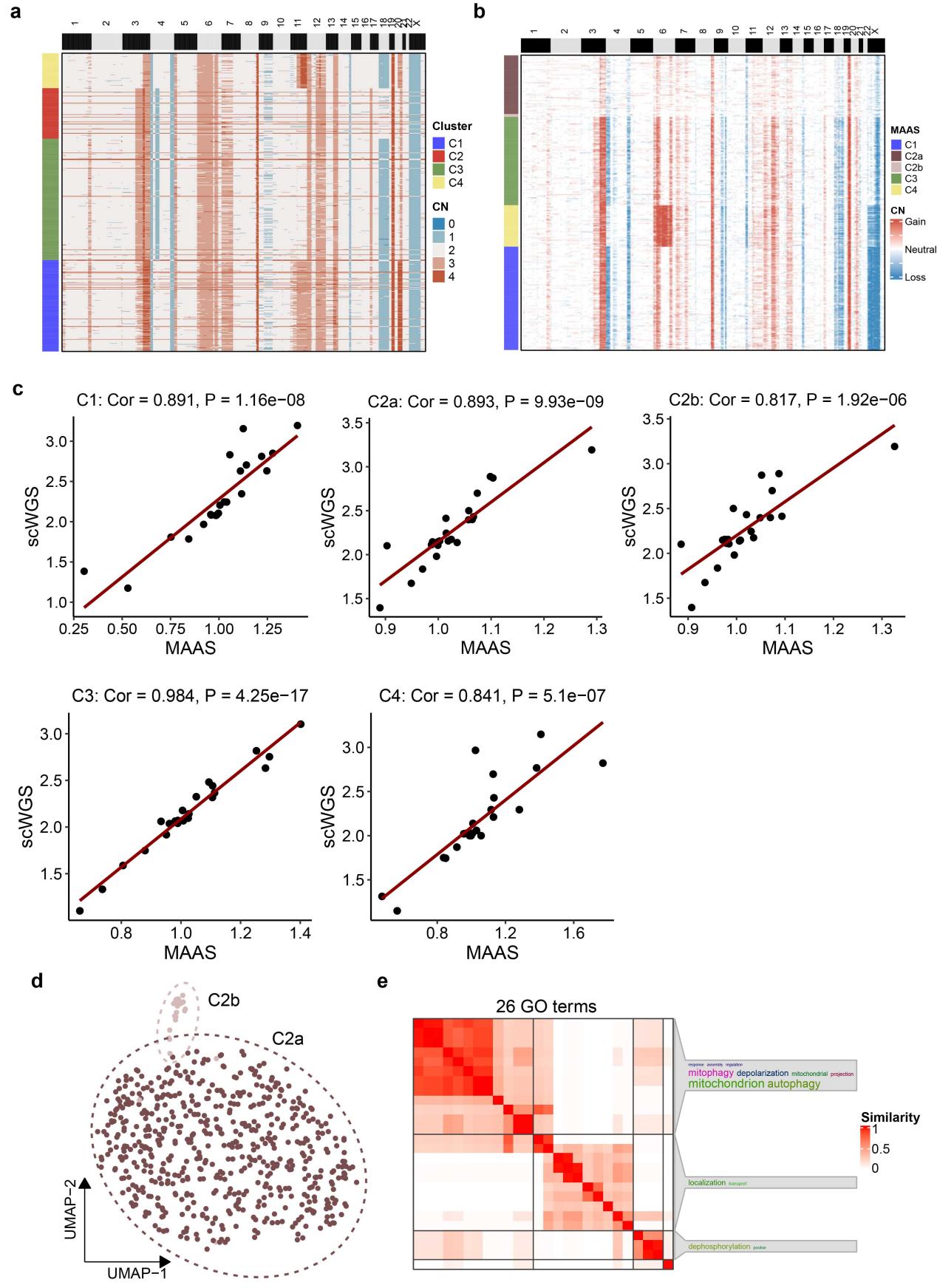

**Supplementary Fig. 5. Identification of gastric cancer subpopulations using scDNA-seq and MAAS. a-b.** Copy number profiles of subpopulations identified by scDNA-derived CNVs (a) and MAAS (b). Each row indicates a cell, and each column indicates a chromosome region (resolution = 1Mb). **c.** Pearson correlation of CNV profiles between MAAS- and scWGS-identified subpopulations. Each point represents a chromosome. The CNV values were averaged across cells. **d.** UMAP embedding of MAAS-predicted clusters of C2, where C2b is a minor subpopulation ignored by scDNA-seq. **e.** Significant biological processes from gene ontology terms (adjusted P-value < 0.05) enriched by genes near differential accessible regions (Supplementary Data 1) between the clusters C2a and C2b. The results were hierarchically clustered using the ‘binary cut’ strategy.

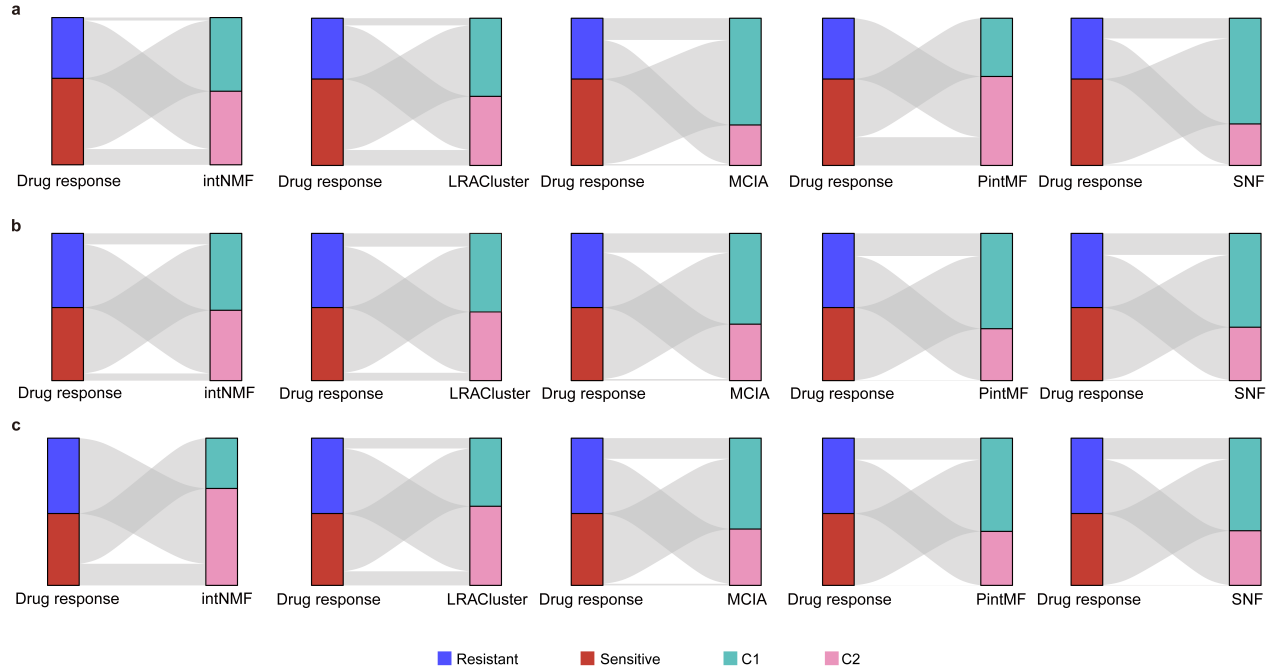

**Supplementary Fig. 6. Sankey plot illustrating the distribution of drug-sensitive and drug-resistant subpopulations compared with MAAS clusters across three different cell lines. a.** .Epidermoid carcinoma (A431). **b.** Hypopharyngeal cancer (FaDu). **c.** Ovarian cancer (SKOV3).

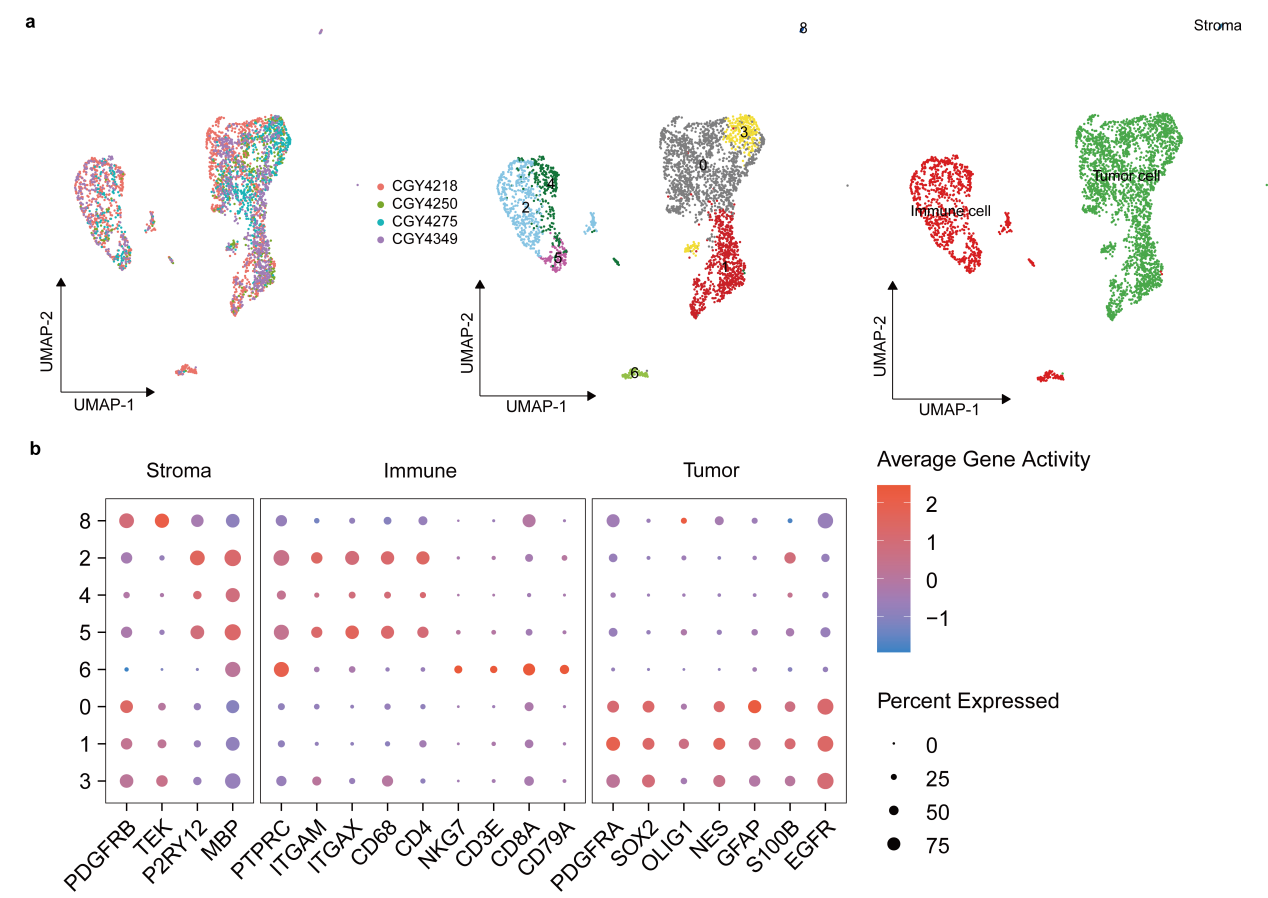

**Supplementary Fig. 7. Cell type annotation of glioma. a.** UMAP plots showing distribution of samples (left), unsupervised clusters (middle) and cell types (right). **b.** Dot plot showing the gene activity scores of cell-type marker genes in the scATAC-seq data.

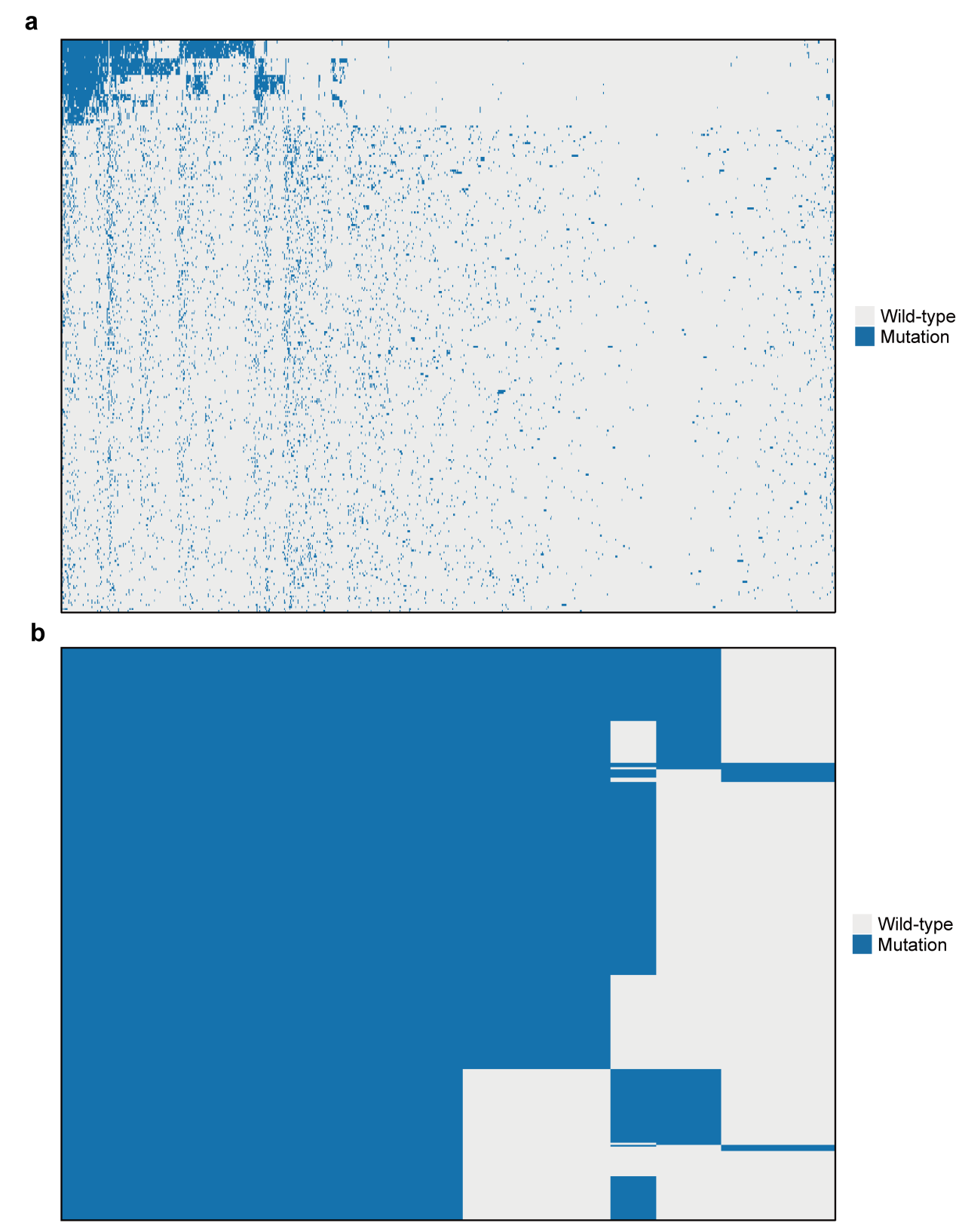

**Supplementary Fig. 8. SNV profile of tumor cells of glioma. a.** Raw SNV profile. **b.** Denoised SNV profile. Somatic SNVs in rows and cells in columns. The heatmap clustered by both rows and columns. Only reccurent mutations across samples are included. Note that since SNVs calling were failed in samples CGY4275 and CGY4349, we excluded these two samples for further analysis.

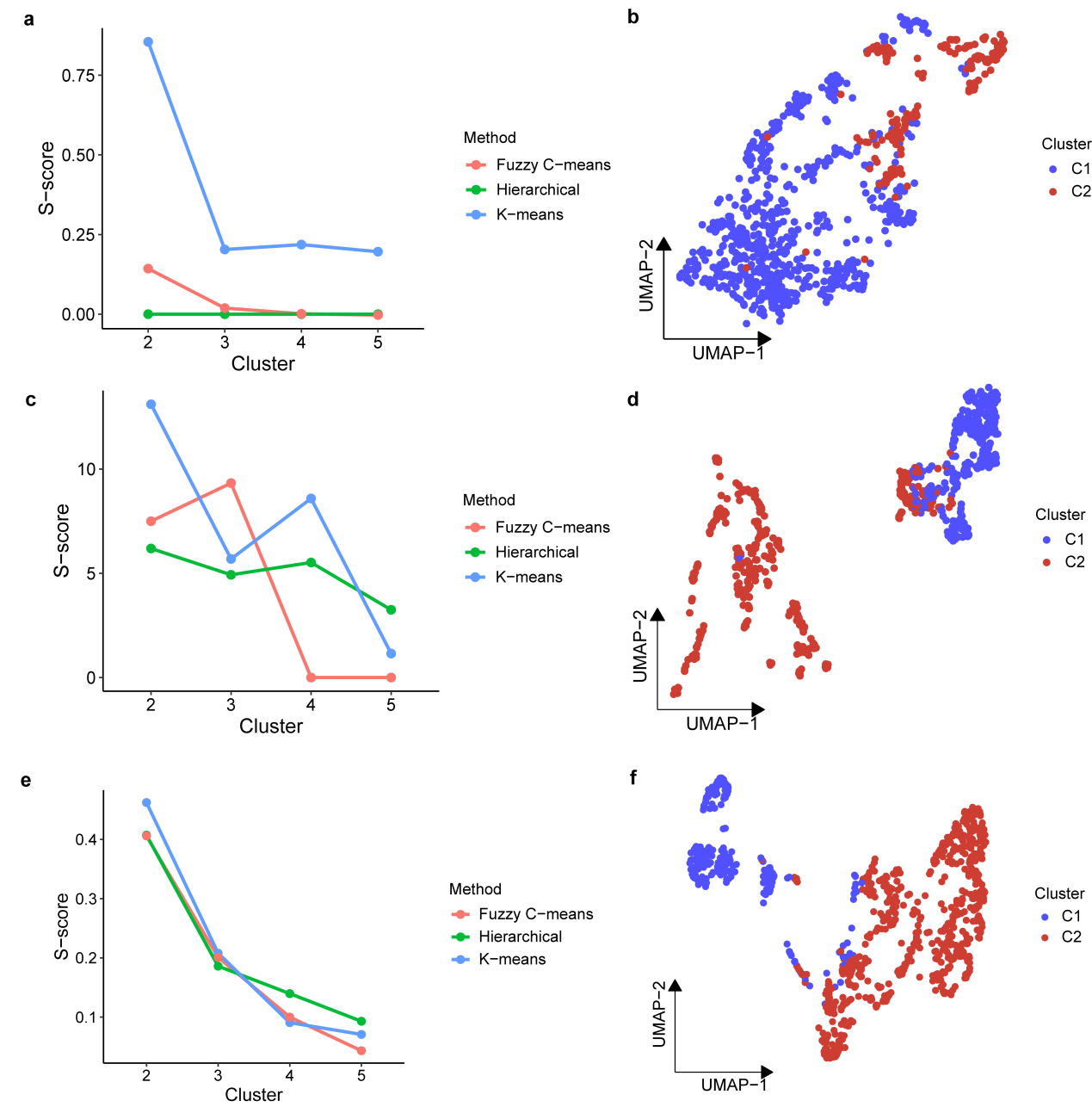

**Supplementary Fig. 9. Tumor cell clusters defined by CNVs and chromatin states of glioma. a,b.** Clustering results based on CNVs by epiAneufinder. **c,d.** Clustering results based on CNVs by Copy-scAT. **e,f.** Clustering results based on chromatin accessibility. The (a,c and e) left panel showing the S-score across different clustering numbers. The (b,d and e) right panel showing the UMAP plots of tumor cells substructures.

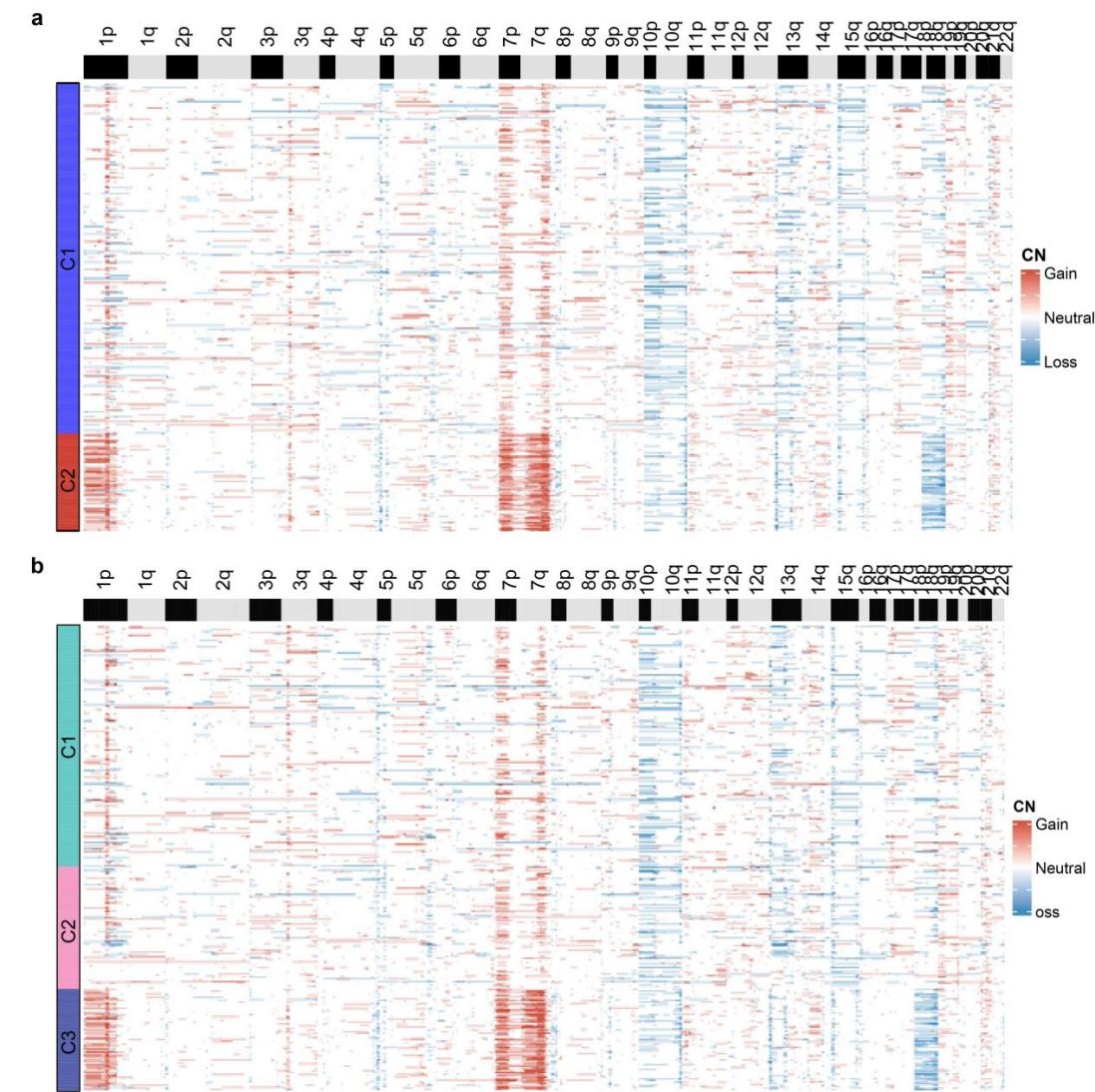

**Supplementary Fig. 10. Copy number profiles of the glioma tumor cell clusters.** **a.** Cell clusters identified by epiAneufinder. **b.** Cell clusters identified by MAAS. Each column indicates each bin (100,000 bp), and each row represents a cell. Blue indicates copy number loss; white indicates copy number neutral; red indicates copy number gain.

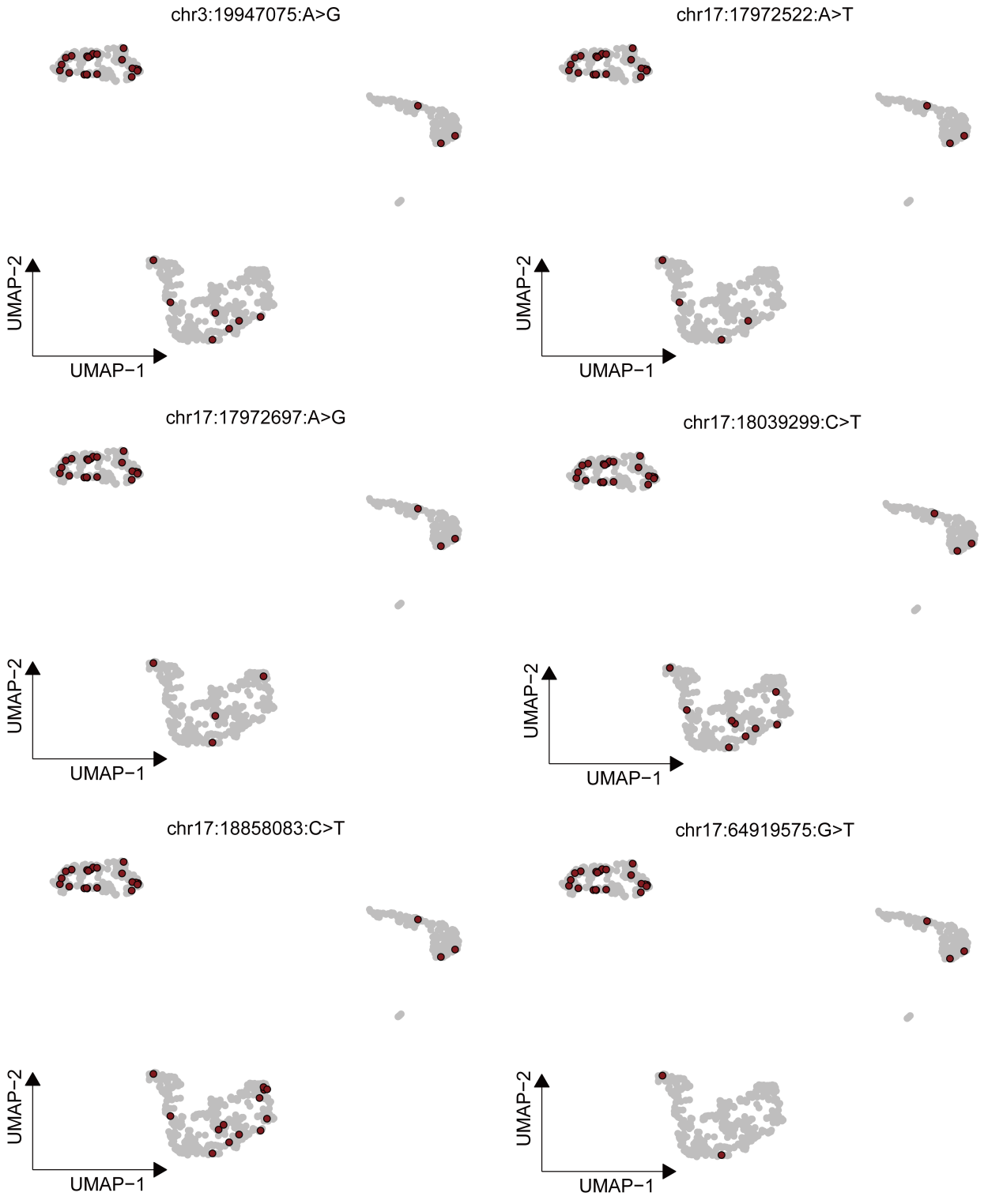

**Supplementary Fig. 11. UMAP plots showing the six cluster 1-specific SNVs (upper left of each UMAP plot), which are highlighted in dark red dots.**

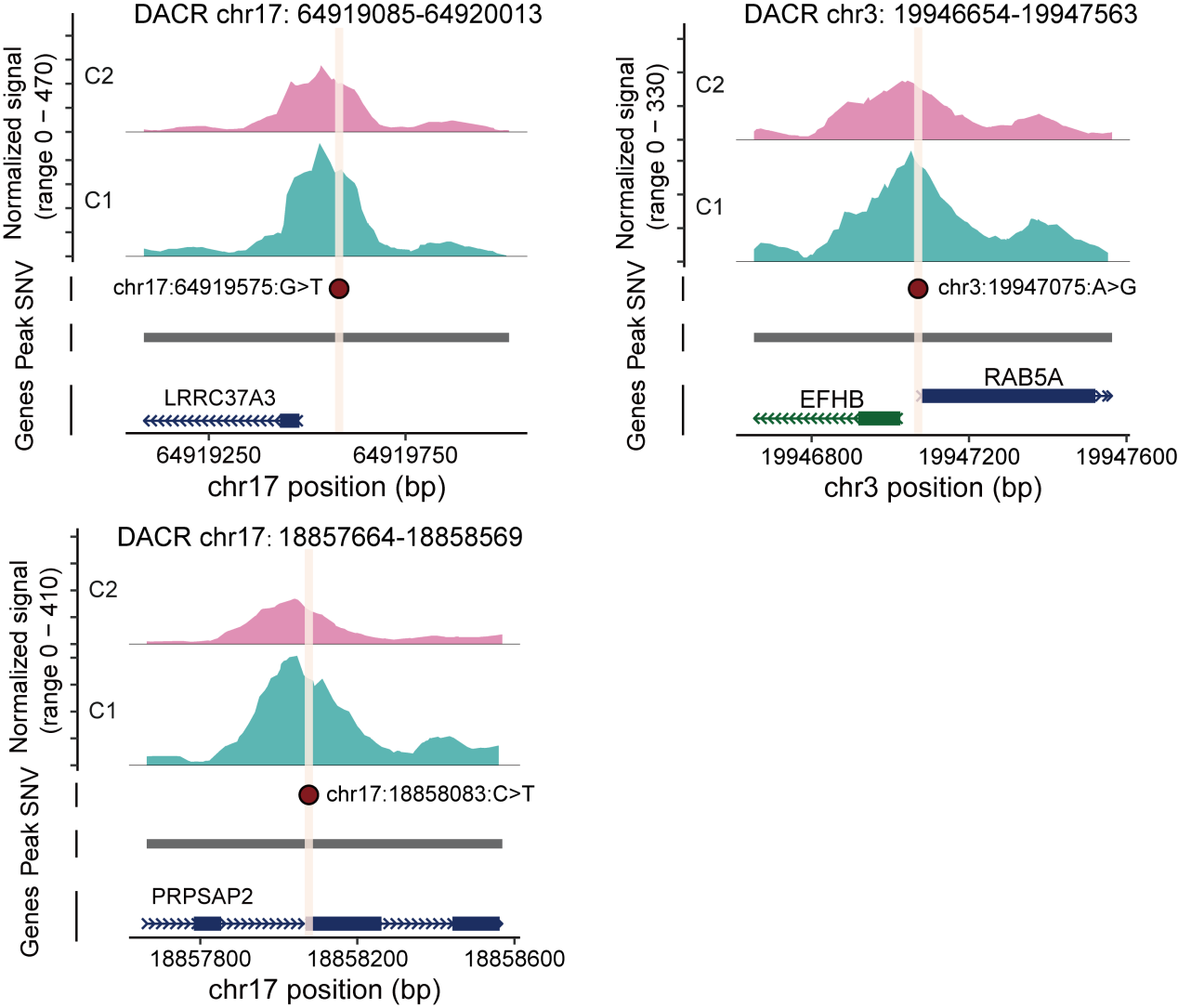

**Supplementary Fig. 12. scATAC-seq (top) peak tracks of clusters 1 and 2 for accessible regions chr17:64919085-64920013 (left), chr17:18857664-18858569 (bottom) and chr3:19946654-19947563 (right), with noncoding SNVs marked by dark red dots.**

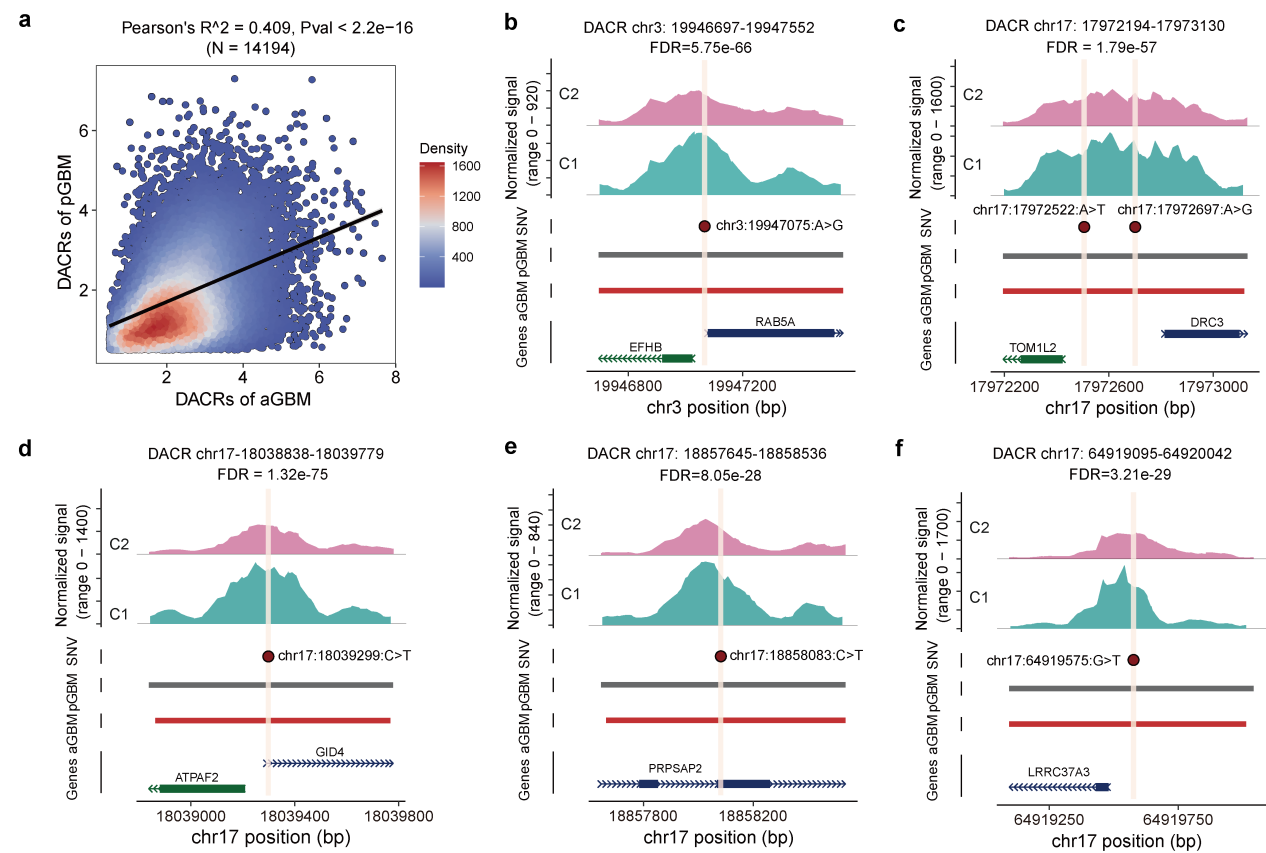

**Supplementary Fig. 13. Differential accessible chromatin regions (DACRs) between adult GBM clusters 1 and 2 validated on a pediatric GBM dataset.** **a.** Correlation of log fold change of DACRs between adult and pediatric GBM. **b-f.** scATAC-seq (top) peak tracks of clusters 1 and 2 for the five accessible chromatin regions containing driving SNVs.

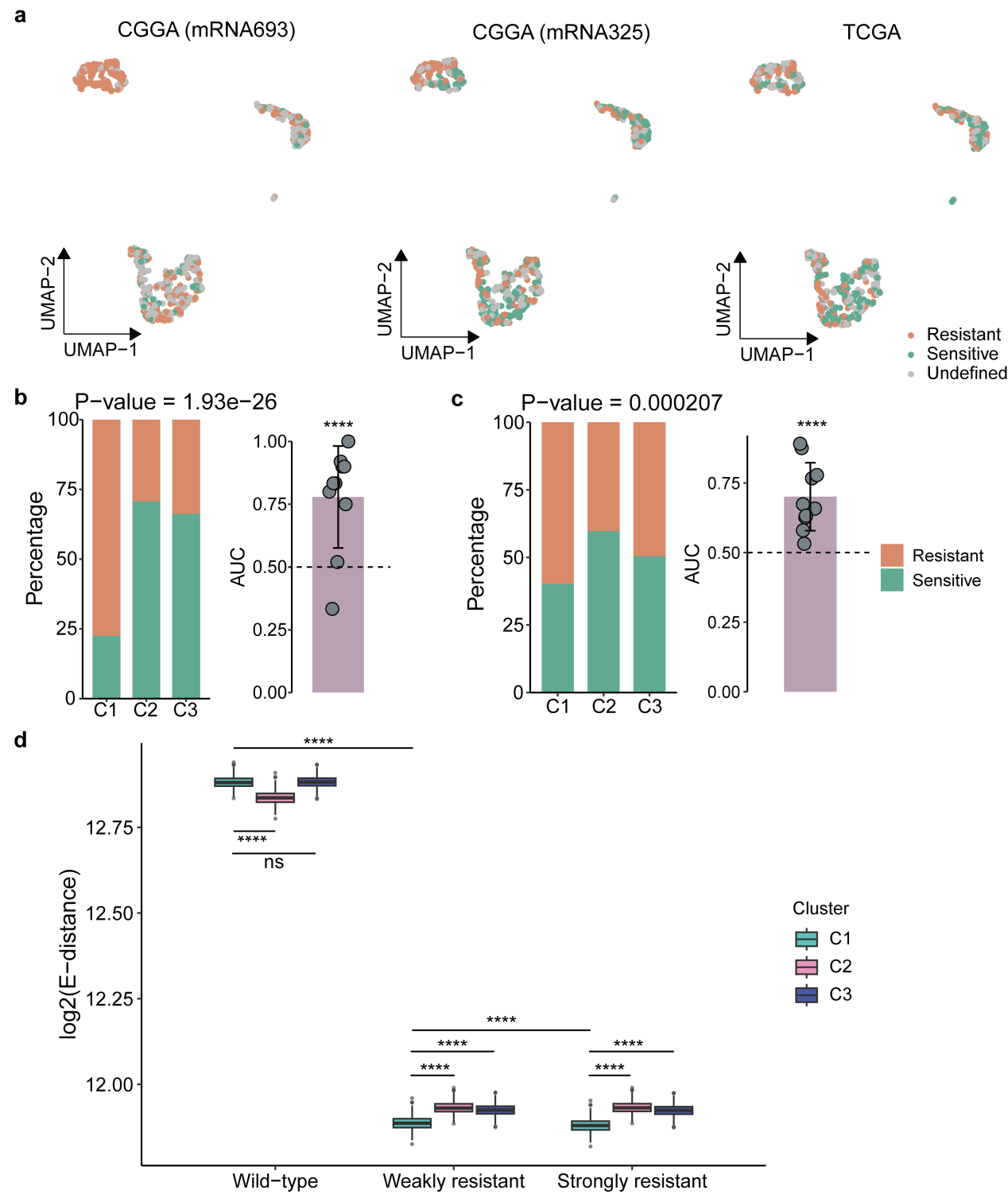

**Supplementary Fig. 14. Responses of MAAS-identified clusters temozolomide (TMZ). a.** UMAP plots showing TMZ-sensitive and resistant cells predicted by Scissor based on three different bulk-seq datasets. **b,c.** Distribution of predicted temozolomide (TMZ)-sensitive and resistant cells across MAAS-identified clusters based on the references of (b) CGGA-mRNA325 and (c) The Cancer Genome Atlas (TCGA) cohorts. The P-value was determined by the chi-square test. The right panel shows the accuracy of predictions measured by area under the curve (AUC). The P-value, calculated by a chi-square test, is shown, along with the accuracy of predictions measured by the AUC. The dotted line represents the baseline of 0.5. Asterisks indicate significance based on permutations. **d.** Energy distance between the three MAAS clusters and wild-type, weakly resistant as well as strongly resistant glioma tumor cells. The center line of the boxplot indicates the median, the box limits show the first and third quartiles, and the whiskers extend to the maximum and minimum values within 1.5 times the interquartile range from the hinge. The P-value was determined by a two-tailed Wilcoxon rank-sum test; **** indicates P-value < 0.0001. ns: no significance

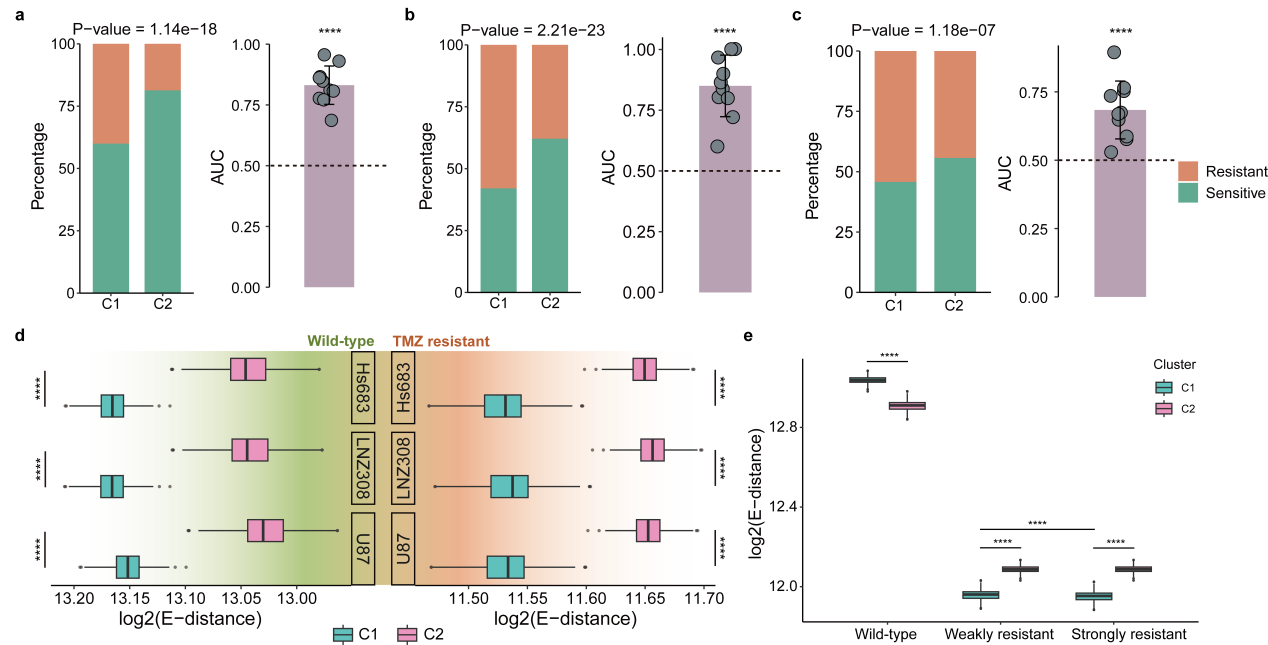

**Supplementary Fig. 15. Responses of MAAS-identified clusters temozolomide (TMZ) in the validation dataset. a-c.** Distribution of predicted temozolomide (TMZ)-sensitive and resistant cells across MAAS-determined clusters by leveraging bulk-seq information from the (a) CGGA-mRNA693, (b) CGGA-mRNA325 and (c) The Cancer Genome Atlas (TCGA) cohorts. The P-value, calculated by a chi-square test, is shown, along with the accuracy of predictions measured by the area under the curve (AUC). The dotted line represents the baseline of 0.5. Asterisks indicate significance based on permutations. **d,e.** Energy distance (E-distance) between the (d) three MAAS clusters and TMZ sensitive and resistant cells across three cell lines, (e) as well as E-distance between MAAS clusters and wild-type, weakly resistant as well as strongly resistant glioma tumor cells. The center line of the boxplot indicates the median, the box limits show the first and third quartiles, and the whiskers extend to the maximum and minimum values within 1.5 times the interquartile range from the hinge. The P-value was determined by a two-tailed Wilcoxon rank-sum test; **** indicates P-value < 0.0001.

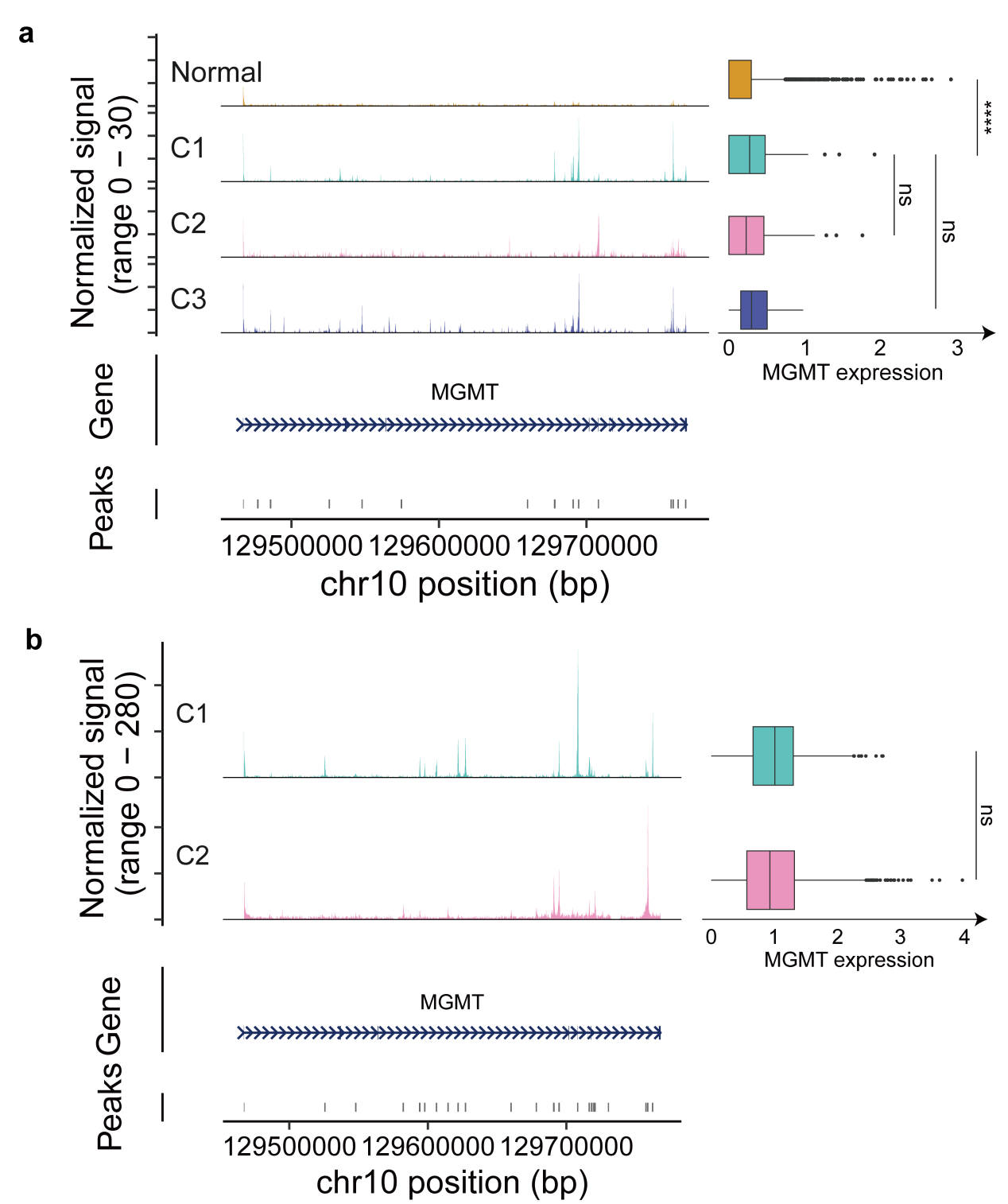

**Supplementary Fig. 16. The TMZ-resisatance of MAAS clusters was independent of *MGMT* status.** scATAC-seq (left top) peak tracks of *MGMT* and gene activity (right) in (a) adult and (b) pediatric GBM. The center line of the boxplot indicates the median, the box limits show the first and third quartiles, and the whiskers extend to the maximum and minimum values within 1.5 times the interquartile range from the hinge. ns, no significance.

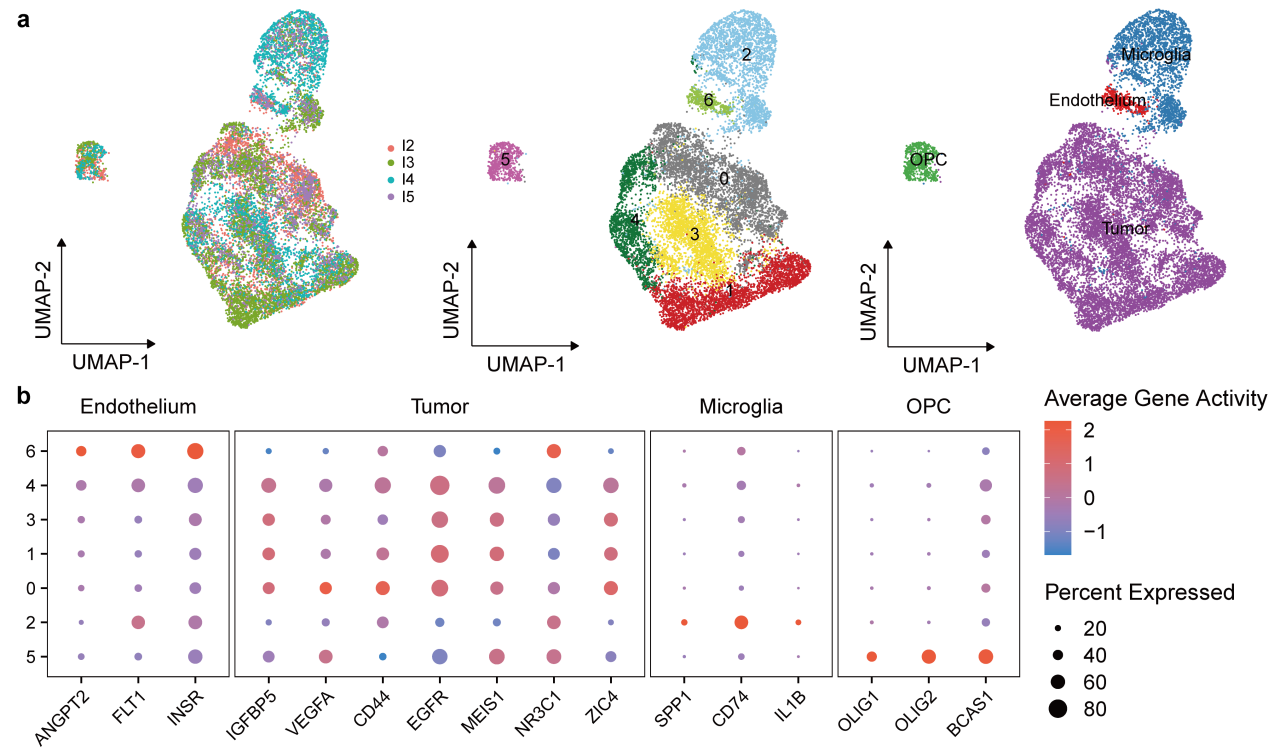

**Supplementary Fig. 17. Cell type annotation of primary pediatric posterior fossa ependymoma. a.** UMAP plots showing distribution of samples (left), unsupervised clusters (middle) and cell types (right). **b.** Dot plot showing the gene activity scores of cell-type marker genes in the scATAC-seq data.

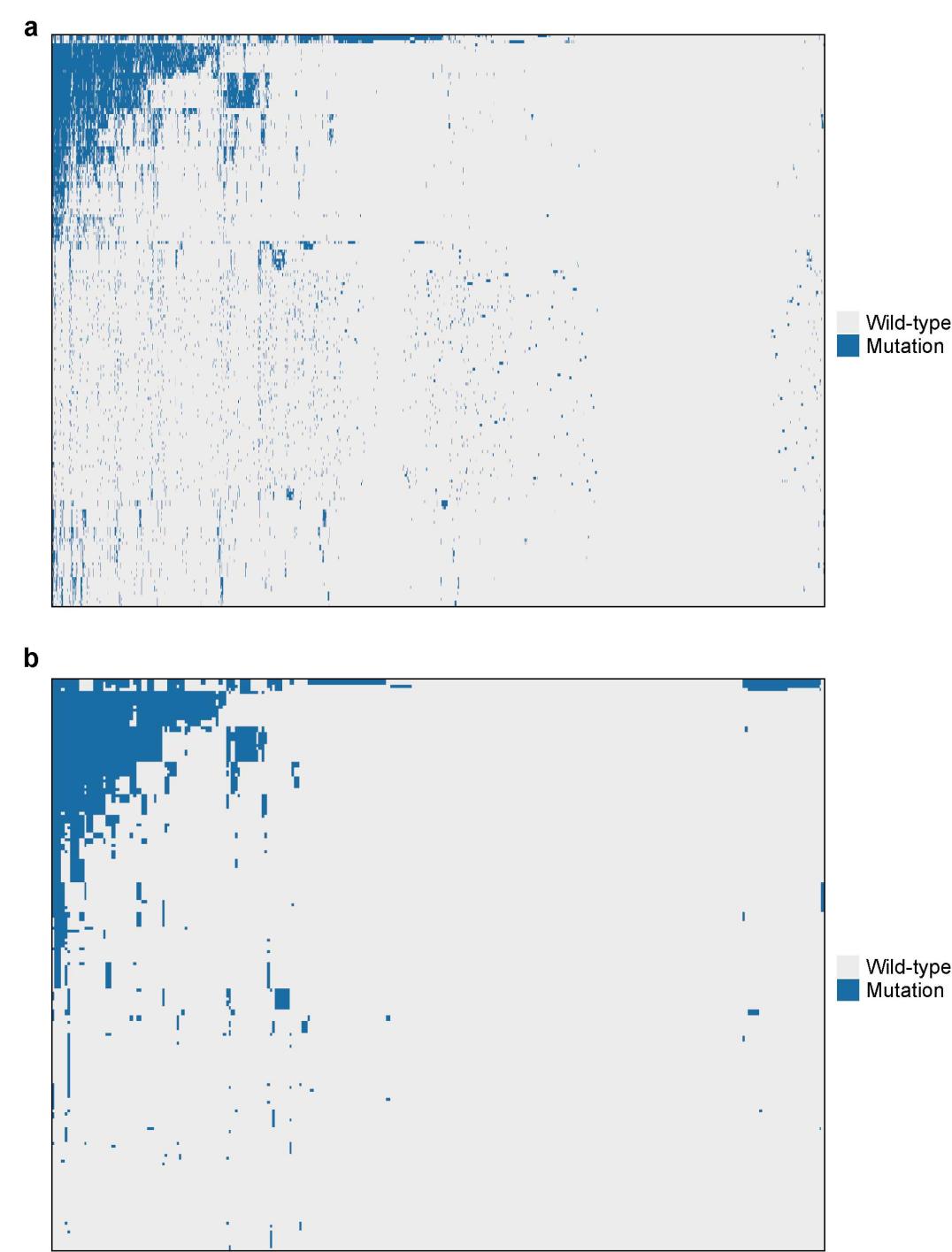

**Supplementary Fig. 18. SNV profile of tumor cells of primary pediatric posterior fossa ependymoma. a.** Raw SNV profile. **b.** Denoised SNV profile. Somatic SNVs in rows and cells in columns. The heatmap clustered by both rows and columns. Only reccurent mutations across samples are included. Note that since SNVs calling were failed in samples I3 and I5, we excluded these two samples for further analysis.

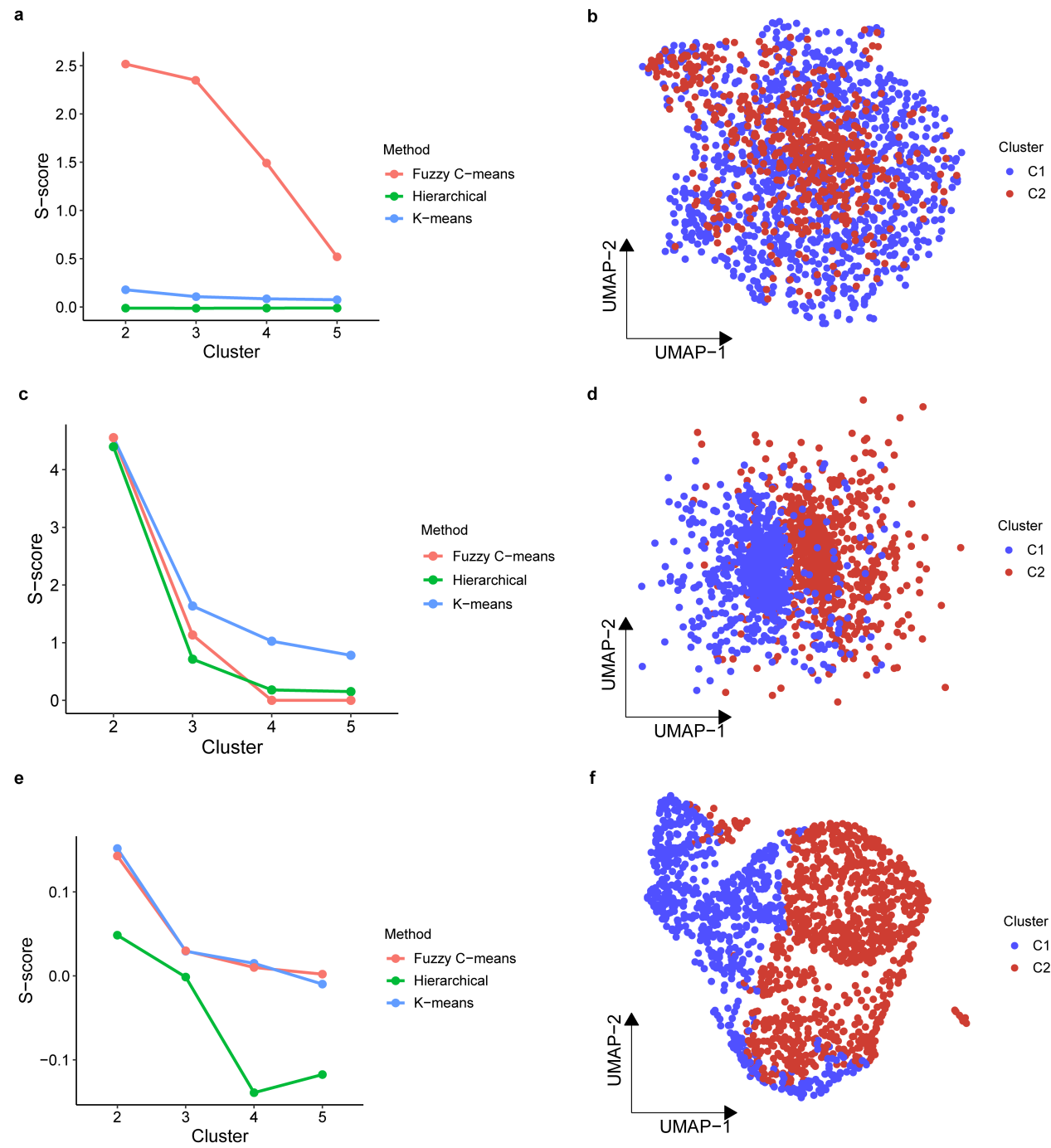

**Supplementary Fig. 19. Tumor cell clusters defined by CNVs and chromatin states of pediatric ependymoma. a,b.** Clustering results based on CNVs by epiAneufinder. **c,d.** Clustering results based on CNVs by Copy-scAT. **e,f.** Clustering results based on chromatin accessibility. The (a,c and e) left panel showing the S-score across different clustering numbers. The (b,d and e) right panel showing the UMAP plots of tumor cells substructures.

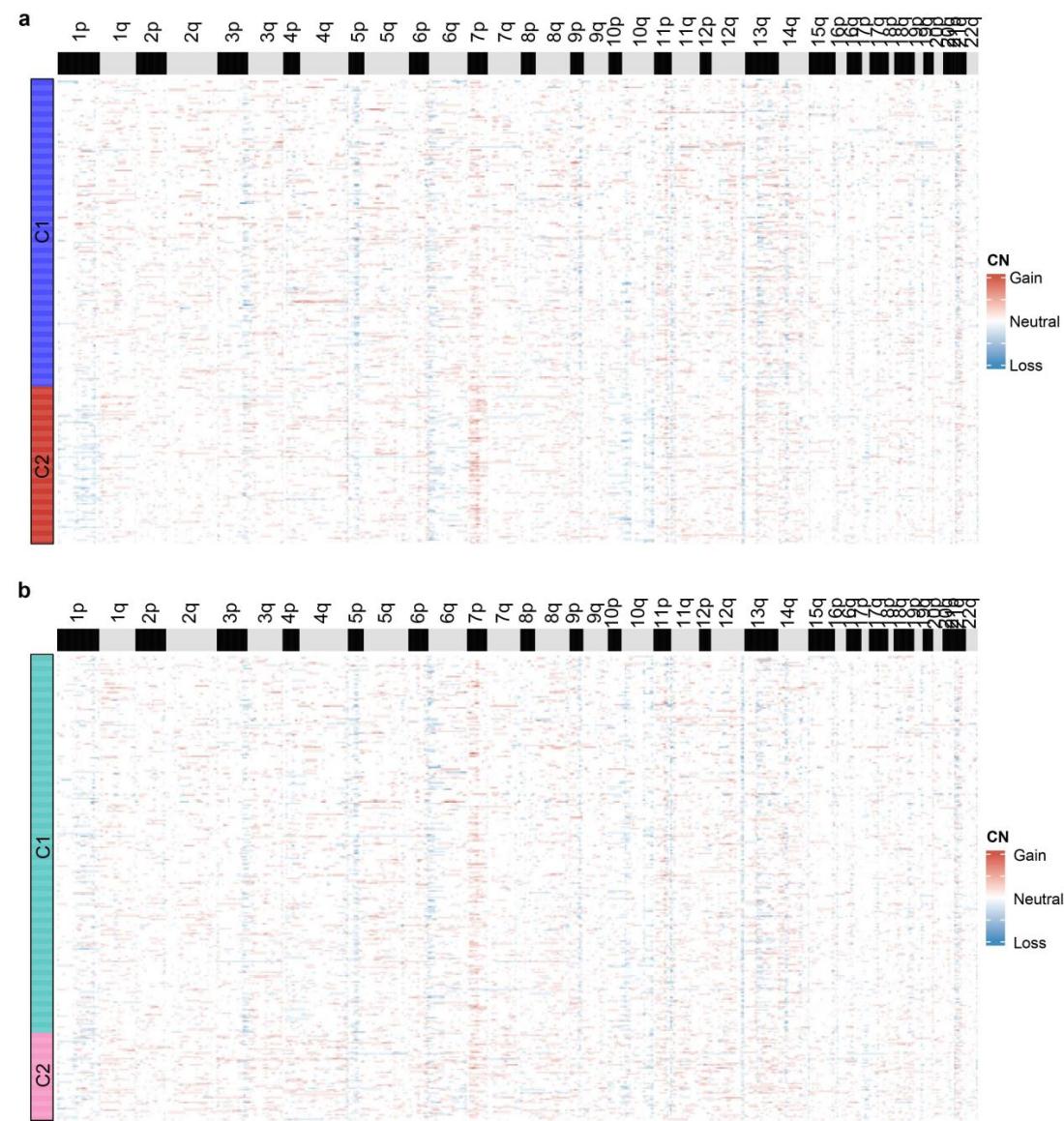

**Supplementary Fig. 20. Copy number profiles of the pediatric ependymoma cell clusters.** **a.** Cell clusters identified by epiAneufinder. **b.** Cell clusters identified by MAAS. Each column indicates each bin (100,000 bp), and each row represents a cell. Blue indicates copy number loss; white indicates copy number neutral; red indicates copy number gain.

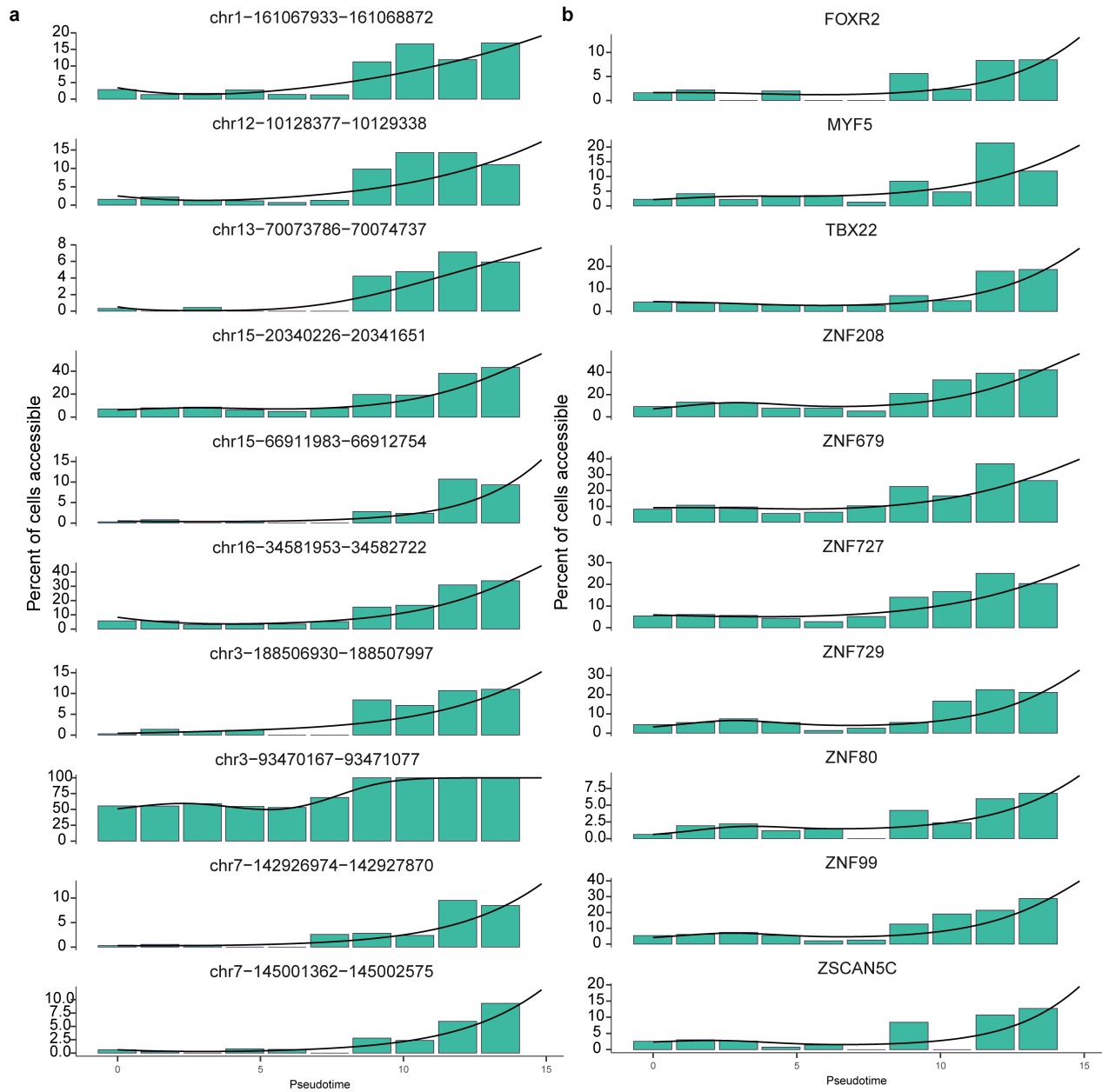

**Supplementary Fig. 21. Histogram showing the percentage of cells that had top-10 differential (a) accessible chromatin regions and (b) transcription factors calculated by aggregating chromatin signals into bins (bins=10) along pseudotime with smoothed curve.**

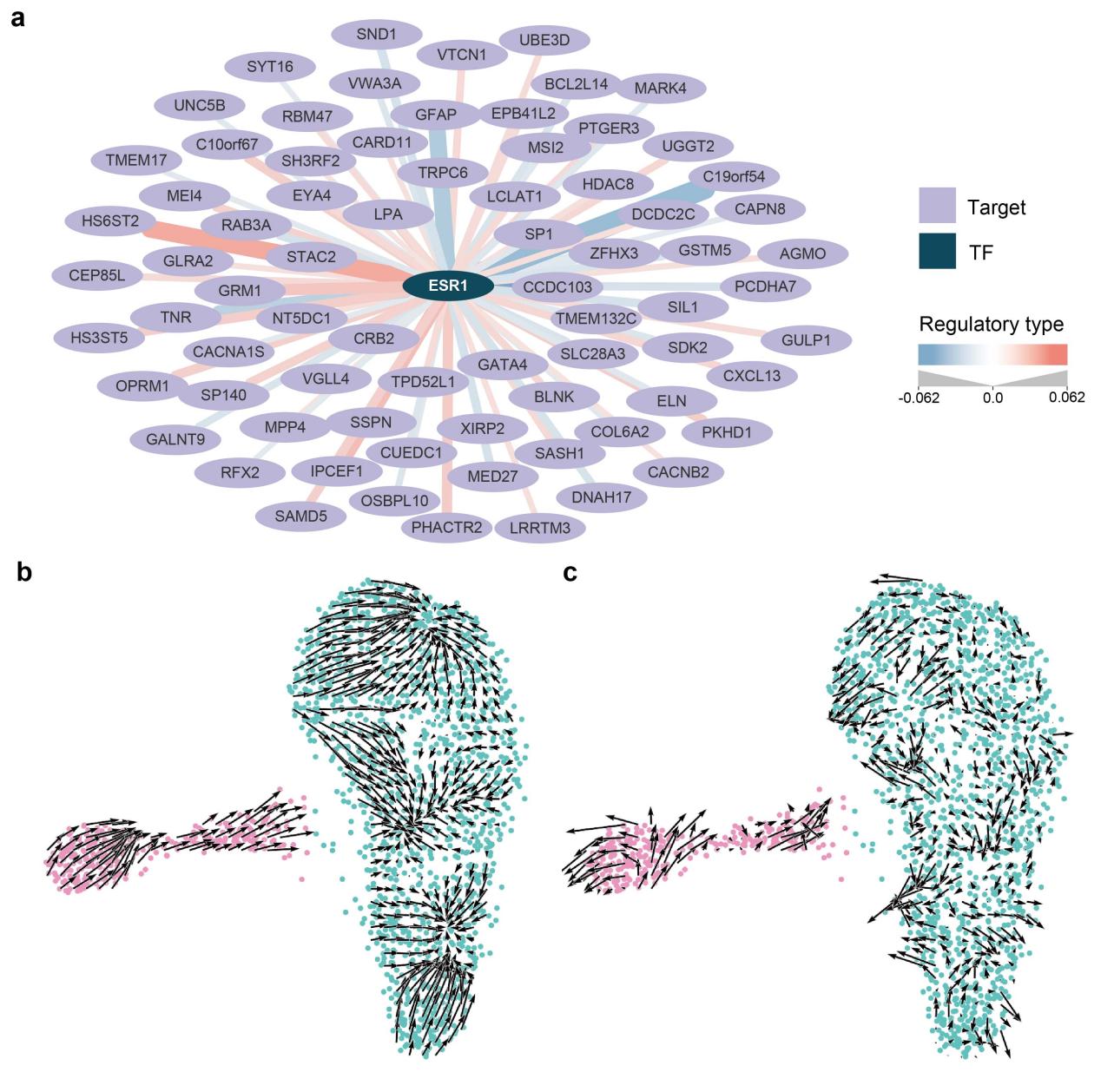

**Supplementary Fig. 22. *In silico* perturbation of *ESR1*. a.** Gene regulatory network (GRN) of *ESR1*. Only TF-target pairs with absolute value of coefficients > 0.01 are shown. **b,c.** Cell fate transition simulations performed with (b) *ESR1* knockout and (c) randomized GRNs.

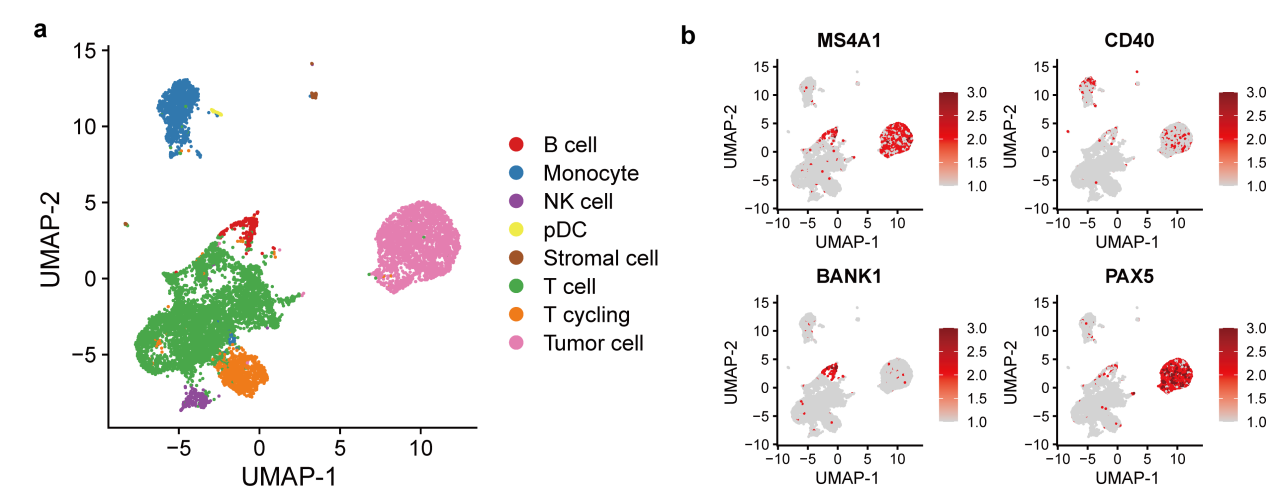

**Supplementary Fig. 23. Cell type annotation of glioma. a.** UMAP plots showing distribution of cell types. **b.** Marker genes of malignant B cells.

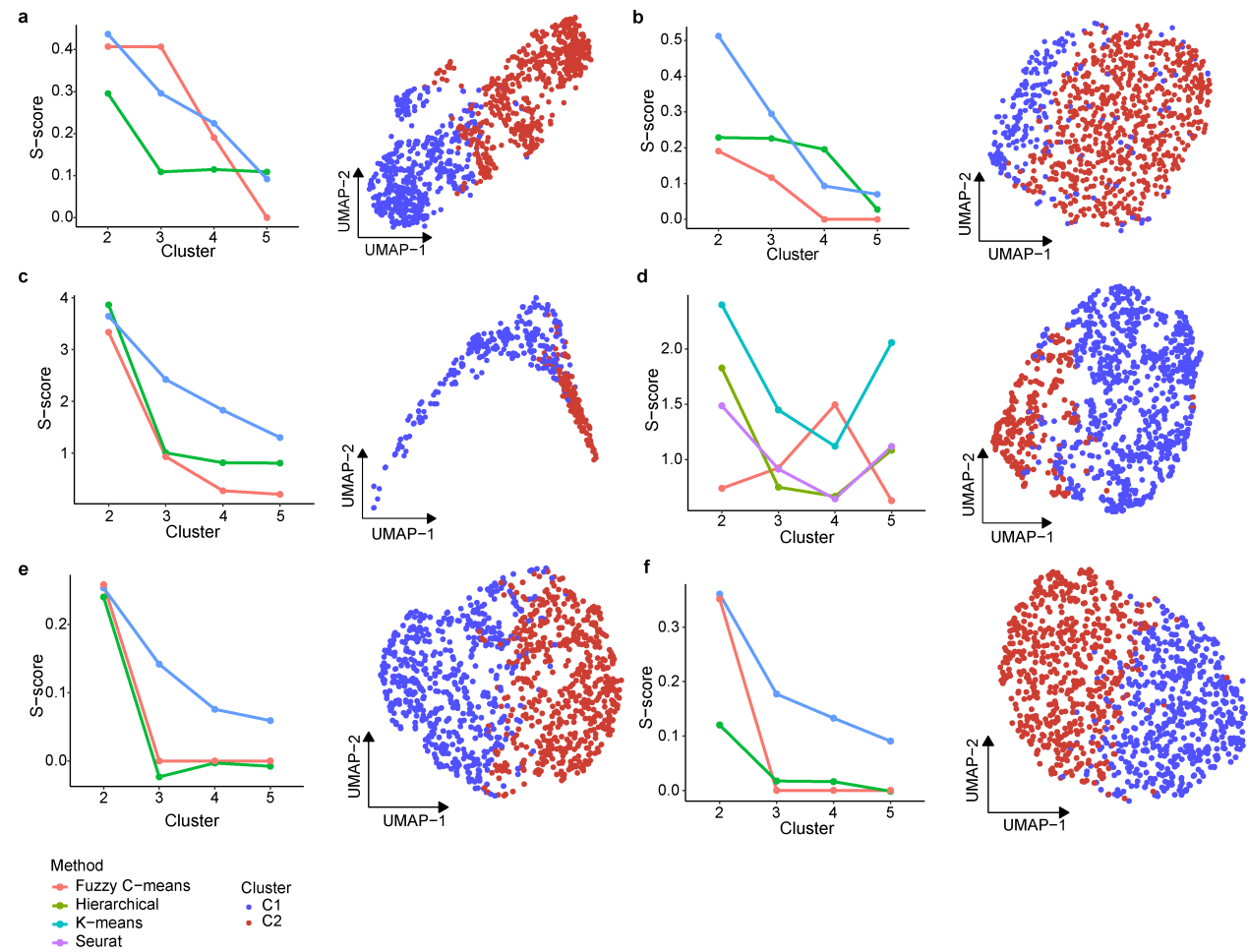

**Supplementary Fig. 24.** Tumor cell clusters identified by traditional single-modality methods by (a) epiAneufinder, (b) Copy-scAT, (c) chromatin accessibility, (d) gene expression, (e) inferCNV and (f) CopyKAT.

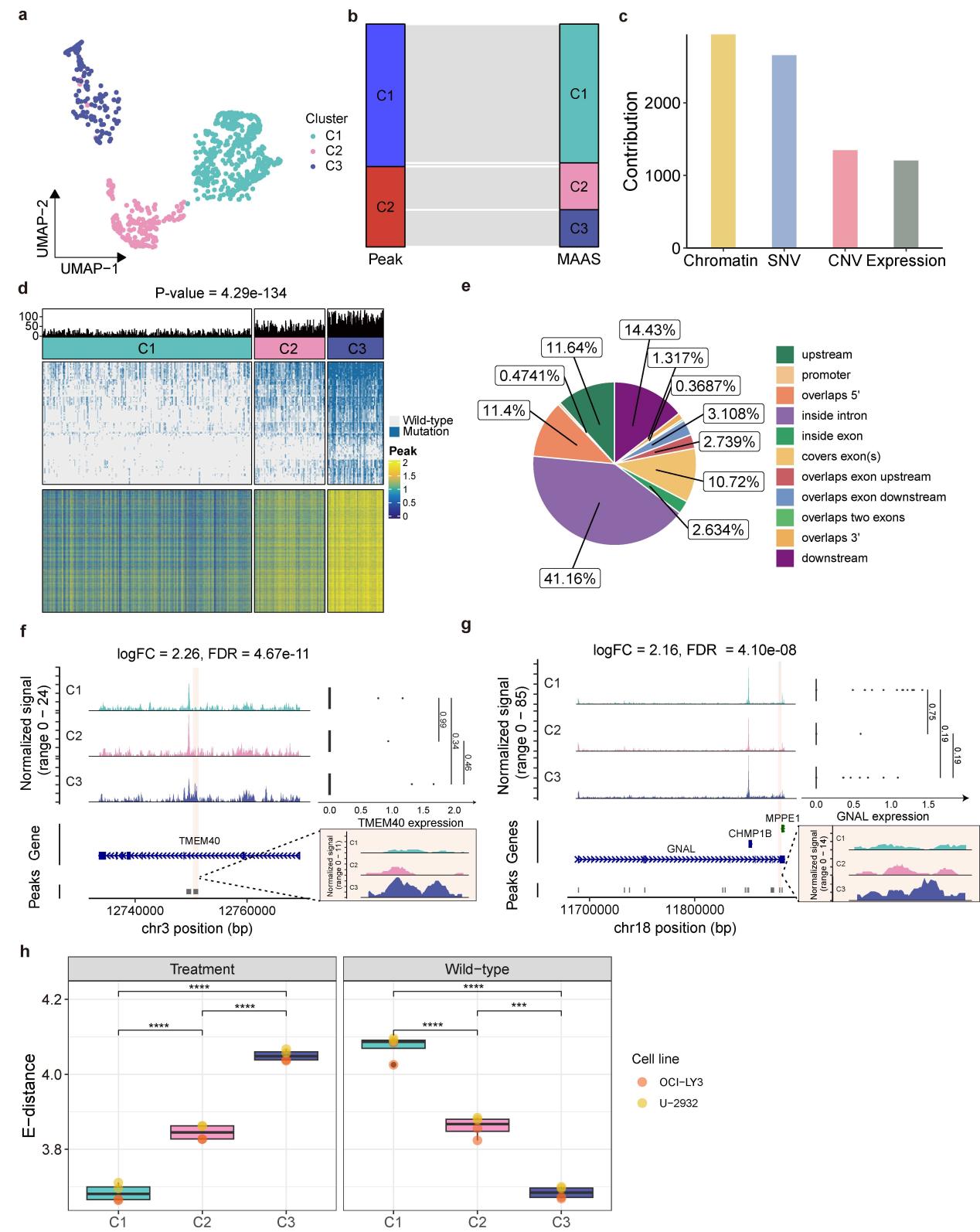

**Supplementary Fig. 25. MAAS identified new B-cell lymphoma cell subpopulations. a.** UMAP embedding of the three tumor cell subpopulations determined by MAAS. **b.** Sankey plot showed the correspondence between clusters identified by chromatin accessibility and those identified by MAAS. **c.** Contribution of each modality to the identification of subpopulations. **d.** Distribution of SNVs and top 1,000 DACRs across the clusters. The P-value for differences in mutational frequency was determined by the Kruskal-Wallis test. **e.** Genomic annotations of DACRs. **f,g.** Chromatin accessibility difference and corresponding gene expression of (f) *TMEM40* and (g) *GNAL* across three clusters. **h.** E-distance of ATAC-seq profile between MAAS-identified clusters and different treatment groups, including wild-type and TAK-981 treated cells on two B-cell lymphoma cell lines (OCI-LY3 and U-2932). The center line of the boxplot indicates the median, the box limits show the first and third quartiles, and the whiskers extend to the maximum and minimum values within 1.5 times the interquartile range from the hinge. P-values were calculated by student’s t-test.

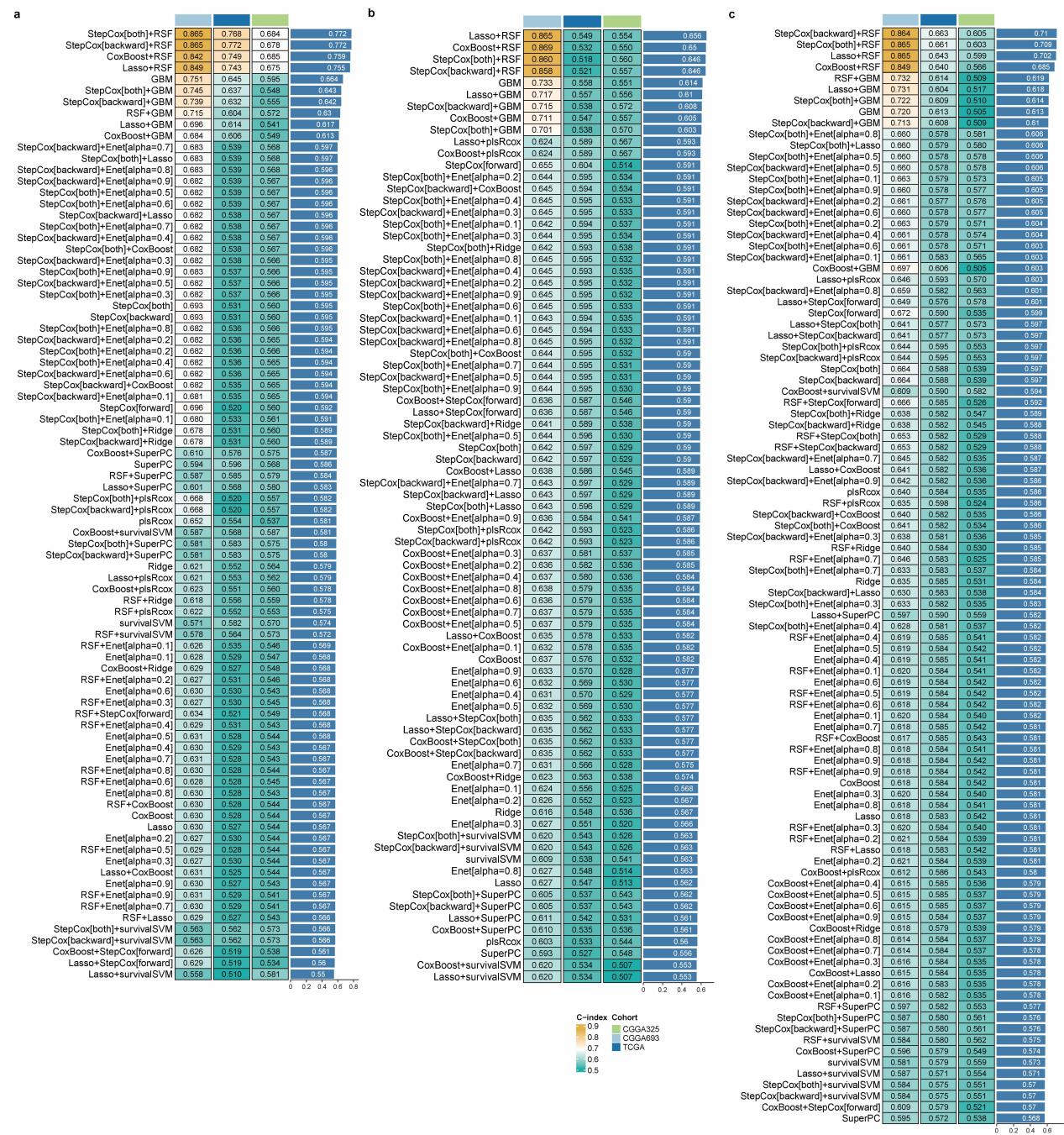

**Supplementary Fig. 26. MAASig performance in the glioblastoma datasets.** C-index of different machine learning models trained on (a) combined TF expression and activity, and single-modality features including (b) expression and (c) activity. Models with less than five genes were excluded for analysis.

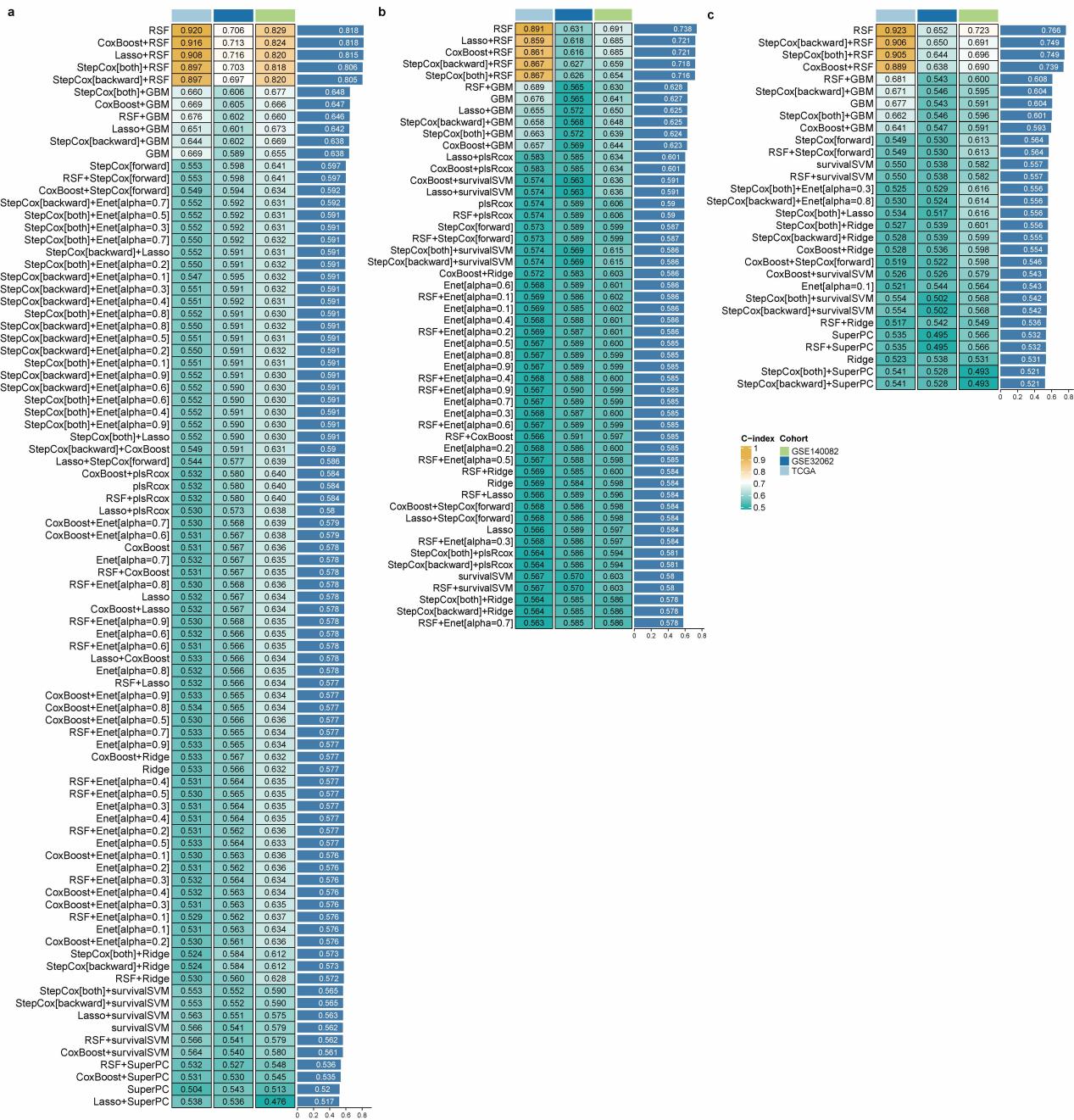

**Supplementary Fig. 27. MAASig performance in the ovarian cancer datasets.** C-index of different machine learning models trained on (a) combined TF expression and activity, and single-modality features including (b) expression and (c) activity. Models with less than five genes were excluded for analysis.

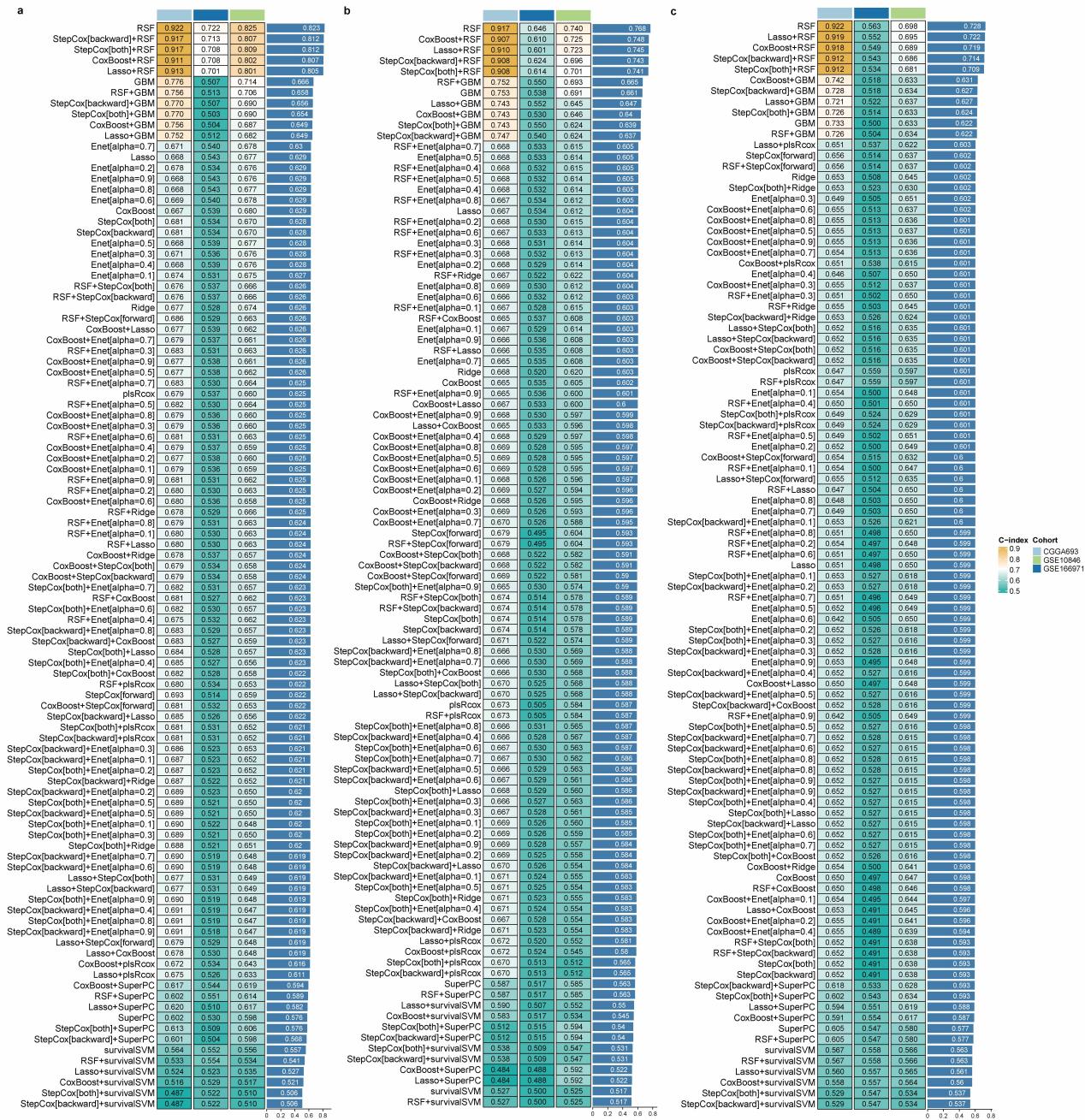

**Supplementary Fig. 28. MAASig performance in the B-cell lymphoma datasets.** C-index of different machine learning models trained on (a) combined TF expression and activity, and single-modality features including (b) expression and (c) activity. Models with less than five genes were excluded for analysis.

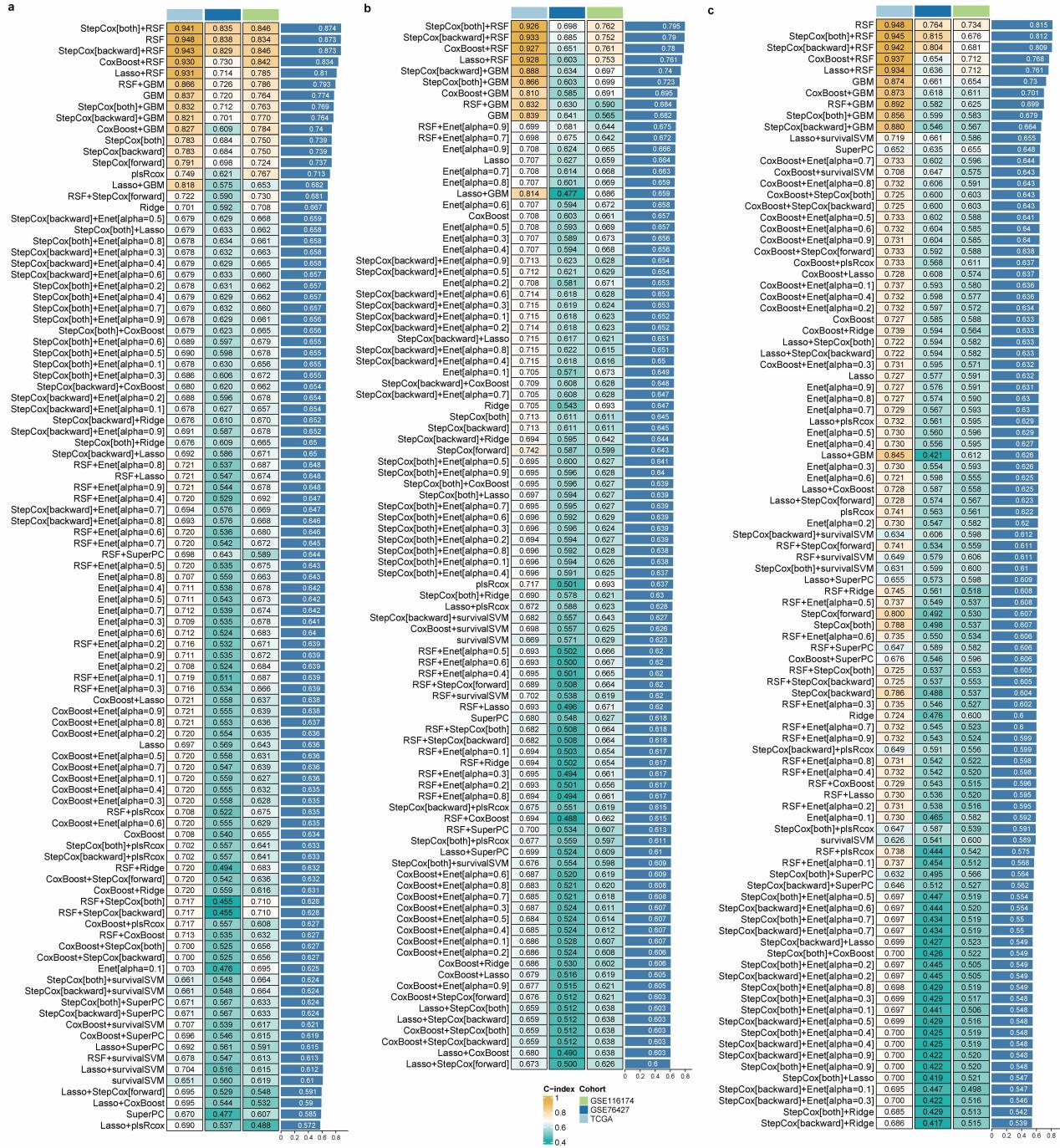

**Supplementary Fig. 29. MAASig performance in the hepatocellular carcinoma datasets.** C-index of different machine learning models trained on (a) combined TF expression and activity, and single-modality features including (b) expression and (c) activity. Models with less than five genes were excluded for analysis.

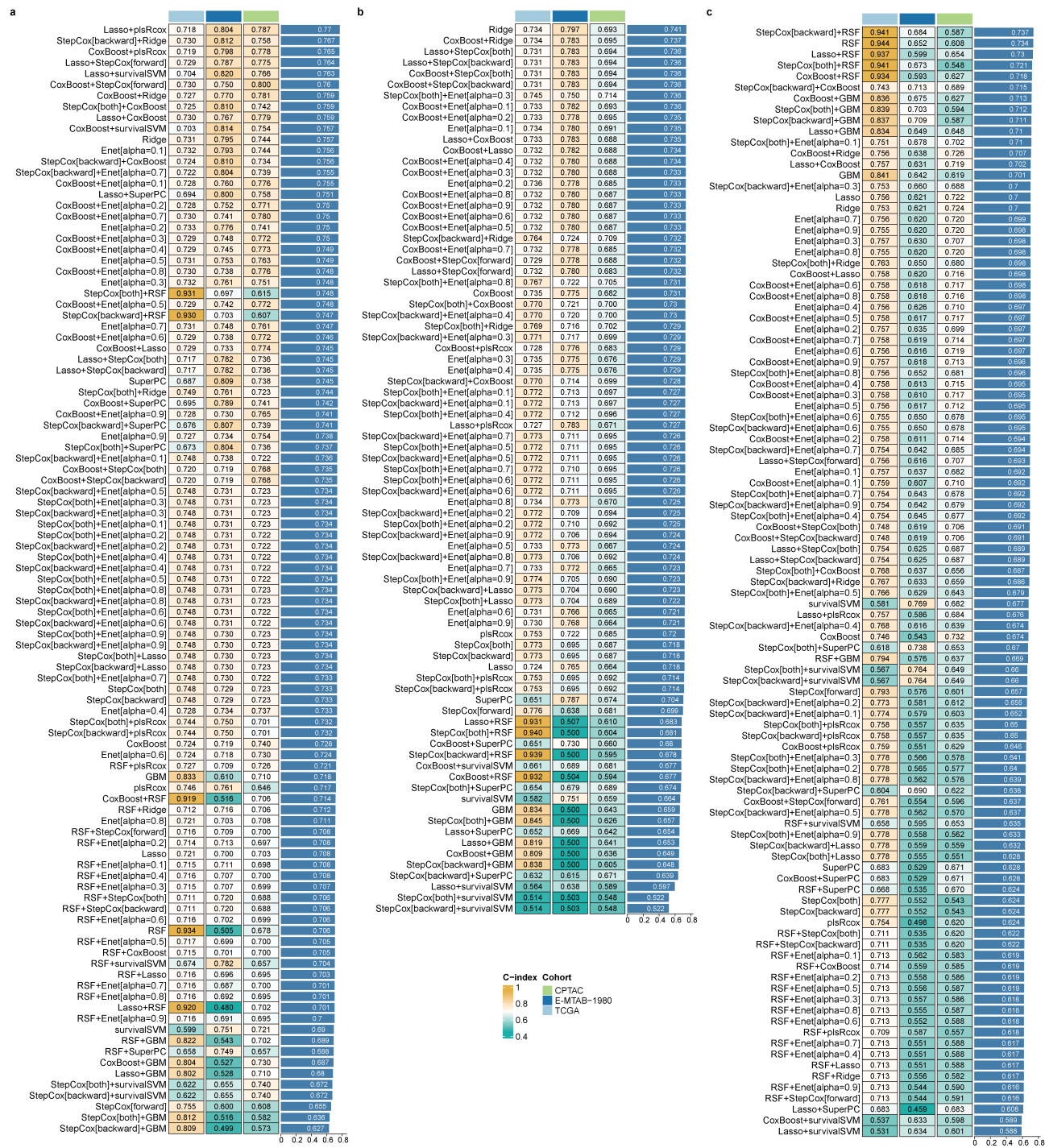

**Supplementary Fig. 30. MAASig performance in the clear cell renal cell carcinoma datasets.** C-index of different machine learning models trained on (a) combined TF expression and activity, and single-modality features including (b) expression and (c) activity. Models with less than five genes were excluded for analysis.

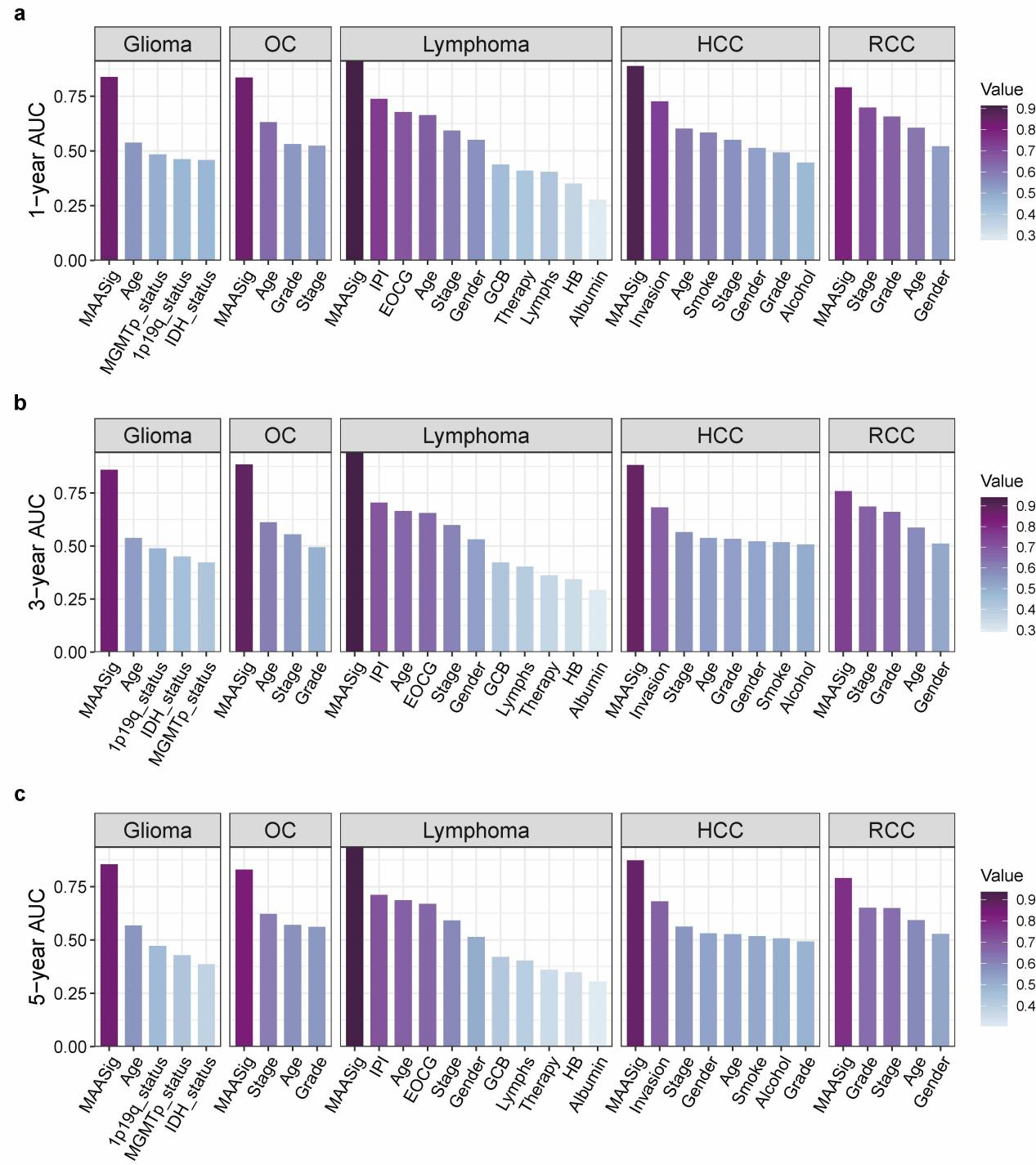

**Supplementary Fig. 31. Prognostic prediction performance.** Average (a) 1-year, (b) 3-year and (c) 5-year AUC of traditional clinical features and MAASig for survival prediction across multiple cancer types, including glioblastoma, ovarian cancer (OC), B-cell lymphoma, hepatocellular carcinoma (HCC), and clear cell renal cell carcinoma (ccRCC).

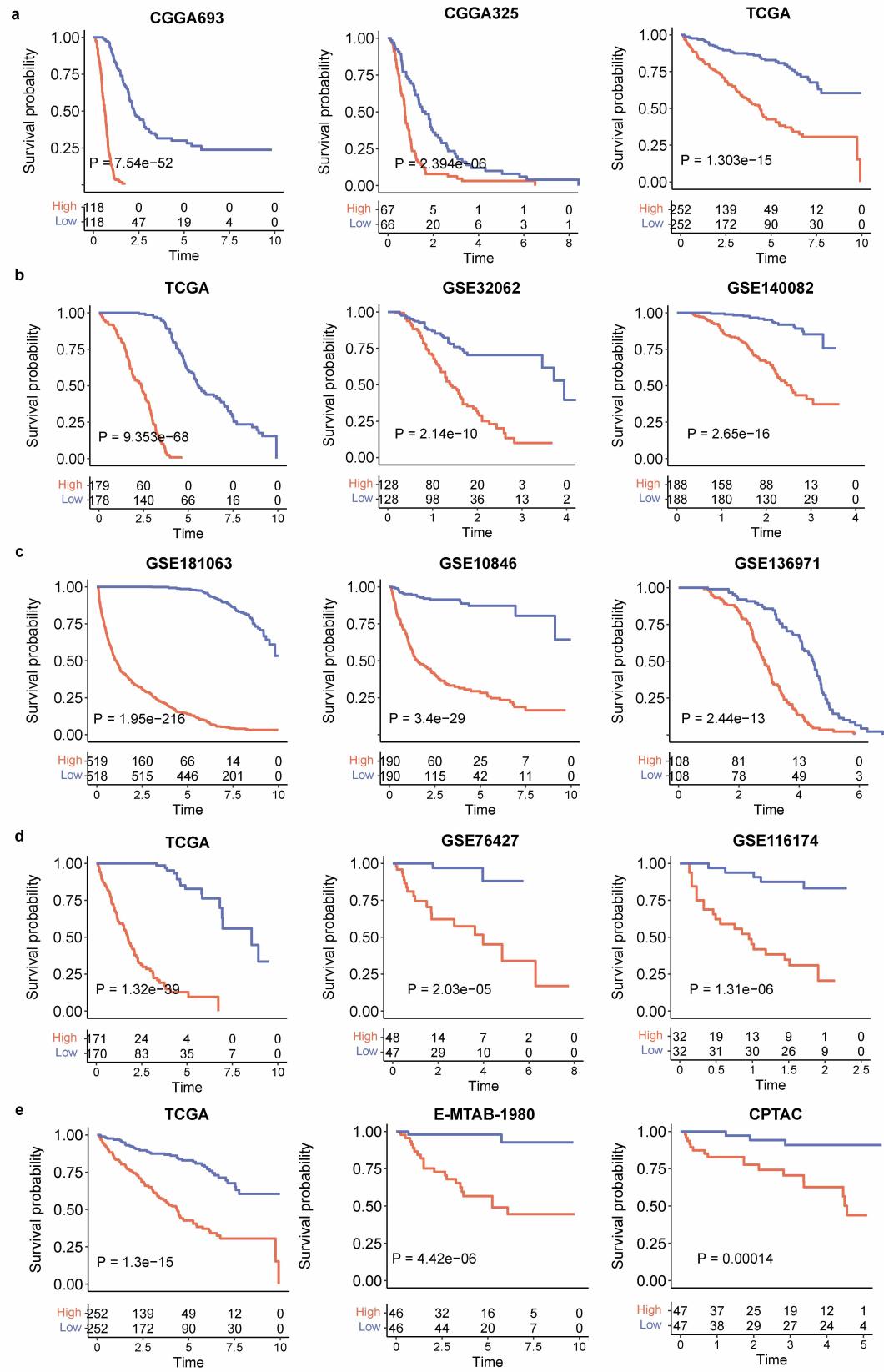

**Supplementary Fig. 34. Kaplan–Meier survival curves demonstrating the clinical relevance of MAASig in different cancers. a.** Glioblastoma. **b.** Ovarian cancer. **c.** B-cell lymphoma. **d.** Hepatocellular carcinoma. **e.** Clear cell renal cell carcinoma. MAASig stratification was determined at the median value. P-values were calculated by the two-tailed log-rank test.

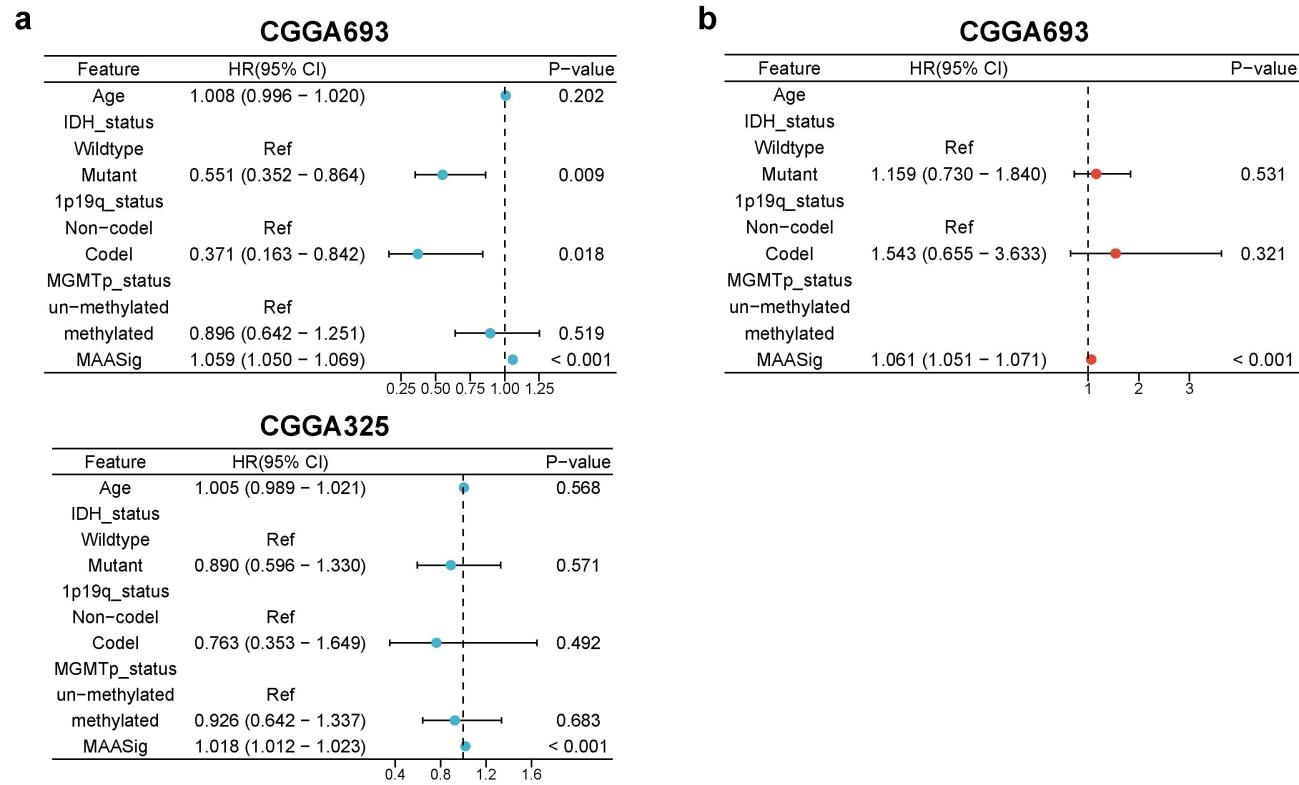

**Supplementary Fig. 33. Cox proportional-hazards model analysis revealed the prognostic value of MAASig in glioblastoma. a.** Univariate Cox model. **b.** Multivariate Cox model. Points represent the hazard ratios, and the horizontal bars extend from the lower limits to the upper limits of the 95% confidence intervals of the estimates of the hazard ratios. P-values were determined by the two-tailed Wald test.

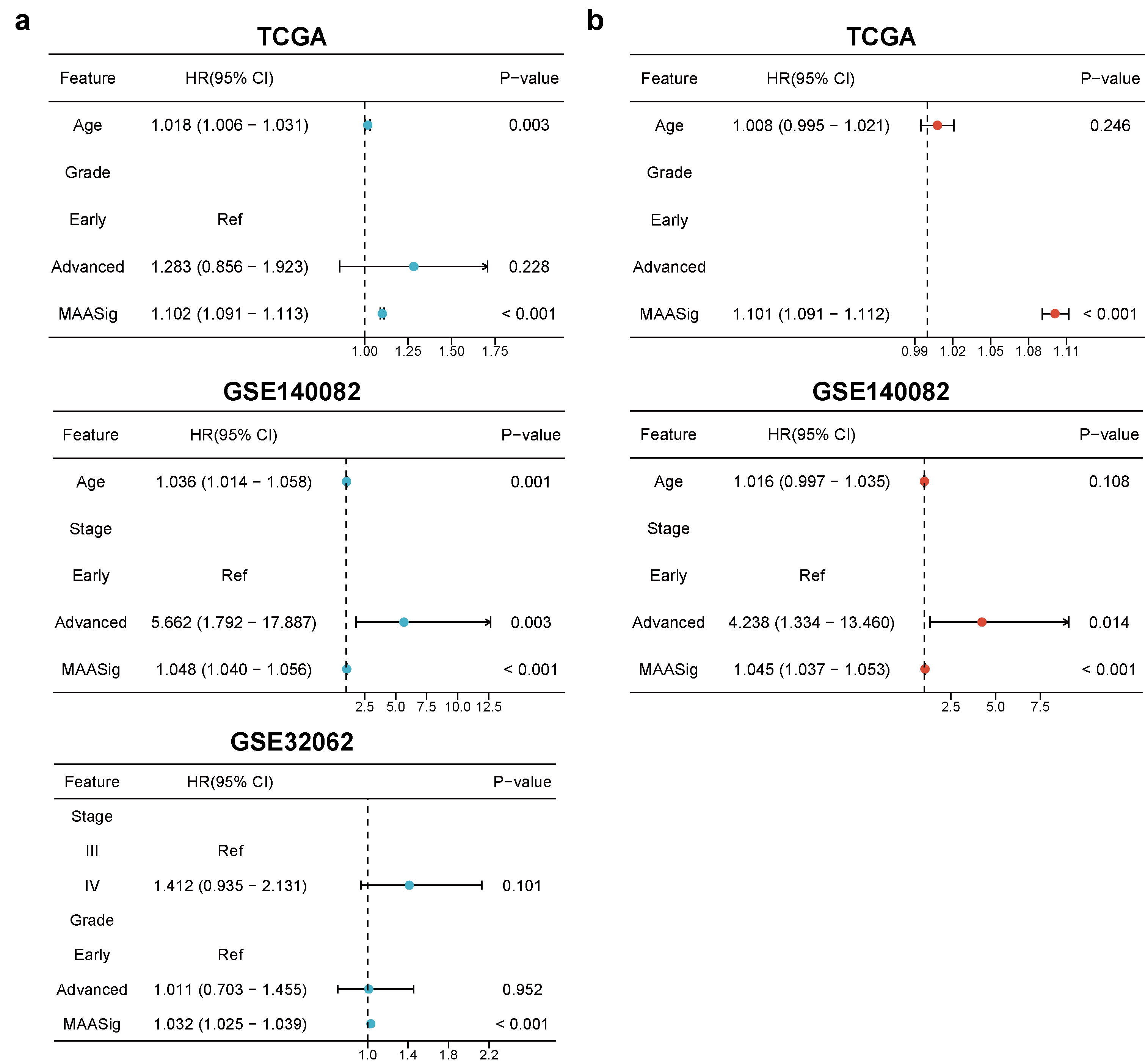

**Supplementary Fig. 34. Cox proportional-hazards model analysis revealed the prognostic value of MAASig in ovarian cancer. a.** Univariate Cox model. **b.** Multivariate Cox model. Points represent the hazard ratios, and the horizontal bars extend from the lower limits to the upper limits of the 95% confidence intervals of the estimates of the hazard ratios. P-values were determined by the two-tailed Wald test.

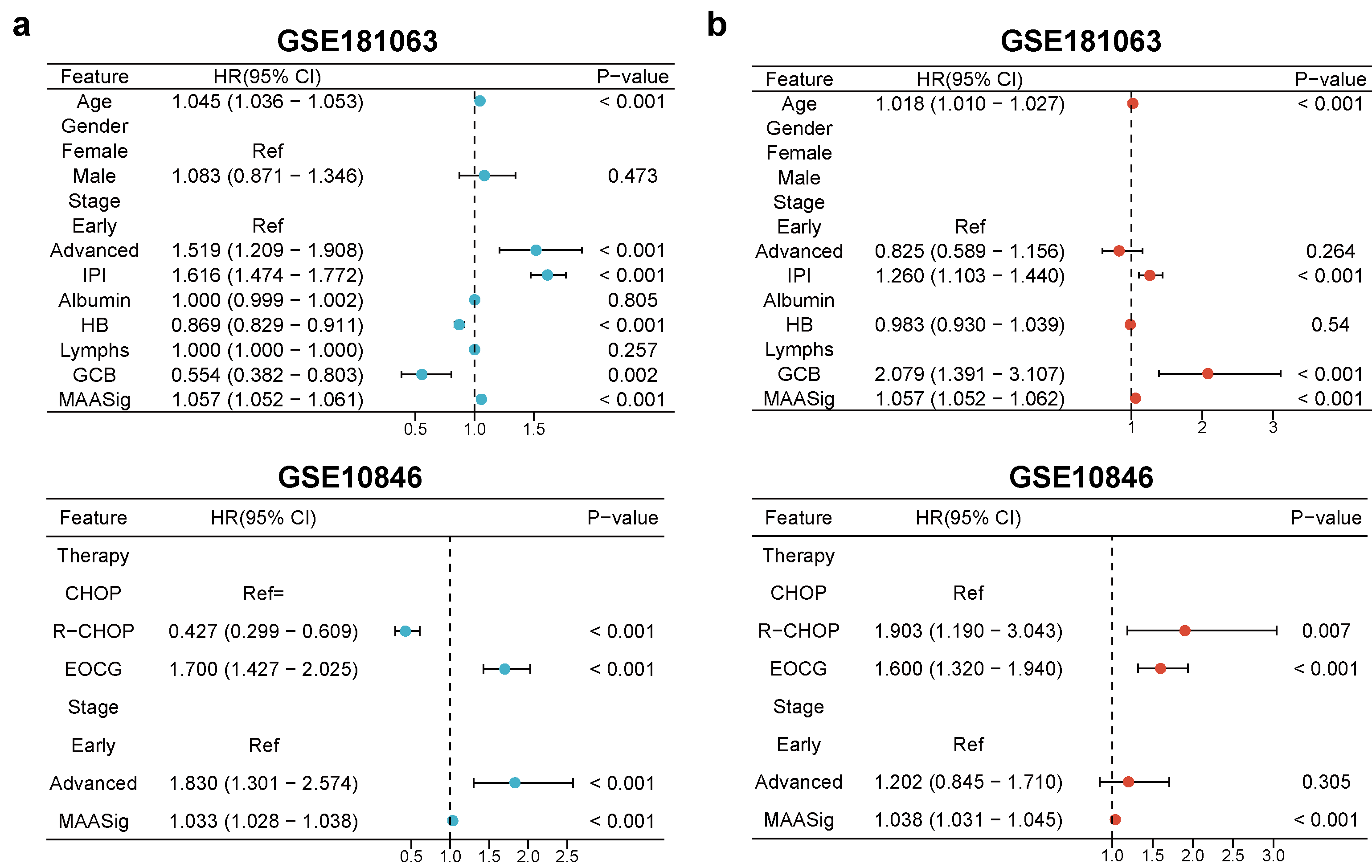

**Supplementary Fig. 35. Cox proportional-hazards model analysis revealed the prognostic value of MAASig in B-cell lymphoma. a.** Univariate Cox model. **b.** Multivariate Cox model. Points represent the hazard ratios, and the horizontal bars extend from the lower limits to the upper limits of the 95% confidence intervals of the estimates of the hazard ratios. P-values were determined by the two-tailed Wald test.

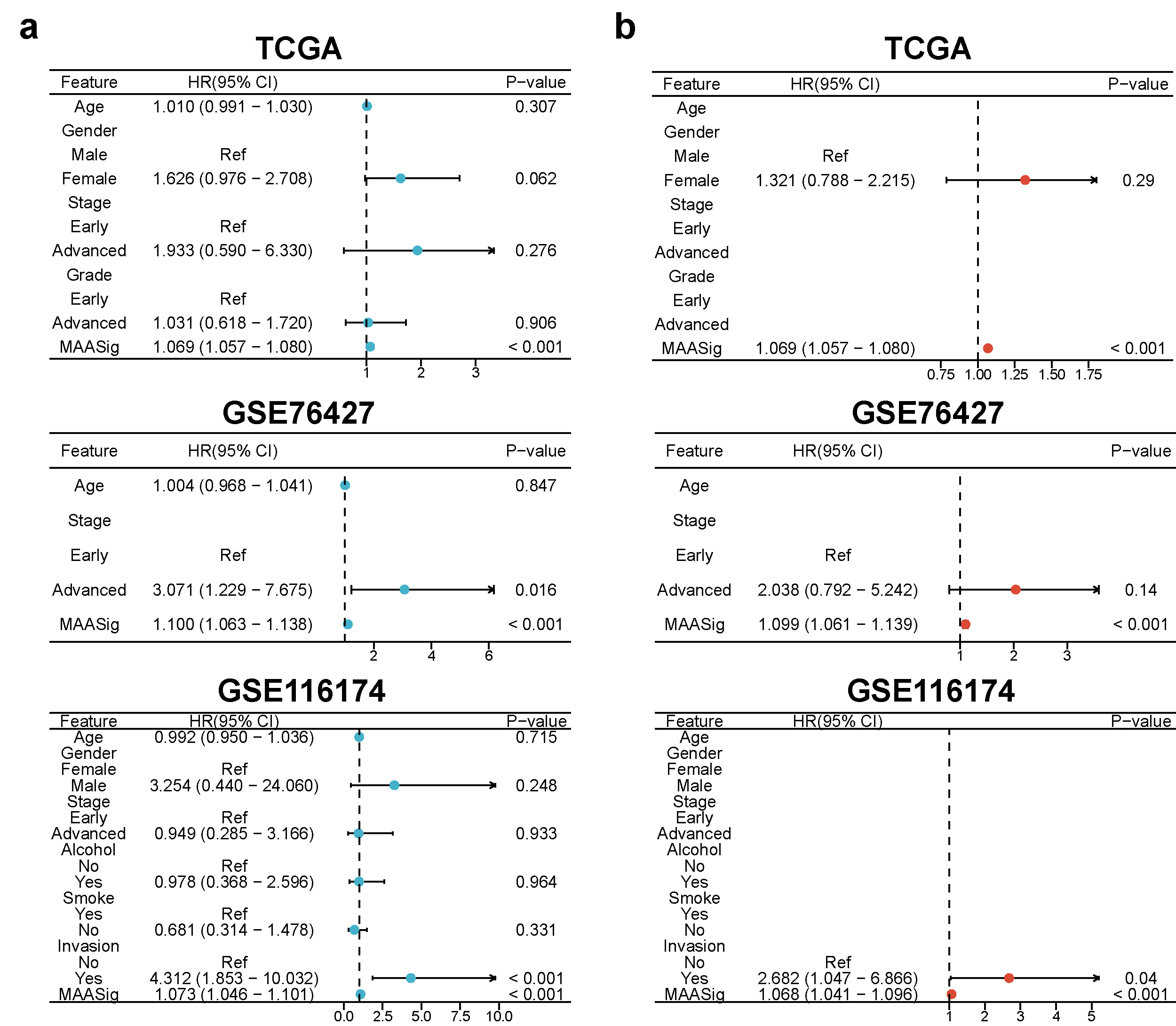

**Supplementary Fig. 36. Cox proportional-hazards model analysis revealed the prognostic value of MAASig in hepatocellular carcinoma. a.** Univariate Cox model. **b.** Multivariate Cox model. Points represent the hazard ratios, and the horizontal bars extend from the lower limits to the upper limits of the 95% confidence intervals of the estimates of the hazard ratios. P-values were determined by the two-tailed Wald test

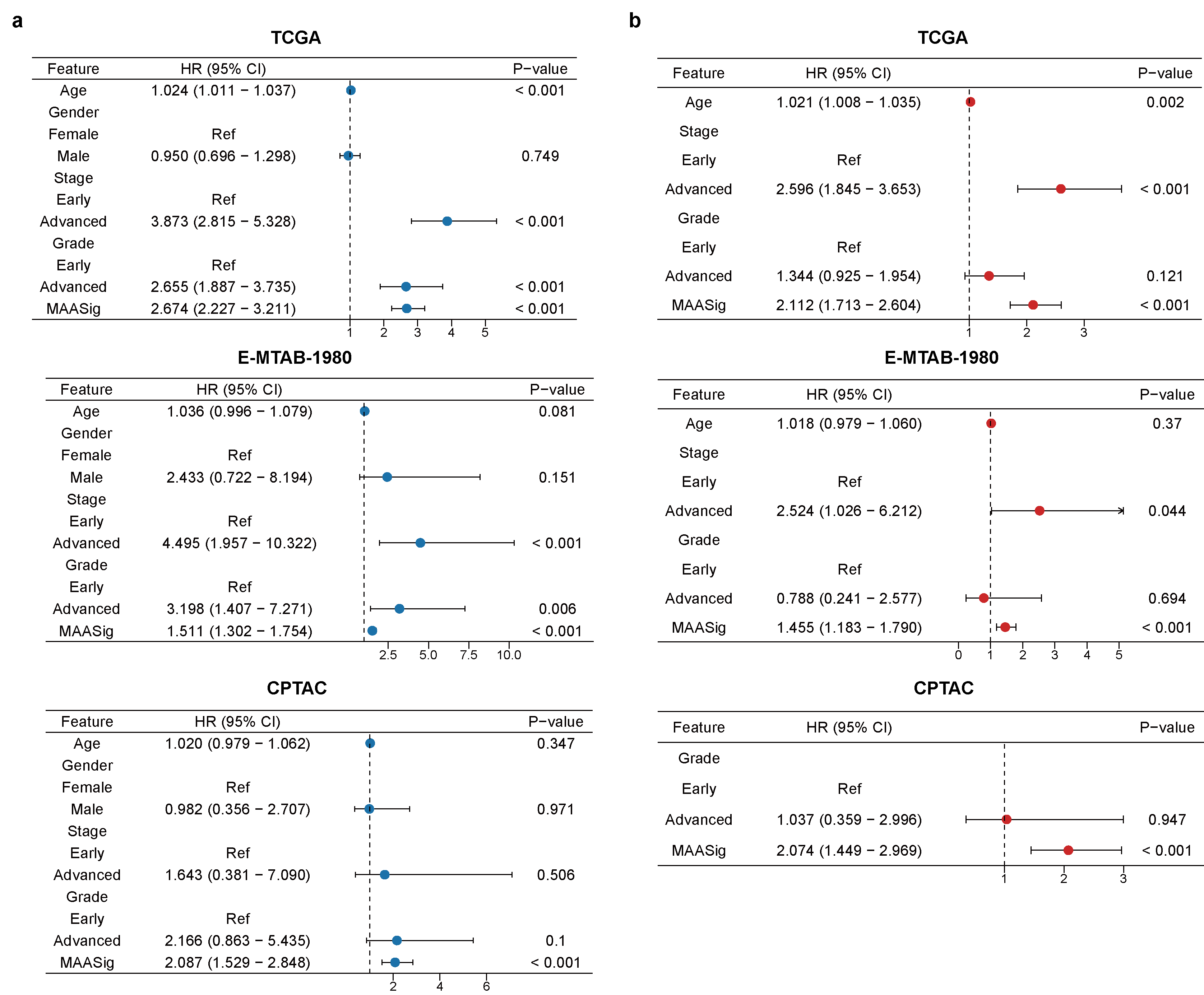

**Supplementary Fig. 37. Cox proportional-hazards model analysis revealed the prognostic value of MAASig in clear cell renal cell carcinoma. a.** Univariate Cox model. **b.** Multivariate Cox model. Points represent the hazard ratios, and the horizontal bars extend from the lower limits to the upper limits of the 95% confidence intervals of the estimates of the hazard ratios. P-values were determined by the two-tailed Wald test.

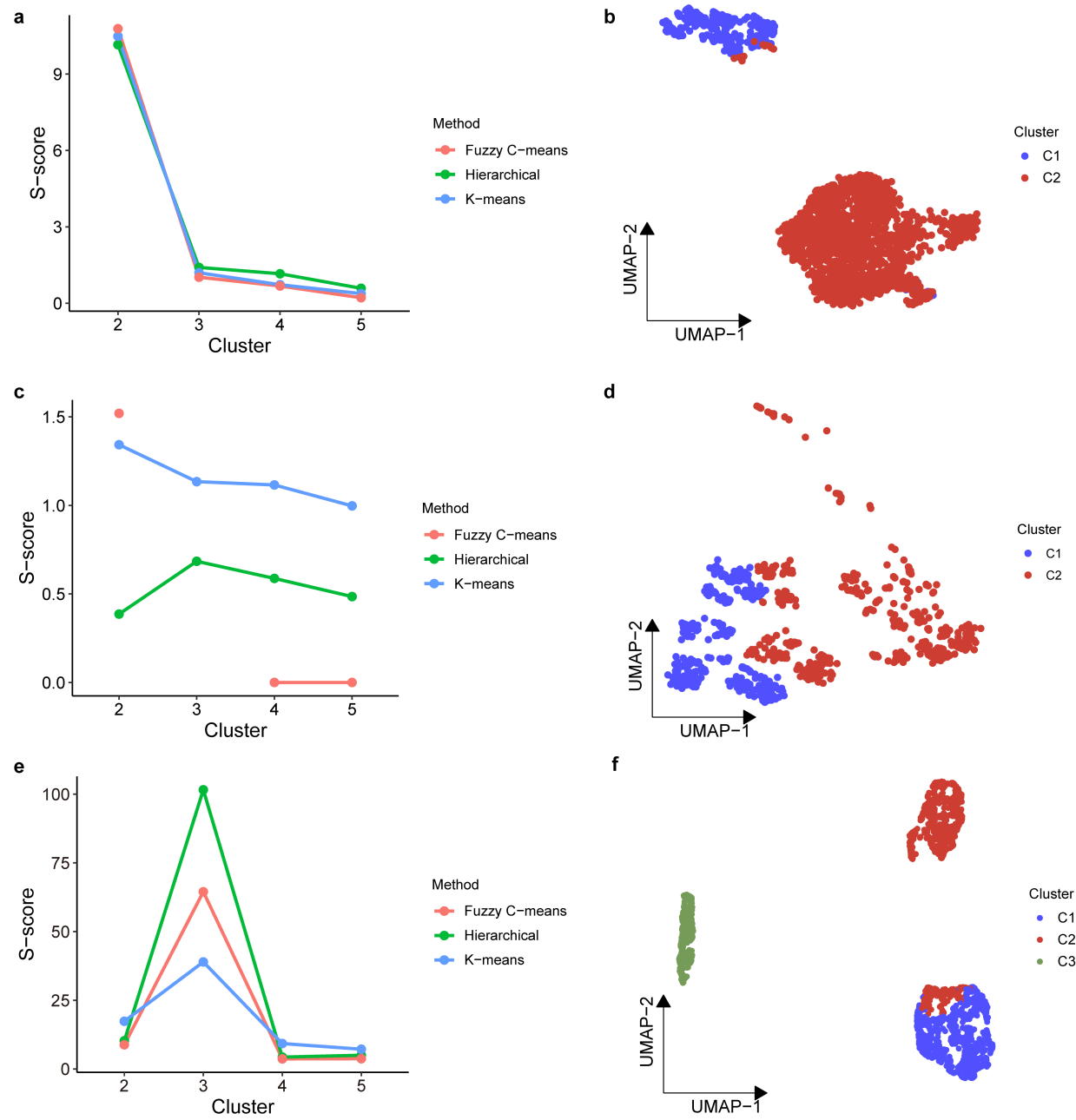

**Supplementary Fig. 38. Tumor cell clusters defined by CNVs and chromatin states of renal cancer. a,b.** Clustering results based on CNVs by epiAneufinder. **c,d.** Clustering results based on CNVs by Copy-scAT. **e,f.** Clustering results based on chromatin accessibility. The (a,c and e) left panel showing the S-score across different clustering numbers. The (b,d and e) right panel showing the UMAP plots of tumor cells substructures.

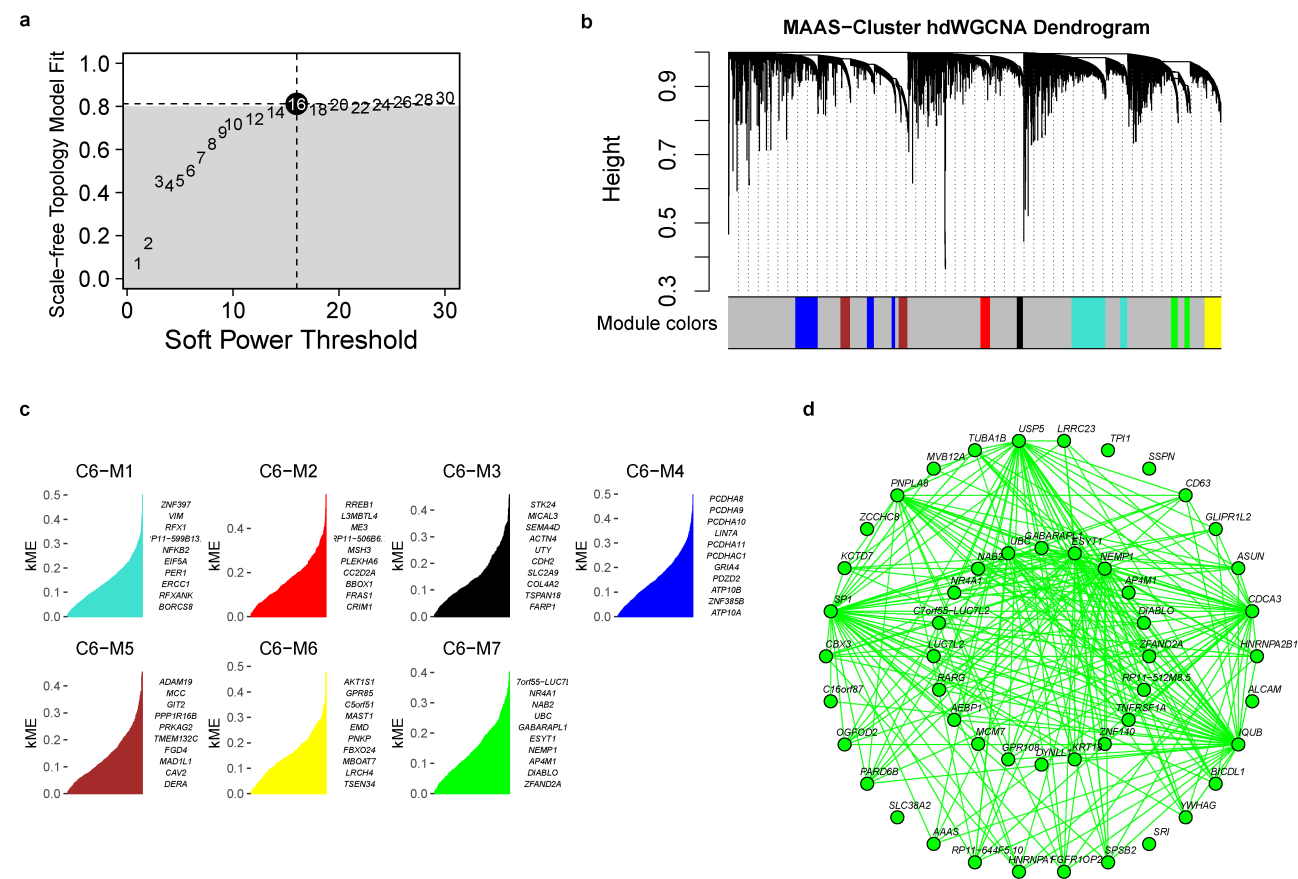

**Supplementary Fig. 39. Identification of gene modules of each MAAS-determined cluster by WGCNA. a.** Determining the soft power threshold of 16 by scale-free topology model fitness. **b.** Dendrogram of gene modules. Grey color indicates genes with unknown module assignment. **c.** Genes with top eigengene-based connectivity (kME) in each module. **d.** Gene network of top-50 genes in module 7 representing MAAS-cluster 6.

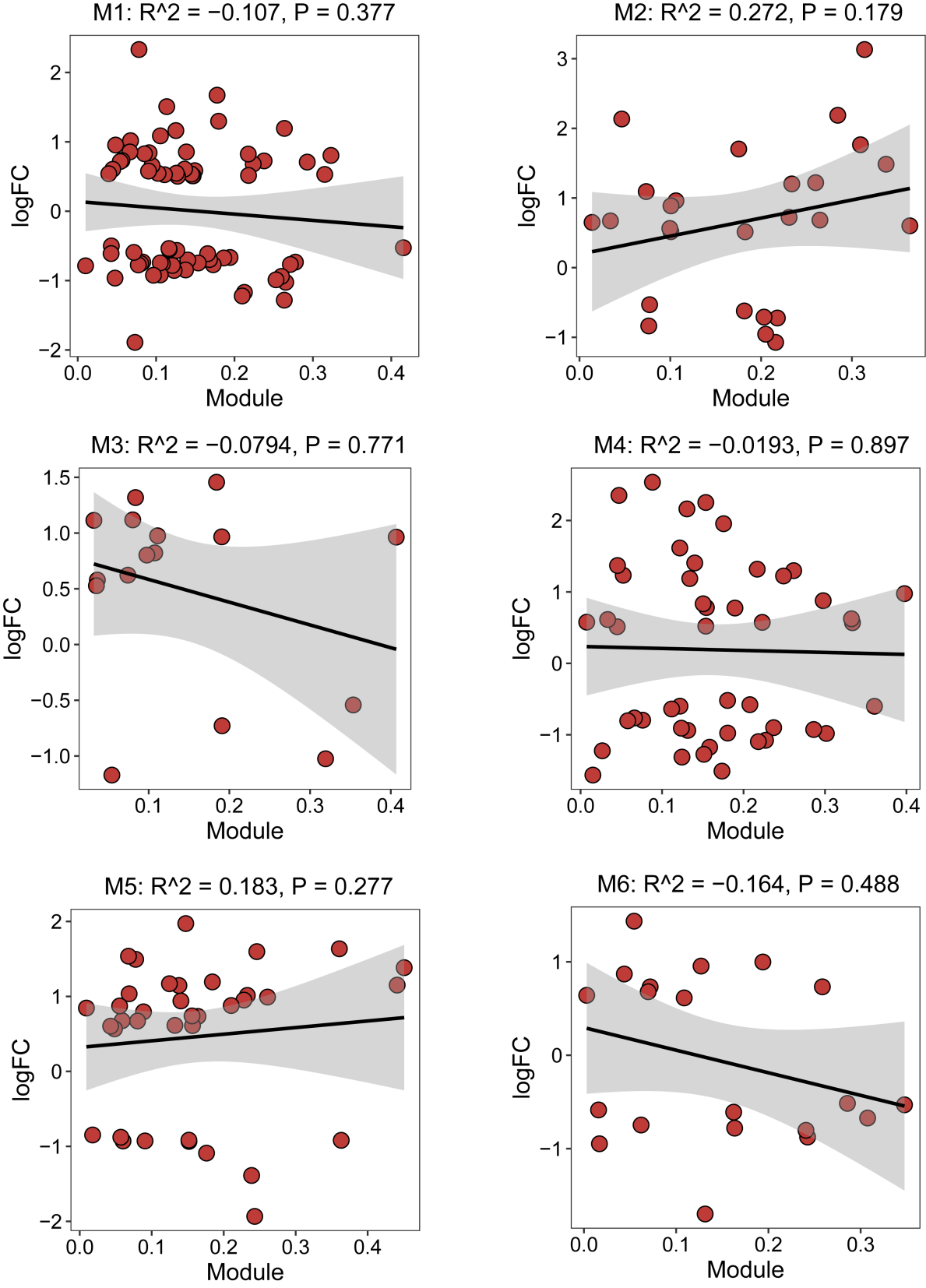

**Supplementary Fig. 40. Spearman correlation between eigengene-based connectivity (kME) of each gene module. The shading represents 95% confidence intervals.**

**Supplementary Table 1. Datasets of MAASig performance estimation.**

| **Cancer type** | **Training set** | **Test set** |
| --- | --- | --- |
| GBM | CGGA693 | CGGA325, TCGA |
| OC | TCGA | GSE140082, GSE32062 |
| B-cell lymphoma | GSE181063 | GSE10846, GSE136971 |
| HCC | TCGA | GSE116174, GSE76427 |
| ccRCC | TCGA | E-MTAB-1980, CPTAC |

**Supplementary Table 2. TFs included in the MAASig of each cancer.**

| **Cancer type** | **Model** | **Signature TFs** | |
| --- | --- | --- | --- |
|  |  | **Expression** | **Activity** |
| GBM | StepCox (both direction) + RSF | *OLIG1*, *ETV6*, *EGR4*, *HOXD11*, *EN1*, *HOXD10* | *EGR2*, *ELF1*, *ELK3*, *EP300*, *ETS1*, *FOXM1*, *KLF5*, *SP3*, *SPI1* |
| OC | RSF | *TFAP2B*, *ZNF148*, *ELK3*, *HEYL*, *ZBTB*, *ZBTB6*, *SREBF1*, *ID3*, *PRDM4* | *ARID4B*, *ELK3*, *ERF*, *NPAS2*, *PURA*, *TP53*, *TWIST1* |
| B-cell lymphoma | RSF | *ELK1*, *TFDP1*, *ELF1*, *ERF*, *ZBTB1*, *KLF16*, *KLF13*, *NFYC*, *TFAP2A*, *ZNF148*, *RREB1*, *CEBPZ*, *ARNTL*, *EPAS1*, *NFYA*, *RELB*, *NAIF1*, *MYPOP*, *E2F8*, *OTX1*, *RELA*, *ZNF423*, *MYBL1* | *ATF6*, *E2F2*, *E2F3*, *EGR2*, *ESR1*, *ETS2*, *ETV2*, *MXI1*, *RFX1*, *SPDEF*, *ZBTB7B*, *ZFX*, *ZNF143* |
| HCC | StepCox (both direction) + RSF | *NRF1*, *ETV4*, *CREB1*, *ATF1*, *HIF1A*, *SP3*, *ZIC5*, *SPZ1*, *GLIS1*, *YBX1*, *TFE3*, *MYCN*, *CREB3L*, *ZKSCAN4*, *IRF5*, *E2F8*, *ZIC2*, *E2F7*, *GRHL1* | *ARNT*, *CTCFL*, *E2F6*, *EGR4*, *ETV3*, *GLI2*, *HIF1A*, *INSM1*, *LYL1*, *MAFA*, *MAZ*, *MXI1*, *MYC*, *SP3*, *TFDP1*, *ZBTB16* |
| ccRCC | Lasso + PLSR Cox | *PURA*, *MYCN*, *IRF6*, *ZIC2*, *OTX1*, *AR*, *BARX1*, *HOXA2*, *IRF7*, *FOXP3*, *RUNX1*, *MSC*, *FOXM1* | *E2F6*, *GLIS3* |

**Supplementary Table 3. Markers of tumor cells.**

| **Cancer type** | **Marker** | **Ref. (PMID)** |
| --- | --- | --- |
| Ovarian cancer | *MUC16*, *WFDC2*, *PAX8*, *EPCAM* | 34739872, 31825847 |
| GBM | *PDGFRA*, *SOX2*, *OLIG1*, *NES*, *S100B*, *EGFR*, chr7p gain/chr10q loss | 35140215, 34644115 |
| PPFE | *IGFBP5*, *VEGFA*, *CD44*, *EGFR*, *MEIS1*, *NR3C1*, *ZIC4* | 35803925 |
| B-cell lymphoma | *MS4A1*, *CD40*, *BANK1*, *PAX5* | 10x Genomics |
| ccRCC | *NDUFA4L2*, *CA9*, *KRT18*, *KRT8*, chr3p loss/chr5q gain | 36607615 |
